# Beyond Spike: Identification of nine highly prevalent SARS-CoV-2-specific CD8 T-cell epitopes in a large Norwegian cohort

**DOI:** 10.1101/2021.10.13.463911

**Authors:** Saskia Meyer, Isaac Blaas, Ravi Chand Bollineni, Marina Delic-Sarac, Trung T. Tran, Cathrine Knetter, Ke-Zheng Dai, Torfinn Støve Madssen, John T. Vaage, Alice Gustavsen, Weiwen Yang, Lise Sofie Haug Nissen-Meyer, Karolos Douvlataniotis, Maarja Laos, Morten Milek Nielsen, Bernd Thiede, Arne Søraas, Fridtjof Lund-Johansen, Even H. Rustad, Johanna Olweus

## Abstract

T-cell epitopes with broad population coverage may form the basis for a new generation of SARS-CoV-2 vaccines. However, published studies on immunoprevalence are limited by small test cohorts, low frequencies of antigen-specific cells and lack of data correlating eluted HLA ligands with T-cell responsiveness. As the protective role of pre-existing cross-reactivity to homologous peptides is unclear, we aimed to identify SARS-CoV-2-specific minimal epitopes recognized by CD8 T-cells among 48 peptides eluted from prevalent HLA alleles, and an additional 84 predicted binders, in a large cohort of convalescents (n=83) and pre-pandemic control samples (n=19). We identified nine conserved SARS-CoV-2-specific epitopes restricted by four of the most prevalent HLA class I alleles in the Norwegian study cohort, to which responding CD8 T cells were detected in 70-100% of convalescents expressing the relevant HLA allele. Only two of these were derived from the Spike protein, included in current vaccines. We found a strong correlation between immunoprevalence and immunodominance. Thus, the CD8 T-cell response to SARS-CoV-2 is more focused than previously believed. Using a new algorithm, we predict that a vaccine including these epitopes could induce a T-cell response in 83% of Caucasians.

## INTRODUCTION

SARS-CoV-2 variants can escape neutralizing antibodies induced by current vaccines^1–8^. Complementary vaccines aimed at evoking cytotoxic T-cell responses towards multiple SARS-CoV-2 proteins may overcome these limitations, and could prove fundamental in protecting immunodeficient patients with no or impaired B-cell responses. Development of such vaccines with broad population coverage will require identification of conserved CD8 T-cell epitopes that are presented on frequent HLA class I alleles, and that are capable of eliciting an immune response in the majority of individuals expressing the relevant HLA allele (i.e. high immunoprevalence).

A broad range of SARS-CoV-2 CD8 T-cell epitopes has been previously reported^9–32^. However, estimates of immunoprevalence vary greatly between studies and are limited by the low number of donors that were typically included per HLA class I allele in each study. A recent meta-analysis showed that the median number was five (inter-quartile range, IQR: 3-8)), with only 1.1% of epitopes (8 of 711) identified in two or more studies with an immunoprevalence >70%^33^. Four out of eight were from ORF1a, which has the highest homology with human common cold coronaviruses (HCoV)^34^, and with the most prevalent reported in 91.3% of convalescents (HLA-A*01:01 ORF1a_1637-1646_; TTDPSFLGRY)^33^. Previous studies of CD4 T-cell responses to SARS-CoV-2 have shown negative^35^ as well as positive^36^ consequences of pre-existing cross-reactivity to HCoV epitopes. Cross-reactive CD8 T cells in healthy unexposed individuals have low avidity and respond poorly to SARS-CoV-2^13^. For vaccine development, it is therefore important to map immunoprevalent CD8 epitopes that are specific for SARS-CoV-2.

In the majority of studies, binding of peptide-HLA (pHLA) multimers to CD8 T cells among peripheral blood mononuclear cells directly after isolation from COVID-19 convalescent individuals was used as readout to identify HLA class I epitopes (*ex vivo* assays)^9, 12–15, 17, 18, 20, 21, 27, 29, 32, 37^. Recent advances in pHLA multimer technology allow the simultaneous detection of hundreds to thousands of antigen-reactive T-cell subsets with high sensitivity^38^. Prediction algorithms are used to select among the wealth of candidate peptides that in theory could bind to each of a multitude of HLA alleles. This can include peptides not naturally processed, and validation of epitopes is therefore important to design vaccines intended to induce T-cell immunity. pHLA multimer positive T cells reactive to SARS-CoV-2 peptides have, however, in the majority of cases not been validated for recognition of endogenously processed antigen to exclude “false” epitopes. Moreover, only one study investigated T-cell responsiveness to eluted HLA ligands, focusing on peptides derived from the M and NSP13 region of ORF1a. Five peptides (M=1, NSP13=4) with 89-100% sequence homology to other corona viruses were identified by mass spectrometry that elicited T-cell responses in healthy individuals, not previously infected with SARS-CoV-2^23^. Thus, a systematic correlation between HLA peptidomics and T-cell responsiveness in SARS-CoV-2 convalescents is currently lacking. Finally, although other studies report estimated population coverage of immunogenic epitopes based on HLA allele frequency, these numbers have not been adjusted for immunoprevalence and may therefore overestimate projected vaccine coverage^33, 39, 40^.

In this study, we identify highly immunoprevalent CD8 T-cell epitopes that are specific for SARS-CoV-2, and endogenously processed and presented on frequently expressed HLA alleles. Using a novel algorithm, we demonstrate the importance of considering immunoprevalence when selecting T-cell epitopes to design vaccines that induce broad population coverage.

## RESULTS

### Broad CD4 and CD8 T-cell responses to SARS-CoV-2 peptide pools among COVID-19 convalescents

We included a large cohort of HLA-typed COVID-19 convalescents (mild to severe disease) and healthy controls (Suppl Table 1) in Norway. Convalescent individuals were included based on a positive SARS-CoV-2 PCR test (n=93), or positive antibody response alone (n=3). Among the 93 individuals from whom serum was available 73 showed antibody responses to the receptor binding domain (RBD) and Nucleocapsid protein of SARS-CoV-2. Fourteen pandemic (PCR negative and antibody negative) (Suppl. Fig 1A) and 30 pre-pandemic control samples were included (Suppl Table 2). The resting state of the T cells across cohorts was confirmed by demonstration of low expression levels of the activation markers CD38/HLA-DR and CD137/CD134 or CD69 on freshly isolated PBMC (Suppl Fig. 1D).

We first investigated functional T-cell responses to pools of overlapping 15-mer peptides covering the structural proteins Spike (S, four pools), Nucleocapsid (N, two pools), Membrane (M, one pool), and Envelope (E, one pool) and the non-structural protein ORF3a (O3a, one pool) of SARS-CoV-2 in convalescent and healthy controls (Suppl Table 2). Upregulation of activation markers on CD4 and CD8 T cells was determined by flow cytometry after a short-term (5 day) culture with peptides to amplify memory responses, and a subsequent peptide restimulation for 20-24 h (Suppl Fig. 2A+B). Strong responses were observed for the majority of the patients (Fig. 1A+B). The average frequencies of responding T cells were higher among CD4 than CD8 T cells, possibly because 15-mers require (more) trimming to fit into the HLA class I binding groove as compared to HLA class II. The highest individual responses were, however, found among CD8 T cells. The responses were significantly higher in convalescents than in controls for all pools except the E pool (E_1-75_) for both CD4 and CD8 T cells, and the M pool covering the N-terminal part (M_1-115_) for CD8 (Fig. 1A). CD4 and CD8 T-cell responses generally correlated well (Fig. 1C). These results are consistent with findings in previous studies^27, 37, 41–46^.

**Fig. 1.**
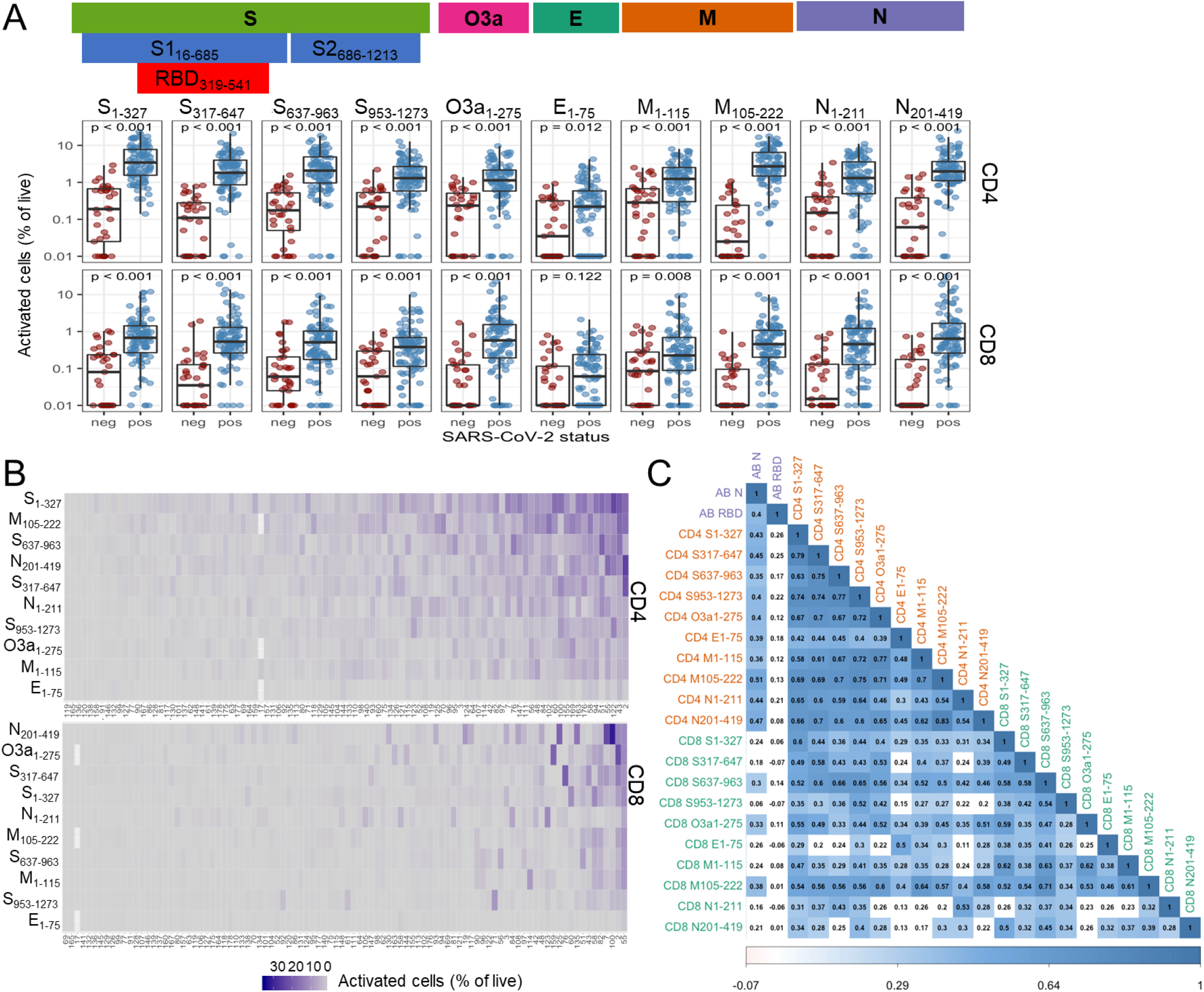
Functional CD4 and CD8 T-cell responses against SARS-CoV-2 peptide pools. (A) Functional CD4 and CD8 T-cell responses against the structural SARS-CoV-2 proteins Spike (S), Envelope (E), Membrane (M) and Nucleocapsid (N) and the non-structural protein ORF3a (O3a) in COVID-19 convalescent (n=96; SARS-CoV-2 pos) and healthy control samples (n=33, including 14 pandemic (SARS-CoV-2 neg) and 19 pre-pandemic) assessed after PBMC stimulation with peptide pools (overlapping 15-mers) measured by expression of activation markers CD134^+^CD137^+^ and CD69^+^CD137^+^ on live CD4 and CD8 T cells, respectively (schematic outline of assay setup and gating strategy in Suppl Fig. 2A+B). Wilcoxon test was used to compare response levels between groups and a significant difference was observed for most pools (p < 0.001), as indicated. (B) Heatmap of individual CD4 and CD8 T-cell responses for each COVID-19 convalescent (n=96). Convalescents are sorted from lowest to highest overall response (columns: sum of response to all antigens). Antigens are sorted based on overall responses in patients (rows). (C) Pairwise associations between antibody responses and functional CD4 and CD8 T-cell response to peptide pools (Spearman correlation). Tiles were colored by the magnitude of response where the correlation was statistically significant (FDR<0.01); for tiles with white background the correlation was not statistically significant.

### Identification of a wide repertoire of SARS-CoV-2-specific HLA ligands by mass spectrometry

We next aimed to determine the exact epitopes to which CD8 T cells responded. To this end, we utilized mass spectrometry to analyze the HLA ligandome of 25 mono-allelic B721.221 cell lines overexpressing the SARS-CoV-2 structural proteins S (3 truncated proteins), M, E, or N and five of the most prevalent HLA class I alleles in the Norwegian (Caucasian) population, including HLA-A*01:01, HLA-A*02:01, HLA-A*03:01, HLA-B*07:02 and HLA-B*08:01 (Suppl Table 3), with a cumulative allele frequency of 86.9%. Two additional cell lines expressing the non-structural protein ORF3a and HLA-A*01:01 or HLA-A*02:01 were analyzed. In total, we identified 50 pHLA-combinations across the selected proteins (Suppl Table 4+5), of which 46 were not reported previously in data sets of eluted HLA ligands^12, 47^. SARS-CoV-2 peptides identified in the discovery approach were further validated using synthetic peptide analogs (Methods, and Suppl File 1). We next complemented this list with additional 9- and 10- mer peptides predicted to bind with high affinity (NetMHCpan4.1 BA_Rank<0.2% and NetMHC4.0 Rank<0.25%) to the selected HLA alleles as well as one identified weak binder from literature^37^ (Suppl Table 5), yielding a total of 137 pHLA-combinations (132 peptides) (Fig. 2A) for functional experiments.

**Fig. 2.**
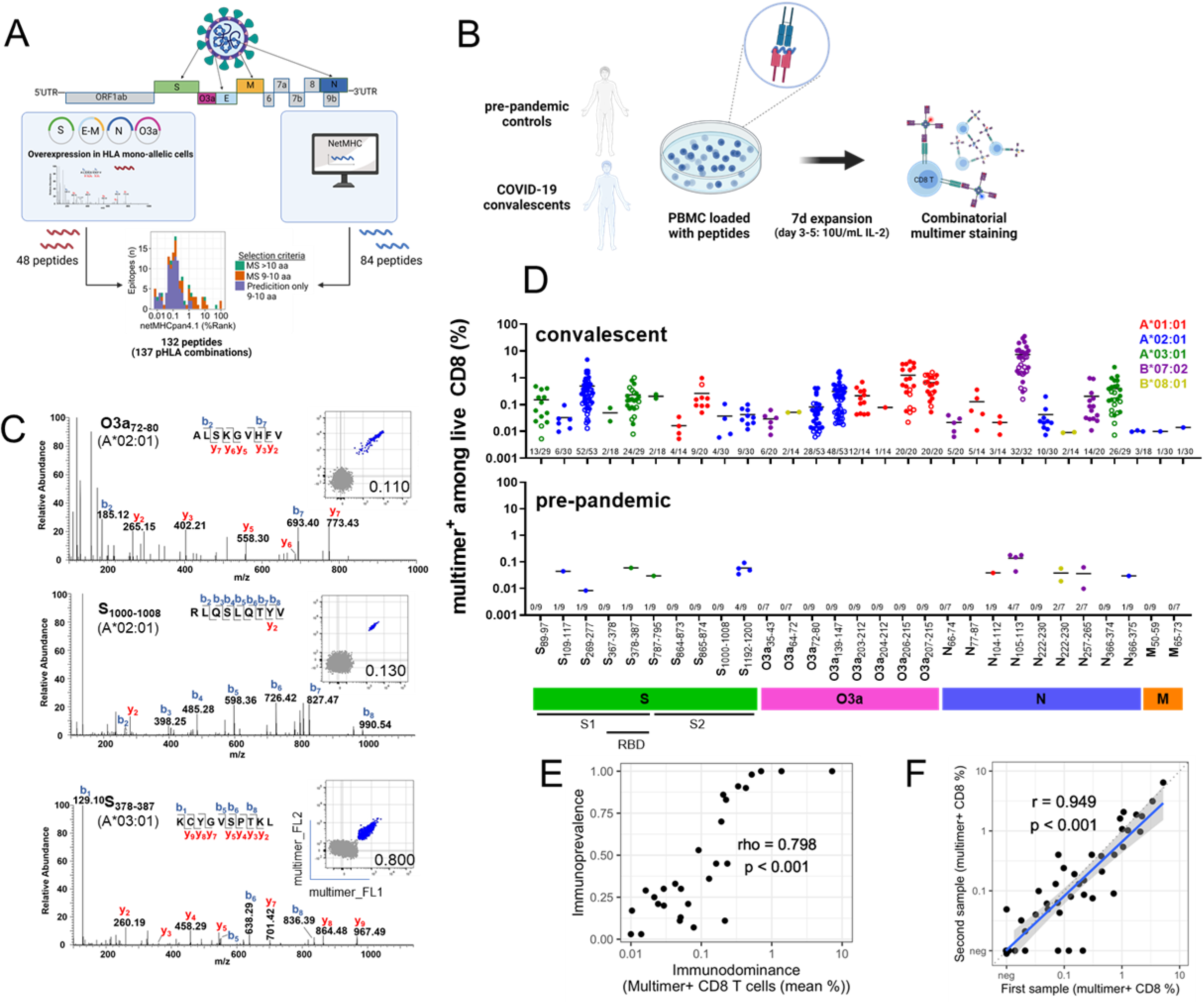
Identification of immunoprevalent and immunodominant CD8 epitopes by screening of directly identified and predicted HLA ligands. (A) Schematic outline for peptide selection. 48 HLA class I-restricted peptides from SARS-CoV-2 proteins expressed in mono-allelic B721.221 cells were identified by mass-spectrometry (MS), an additional 83 by prediction (NetMHCpan4.1 (BA-Rank: <0.2%); NetMHC4.0 (Rank <0.25%)), and one from literature, yielding a total of 132 peptides (137 pHLA-combinations) (overview in Suppl Table 5). (B) Schematic outline of multimer staining assay for the identification of CD8 epitopes. PBMC of pre-pandemic (n=19) and COVID-19 convalescent (n=83) samples were loaded with peptides (n=132; 100ng/mL per peptide), followed by a 7-day expansion. Immunogenic peptides were identified by combinatorial multimer staining (gating strategy in Suppl Fig. 3). (C) Epitopes induce responses of different magnitudes in COVID-19 convalescents after *in vitro* stimulation. Representative flow plots showing multimer staining for three peptides identified by MS, confirmed to be immunogenic. (D) Magnitude of CD8 T-cell responses to the 29 immunogenic SARS-CoV-2 derived peptides, as determined by multimer staining in convalescent and pre-pandemic samples (individual data points with mean). • Cohort 1 (50 convalescents + 19 pre-pandemic samples analyzed); ○ Cohort 2 (33 convalescents). For each peptide the # of responses identified among the # of individuals tested is displayed above the x-axis. Color code: HLA-restriction. (E) Correlation between the mean size of multimer positive populations across individuals with a response (i.e. immunodominance, x-axis) and the proportion of convalescent individuals who showed an immune response (i.e. immunoprevalence; y-axis) for each epitope where a specific CD8 T-cell response was found by multimer staining (Spearman correlation, rho=0.798, p<0.001). (F) Multimer responses at different time points from a single individual (Pearson correlation, r=0.949, p<0.001). Each point represents a sample pair from one individual (7 convalescents; on average 56 [38-81] days between sampling). X-axis: first measurement, and y-axis: second measurement.

### Highly immunoprevalent and immunodominant SARS-CoV-2-specific epitopes can be identified following short-term *in vitro* expansion

To identify the peptides with the highest immunoprevalence among the candidates, we initially included 50 out of 84 of the convalescents expressing at least one of the five selected prevalent HLA alleles (range 14-30 convalescents per allele) for further testing. To evaluate the pre-existence of responding memory T cells we included 7-9 pre-pandemic samples per HLA allele (total of 19 pre-pandemic samples from different donors). To identify responding CD8 T cells with high sensitivity, we expanded memory responses during a short-term *in vitro* culture for 7 days in presence of peptides, prior to labelling with pHLA multimers (Fig. 2B). An individual was classified as responder to a peptide if the multimer population 1) contained at least 5 clearly double-positive events (each multimer conjugated to two different fluorochromes), 2) constituted ≥0.005% of the live CD8, and 3) formed a tight cluster, similar to previously defined parameters^48^. Using these strict criteria, we identified 29 CD8 T-cell epitopes, which gave a response in a median of 30% of convalescents per epitope (range 3-100%) (Fig. 2C+D; Suppl Fig. 4A). Among pre-pandemic samples, we found responses to 10 of these epitopes (Fig. 2D, Table 1 and Suppl Fig. 4B), and none to peptides that did not induce a response in convalescents (Suppl Table 6). For six of these a response was found in only one donor (11%), whereas 29-57% of donors responded to the remaining four epitopes. All 29 immunogenic peptides were specific for SARS-CoV-2. When calculating the mean homology across the four most frequent HCoV, the 10 epitopes that induced a response in pre-pandemic samples were, however, significantly more homologous to HCoV than the 19 that did not (44%, IQR: 26-63%, vs 15%, IQR: 7-33%, p=0.02, Wilcoxon test). Higher frequencies of CD8 T cells responsive to SARS-CoV-2 peptide pools have recently been found in tonsil as compared with peripheral blood in pre-pandemic samples from individuals exposed to HCoV^49^.

**Table 1:**
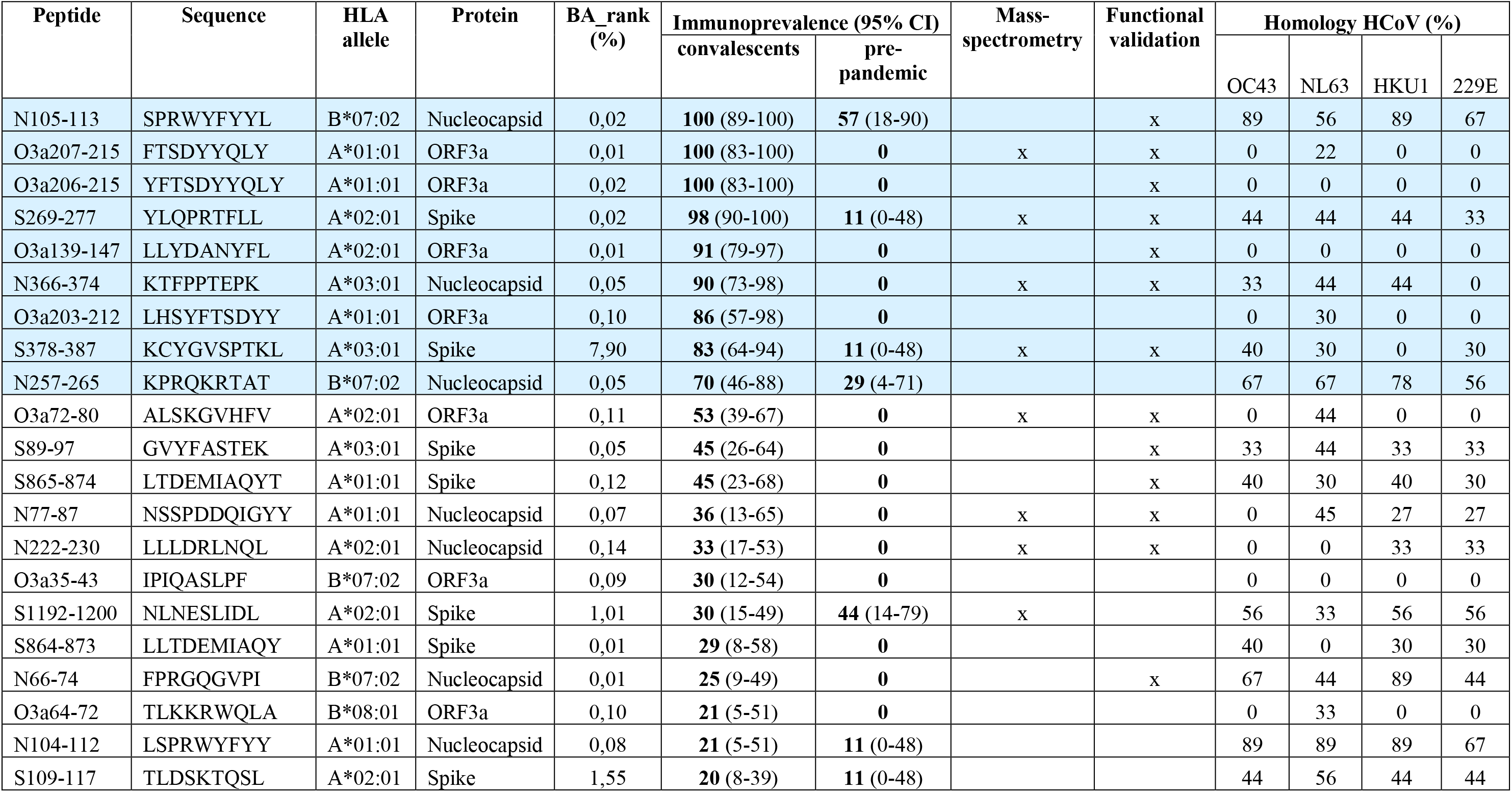
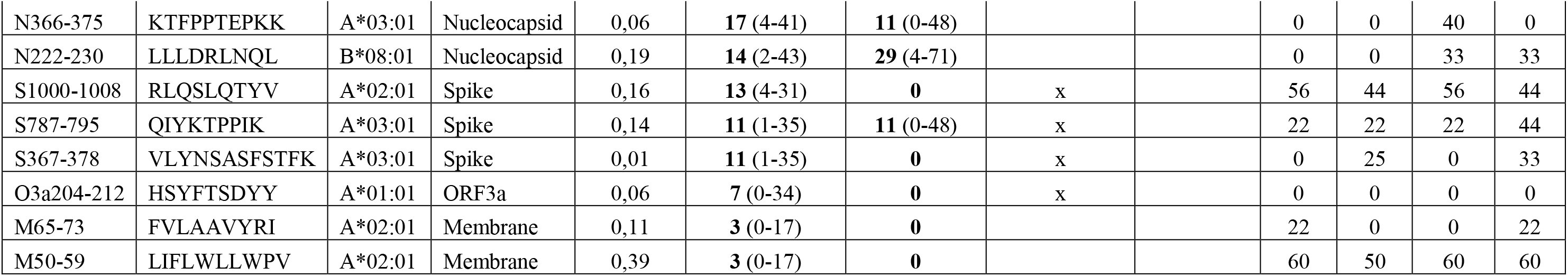
Immunogenic SARS-CoV-2 peptides (n=29) identified by multimer staining in convalescent and pre-pandemic samples sorted from highest to lowest immunoprevalence in convalescents expressing the relevant HLA allele. Peptides with immunoprevalence >70% are shaded in blue. Functional validation = T-cell lines functionally tested. Homology: % sequence identity with human common cold coronaviruses (HCoV).

However, we observed one peptide with 89% homology to another coronavirus which did not induce any response in the pre-pandemic samples, demonstrating that high homology alone is not sufficient for cross-recognition (Table 1).

Among the epitopes identified in this first cohort, four were recognized by 100% of the tested individuals (S1 (1), N (1) and O3a (2)), and an additional four (N, S and O3a) were recognized by 80% or more (Fig. 2D, closed symbols, and Suppl Table 6). To further improve the confidence of our prevalence estimates, we determined CD8 T-cell responses to 10 epitopes with estimated immunoprevalence >50% in a second cohort consisting of 33 convalescent donors not included in the first cohort (Fig. 2D, open symbols). For nine epitopes the estimated immunoprevalence was very similar, while one epitope was found less frequently in the second cohort (Fisher’s exact test p < 0.05) (Suppl Fig. 5A+B). Combining all data, nine epitopes showed an immunoprevalence of 70% or higher (Table 1 and Suppl Table 7). Six of these epitopes were immunogenic in at least 90% of individuals. Two highly immunoprevalent epitopes (>80%) were novel in this study, with no data previously reported in the Immune Epitope Database (IEDB): S_378-387_ (HLA-A*03:01), a 10-mer with poor predicted HLA binding that was identified by MS; and O3a_203-212_ (HLA-A*01:01). Strikingly, there was a strong correlation between immunoprevalence (frequency of responding donors) and immunodominance (size of the multimer positive population in responding individuals) among the 29 immunogenic epitopes (Spearman’s rho=0.798, p<0.001) (Fig. 2E), suggesting preferential processing and presentation of selected peptides across the different SARS-CoV-2 proteins. Demonstrating the reproducibility of our multimer assay, we found a strong correlation between the magnitudes of response in samples obtained at different time points from the same individual (r=0.949, p<0.001) (Fig. 2F). These data also indicate that a single sample accurately represents the immune status of a given individual.

### Identified immunoprevalent epitopes are naturally presented and recognized by T cells

We next validated that T cells responding to the identified epitopes did indeed recognize naturally processed and presented antigens. To this end, multimer positive T cells from convalescent individuals and one pre-pandemic sample were sorted for expansion and generation of T-cell lines. Activation of expanded T-cell lines was measured by flow cytometry after coculturing them with mono-allelic B721.221 cells transduced, or not, to endogenously express the relevant SARS-CoV-2 protein, using peptide-loaded target cells as a positive control. All of the 13 tested epitopes, derived from different SARS-CoV-2 proteins and including 7 of the most immunoprevalent epitopes, were validated (Fig. 3A and Table 1). Five T-cell lines were also tested for cytotoxic ability (five peptides), confirming efficient killing (Fig. 3B). For two epitopes, we were unable to successfully expand T-cell lines from sorted cells. A T-cell line recognizing the 9-mer (FTSDYYQLY) cross-reacted with the corresponding 10-mer (YFTSDYYQLY), and vice versa (Fig. 3C), an observation made also for additional T-cell lines generated from several convalescent individuals, reactive to the same two peptides (Suppl Fig. 6B). The sequences of all but one (N_77-87_; immunoprevalence 36%) of the identified epitopes are conserved in the as of today reported SARS-CoV-2 variants of concern (Suppl Fig. 6B; Suppl Table 7).

**Fig. 3.**
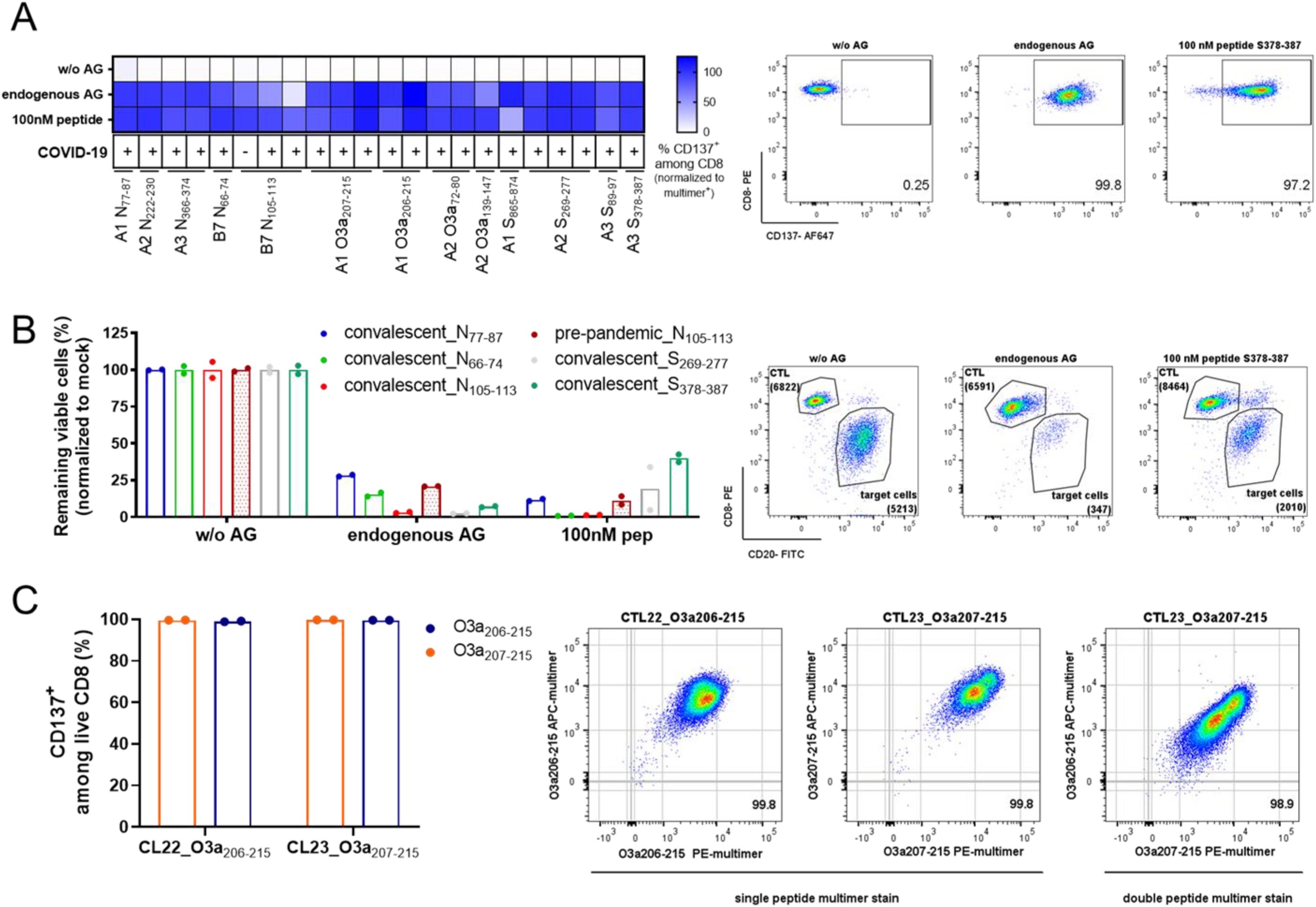
Functional validation of CD8 epitopes. (A+B) Functional validation of immunogenic peptides by co-culture of multimer positive T-cell lines derived from 11 convalescent and one pre-pandemic samples (indicated by COVID-19 status as + or −, respectively) for 20 h with mono-allelic B721.221 cells transduced to express the relevant protein (endogenous antigen (AG)), or not (w/o AG), or loaded with 100 nM peptide as a positive control (n=1 with duplicates or triplicates). (A) Activation of T cells was determined as % CD137^+^ cells among live CD8. The % response is normalized to % multimer^+^ live CD8 T cells in each T-cell line. Representative flow plots are shown for the MS identified peptide S_378-387_ to the right. (B) Target cell killing mediated by multimer^+^ T-cell lines. Values are normalized to matching mock control and displayed as % remaining viable cells. Representative flow plots for the MS identified peptide S_378-387_ to the right show CD20^+^ B721.221 target cells and CD8^+^ CTLs (numbers represent absolute counts). (C) Cross-recognition of T-cell lines sorted from one convalescent for reactivity to ORF3a-derived peptides O3a_207-215_ (FTSDYYQLY; 9-mer) or O3a_206-215_ (YFTSDYYQLY ; 10-mer) to the other peptide. CL22 (sorted for reactivity to O3a_206-215_) and CL23 (sorted for reactivity to O3a_207-215_) recognize the 9- and 10- mer equally well (n=1 with duplicates or triplicates). (Left) Activation marker expression (CD137) after 20 h co-culture with peptide-loaded (100 nM) mono-allelic B721.221 cells. (Right) Flow plots 1+2: dual-color multimer staining of CL22 and CL23 with multimers complexed with relevant peptide. Flow plot 3: multimer staining of CL23 with multimers complexed with 9-mer (x-axis) and 10-mer (y-axis).

### CD8 T-cell immunity to SARS-CoV-2 is more focused than previously assumed

To place our results in context of previous studies, we pooled publicly available data from IEDB covering 53 (39%) of the 137 pHLA-combinations tested in this study. Our analysis of data from IEDB corresponded well with a recent review and meta-analysis of CD8 T-cell responses to SARS-CoV-2 epitopes^33^. Overall, CD8 T-cell responses in our study were directed toward a less diverse set of epitopes and dominated by a few with high immunoprevalence (Fig. 4A). One potential explanation for this was the use of T-cell expansion prior to staining with HLA multimers in this study. Indeed, when comparing our results specifically with other studies employing direct *ex vivo* HLA multimer staining, there was a strong tendency towards higher immunoprevalence in our study (median 45 vs 20%, Wilcoxon signed rank test p = 0.003) (Fig. 4B). Moreover, there was no evidence that our assay resulted in unspecific responses, as *in vitro* validation experiments confirmed T-cell activation and target cell killing for all tested epitopes (Fig. 3A+B), as well as high reproducibility (Fig. 2F). Epitopes identified in only 1-2 convalescents may be of concern with regard to specificity if not validated, but such rare epitopes constituted only 20.7% (n=6) in our study, of which 3 were validated by MS, as compared with 46.7% in IEDB (Fisher’s exact test p=0.04). Although there was a clear trend for the same epitopes to be identified across multiple studies, estimates of immunoprevalence varied greatly. Each estimate was generally uncertain, as reflected by wide 95% confidence intervals, due to the low number of donors tested in most studies (Fig. 4C). In contrast, we could demonstrate that immunoprevalence was significantly higher than 70% (p <0.05) for 7/9 epitopes. Taken together, the results of our study indicate that CD8 T-cell immunity to SARS-CoV-2 is more focused than previously assumed, dominated by a limited set of epitopes that elicit strong immune responses in virtually every infected individual.

**Fig. 4.**
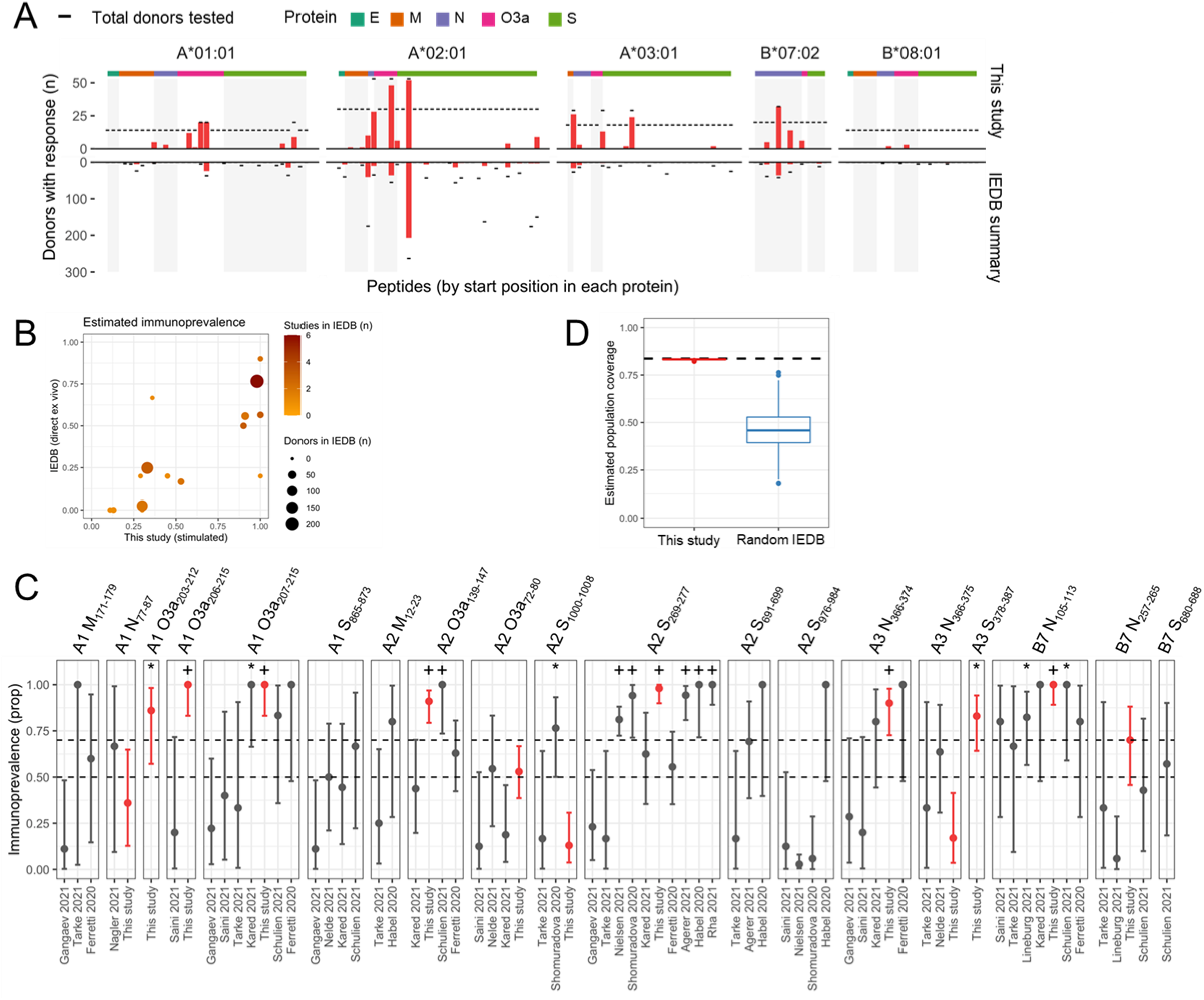
Comparison with published data and population coverage. (A) Comparison of CD8 responses to epitopes identified in the current study with previously published data from the Immune Epitope Database (IEDB). For all epitopes the total number of individuals tested is shown as black lines and the number of responding donors as red bars. Our data are shown on top; a summary of studies IEDB below. Epitopes are grouped on the x-axis according to HLA-restriction and sorted by protein of origin and amino acid position. (B) Immunoprevalence in our study (x-axis) and in IEDB (y-axis), including only data from IEDB where direct *ex vivo* multimer staining was performed, in contrast to our *in vitro* expansion strategy. Immunoprevalence estimates for the same epitopes in our study vs IEDB (median 41 vs 20 %, Wilcoxon signed rank test p = 0.002). (C) Immunoprevalence data with exact binomial 95 % confidence intervals from individual studies for every epitope reported in 50 % or more of individuals in our study or IEDB^9, 10, 12–22, 32^, among epitopes tested in our study. Our data in red, other studies in gray. Binomial tests were performed to assess whether the immunoprevalence in each study was significantly higher than 0.5 and 0.7, respectively (*: p<0.05 for immunoprevalence >50%; +: p<0.05 for immunoprevalence >70%). (D) Coverage in the European Caucasian population for immunoprevalent peptides identified in this study as compared with 1000 random sets of immunogenic peptides from IEDB. The dashed black line represents population coverage based solely on HLA allele frequencies. Red box plot represents estimated population coverage with our top 9 epitopes (immunoprevalence 70% or higher) after adjusting for immunoprevalence, using bootstrapping to quantify uncertainty (1000 iterations). In blue are results from 1000 random draws of 9 immunogenic epitopes from IEDB, while keeping the HLA allele distribution the same as for our top 9 epitopes. Boxplots show median and inter-quartile range (IQR); whiskers extend to +/− 1.5 times the IQR and more extreme values are drawn as dots.

### Immunoprevalence of epitopes is a key determinant of vaccine population coverage

Population coverage after vaccination against a set of epitopes depends on 1) the frequency of relevant HLA alleles in the population, and 2) the probability that an individual will mount an immune response to at least one epitope, given their combination of HLA alleles. Based on HLA allele frequency distribution alone, we estimated that our set of nine immunoprevalent epitopes could induce an immune response in 83.7% of the European Caucasian population^39^. Assuming that immunoprevalence in the convalescent setting can be used as proxy for vaccine responses, we estimated an adjusted population coverage of 83.3% (95% CI 82.8-83.5%) for our top nine epitopes (Methods; Suppl Data 1; Fig. 4D; Suppl Fig. 7A+B). This is almost identical to the coverage based solely on HLA allele frequency, since the immunoprevalence for these nine epitopes is very high. We next evaluated the impact of immunoprevalence on vaccine coverage by calculating the adjusted population coverage of 1000 random sets of nine immunogenic epitopes from IEDB with varying immunoprevalence, keeping the distribution identical to that of the HLA alleles restricting our top epitopes. This demonstrated a striking variation in population coverage despite identical HLA background, ranging from 17.8% to 75.5% with a mean of 46.3% across random epitope sets; approximately half of the coverage that was achieved with an optimal epitope set (Fig. 4D). Hence, development of T-cell vaccines should look beyond the binary question of whether an epitope is immunogenic or not, and consider immunoprevalence a key factor to identify epitopes that will provide broad population coverage.

## DISCUSSION

Vaccines that induce broad T-cell responses may boost immunity as protection from current vaccines against SARS-CoV-2 is waning. From a manufacturing standpoint, and to deliver the highest possible dose of the most immunogenic antigens, it is rational to limit the number of epitopes to those inducing the strongest immune responses in the highest proportion of individuals in a population. Our data show that the CD8 T-cell response to SARS-CoV-2 is more focused than previously believed. We identified nine highly immunoprevalent (≥70%) SARS-CoV-2 specific CD8 T-cell epitopes restricted by four of the most prevalent HLA class I alleles in Caucasians.

To our knowledge, only one other study investigated the immunogenicity of eluted HLA ligands in COVID-19 convalescents, where two peptides were tested for tetramer binding in 3-5 COVID-19 patients^12^. Our data show that eluted peptides were no more likely to be immunogenic than those included solely on basis of high predicted binding affinity (Fisher’s exact test, p=0.7 and p=0.5, respectively). We did, however, identify two epitopes by MS that would have been excluded using prediction only as they were predicted to be weak binders. Seventeen peptides identified by MS also had high predicted HLA-binding affinity, of which 10 were immunogenic (58%). Combining MS and HLA binding prediction could therefore provide an efficient strategy to identify peptides with a very high likelihood of being immunogenic.

We developed an algorithm that predicts total CD8 T-cell response coverage for a given set of epitopes in a population with known HLA distribution, when immunoprevalence for the epitopes has been determined. Among the 29 SARS-CoV-2-specific epitopes identified here, we predict that nine epitopes give an immune response in 83% of Caucasians. This prediction is in good agreement with results from a recent study showing that the two epitopes from the Spike protein were among the most immunoprevalent after mRNA vaccination^50^. However, only 2/9 epitopes detected here are derived from the Spike protein. Broader protection could thus be induced if epitopes from additional SARS-CoV-2 proteins are included in vaccines. Finally, although we included a large number of COVID-19 convalescents representing the most frequent HLA alleles in a Caucasian population, development of a T-cell vaccine with global potential will require data representative of additional HLA alleles that are prevalent in other populations than the one studied here.

## MATERIALS AND METHODS

### Study design

Following the discovery of SARS-CoV-2, we set out to identify candidate epitopes for a CD8 T-cell vaccine in a large cohort of COVID-19 convalescents and healthy controls. Samples were prospectively collected as part of a large biobanking effort in Norway. We performed no formal sample size calculations and individuals were consecutively enrolled. Data from all individuals with sufficient available biological material and where experiments were technically successful were included in downstream analysis. Experimental and statistical analysis design is outlined in detail in respective sections below.

### Subject details

The project (REK # 124170) was approved by the Regional Research Ethics Committee according to the Declaration of Helsinki. Eligible participants provided informed consent by signing an online electronic consent form. Blood collection of all individuals was performed by the Oslo Blood Bank.

#### Healthy pre-pandemic (Pre) controls

Buffy-coats from healthy donors sampled before the pandemic (n = 30) were obtained between 2015-2018 and PBMC isolated using a standard Ficoll isolation protocol. PBMC were cryopreserved in 60% FBS / 30% RPMI / 10% DMSO and stored in liquid nitrogen.

#### Healthy pandemic (pan) control donors, and convalescent COVID-19 donors

Blood from healthy pandemic controls with a SARS-CoV-2 negative PCR test (n = 14), or from 96 convalescent individuals (22 hospitalized) with mild to severe symptoms and at least 14 days without symptoms, was collected in ACD, CPTA or CPT tubes and processed following the manufacturer’s instructions. Convalescents were included after confirmation of a positive PCR test (n=93). Three convalescents for whom a PCR test was lacking were included based on high anti-RBD and anti-Nucleocapsid antibody titers. After initial centrifugation, serum samples were taken and stored at 4°C. PBMC were frozen in 60% FBS / 30% RPMI / 10% DMSO and stored in liquid nitrogen. There were no differences between convalescent individuals who had been hospitalized due to COVID-19 as compared to those who had not with respect to SARS-CoV-2 antibody titers or functional CD4 and CD8 T cell responses to peptide pools (Suppl Fig. 1B and 2C). We made similar findings when comparing pre-pandemic and other healthy control samples (Suppl Fig. 2D). We therefore considered all convalescent and healthy donors, respectively, as single cohorts in the relevant analyses.

### Antibodies

RE goat-anti-human IgG-Fc (Jackson Immunoresearch); PE anti-human HLA-A,B,C (clone W6/32; dilution 1:30; Biolegend; Cat no 311406), PerCP/Cyanine5.5 anti-human CD3 (clone SK7; dilution 1:100; Biolegend; Cat no 344808), Alexa Fluor® 700 anti-human CD4 (clone RPA-T4; dilution 1:200; Biolegend; Cat no 344808), FITC anti-human CD8a (clone RPA-T8; dilution 1:100; Biolegend; Cat no 301050); Alexa Fluor 647 anti-human CD137 (clone 4B4-1; dilution 1:100; Thermo Fisher Scientific; Cat no A51019); PE anti-human CD134 (clone OX40; dilution 1:100; Biolegend; Cat no 350004), BV421 anti-human CD69 (clone FN50; dilution 1:100; BD Biosciences; Cat no 562884); BV605 anti-human HLA-DR (clone L243; dilution 1:100; Biolegend; Cat no 307640); BV785 anti-human CD38 (clone HIT2; dilution 1:100; Biolegend; Cat no Biolegend); BV785 anti-human CD19 (clone HIB19; dilution 1:100; Biolegend; Cat no 302240); BV785 anti-human CD56 (clone 5.1H11; dilution 1:100; Biolegend; Cat no 362550); BV785 anti-human CD14 (clone M5E2; dilution 1:100; Biolegend; Cat no 301840); BV785 anti-human CD4 (clone RPA-T4; dilution 1:100; Biolegend; Cat no 300554); PE anti-human-CD8 a (Clone RPA-T8; dilution 1:200; Biolegend; Cat no 301008); FITC anti-human CD20 (clone 2H7; dilution 1:200; Biolegend; Cat no 302304)

### Plasma antibody titer determination

A multiplexed bead-based flow cytometric assay, referred to as microsphere affinity proteomics (MAP), was adapted for detection of SARS-CoV-2 antibodies^51^. Amine-functionalized polymer beads were color-coded with fluorescent dyes as described earlier, and reacted successively with amine-reactive biotin (sulfo-NHS-LC-biotin, Proteochem, USA) and neutravidin (Thermo Fisher). A DNA construct encoding the receptor-binding domain of Spike-1 protein (RBD) from SARS-CoV-2 was provided by Florian Krammer, and the protocol described by Amanat and colleagues^52^ was used to produce recombinant protein in Expi293F cells. Bacterially expressed full-length Nucleocapsid from SARS-CoV-2 was purchased from Prospec Bio (https://www.prospecbio.com). Viral proteins solubilized in PBS were biotinylated chemically using a 4:1 molar ratio of sulfo-NHS-LC-biotin to protein. Free biotin was removed with G50 sephadex spin columns. Biotinylated proteins were bound to neutravidin-coupled microspheres with fluorescent barcodes. Beads with Neutravidin only were used as reference for background binding. The bead multiplex was incubated for 1 h with serum diluted 1:1000 in PBS supplemented with Neutravidin (10 μg/mL), d-Biotin (10 μg/mL), bovine serum albumin (1%) and sodium azide (0.1%). The beads were washed twice in PBS with 1% Tween 20 (PBT), labelled with PE-conjugated goat-anti-human IgG-Fc for 20 min, washed again and analyzed by flow cytometry (Attune Next, Thermo Fisher). Specific binding was measured as the ratio of R-Phycoerythrin fluorescence intensity of antigen-coupled beads and neutravidin-only beads. Samples containing antibodies both to Nucleocapsid and RBD were considered to be positive. Reference panels containing samples from 287 individuals with PCR-confirmed SARS-CoV-2 infection and 1343 pre-pandemic samples were used to set the cutoff. A cutoff of five and ten for RBD and Nucleocapsid respectively, yielded a specificity of 100%. The sensitivity was 84%^53^.

### HLA typing

DNA was isolated from 1 × 10^6^ PBMC using the DNeasy Blood & Tissue Kit (Qiagen, Cat No 69506), following the manufacturer’s instructions. HLA typing was performed by NGS sequencing using NGSgo®-AmpX HLA GeneSuite (NGSgo®-AmpX HLA-A, B, C, DRB1, DQB1 & NGSgo®-AmpX HLA-DPA1, DPB1, DQA1, DRB3/4/5 kits) for sample library preparations and ran on a Miseq sequencer (Illumina), following the manufacturer’s instructions.

### HLA mono-allelic B721.221 cells

HLA class I-deficient B721.221 cells (IHW00001, FRED HUTCH Research Cell bank) were retrovirally transduced to express single HLA alleles (HLA-A*03:01, HLA-A*01:01, HLA-A*02:01, HLA-B*07:02, HLA-B*08:01; Gene Bank HG794390, HG794373, HG794376, HG794392, and HG794374, respectively). Cells were retrovirally transduced using a previously published protocol^54^. After staining with a PE anti-human HLA-A,B,C antibody, stable HLA expressing cell lines were sorted using the Sony SH800 cell sorter. The HLA mono-allelic B721.221 cells were then transduced with plasmids coding for SARS-CoV-2 proteins (Spike 1 (aa 1-541), Spike 2 (aa 490-975), Spike 3 (aa 888-1273), Spike, Nucleocapsid, Membrane-T2A-Envelope, and ORF3a; GenBank: MN908947.3) under the control of a tetracycline-inducible promoter (Retro-X™ Tet-One™ Inducible Expression System, Takarabio) with a puromycin selection marker. Transduced cells were selected by culturing the cells in the presence of 2 μg/mL puromycin (replenished every 48 h) (Gibco; Cat no A1113802) for 7-14 days, followed by further expansion in medium containing 0.5 μg/mL puromycin.

### Mass-spectrometry

HLA mono-allelic B721.221 cells transduced with SARS-CoV-2 proteins were expanded to 100 × 10^6^ cells and expression of the target proteins was induced by culturing the cells in the presence of doxycycline (1 μg/mL; replenished every 24 h) for 48 h. The cells were then lysed in 1 mL of lysis buffer (PBS containing 1% lauryl maltoside, 1 mM EDTA, 1 mM PMSF and 1:200 Sigma protease inhibitor) for 1 h on ice. HLA peptide complexes were purified by immunoprecipitation^55^. Briefly, the HLA peptide complexes were captured on to beads coated with pan HLA class I antibody, and the beads were then washed with 3mL each of 0.1 M Tris-HCl / 150 mM NaCl, 0.1 M Tris-HCl / 400 mM NaCl, again with 0.1 M Tris-HCl / 150 mM NaCl, and finally 0.1 M Tris-HCl. All peptide elutions were desalted with Discovery DSC-18 SPE column, vacuum concentrated and dissolved in 25 μL of 3% acetonitrile containing 0.1% FA. The peptide solution (5 μL) was analyzed using an Ultimate 3000 nano-UHPLC system (Dionex, Sunnyvale, CA, USA) connected to a Q Exactive mass spectrometer (ThermoElectron, Bremen, Germany) equipped with a nano electrospray ion source as described previously^55^. A flow rate of 300 nL/min was employed with a solvent gradient of 3-35% B in 53 min, to 50% B in 3 min and then to 80% B in 1 min. The samples were also analyzed using a longer solvent gradient of 3-35% B in 100 min, to 50% B in 13 min and then to 80% B in 2 min. Solvent A was 0.1% formic acid and solvent B was 0.1% formic acid / 90% acetonitrile. The mass spectrometer was operated in the data-dependent mode to automatically switch between MS and MS/MS acquisition. The method used allowed sequential isolation of up to the twelve most intense ions, depending on signal intensity (intensity threshold 1e5), for fragmentation using higher-energy collision induced dissociation (HCD) at a resolution R = 17,500 with NCE 27. The raw data was then analyzed with PEAKS software (Bioinformatics Solutions Inc). The tandem mass spectra were matched against the Uniport homo sapiens database appended with the SARS-CoV-2 proteins. Precursor mass tolerance was set to 10 ppm, methione oxidation was considered as variable modification, enzyme specificity was set to none and a product ion tolerance of 0.05 Da was used. SARS-CoV-2 peptides identified from the discovery approach were further validated using synthetic peptide analogs and raw data were analyzed using the Xcalibur^TM^ software.

### Synthetic peptides

#### Peptide megapools

15-mer peptide libraries with 11 amino acids overlap covering five SARS-CoV-2 proteins (Spike, Nucleocapsid, Envelope, Membrane, and ORF3a; GenBank: MN908947.3) were synthesized at Genscript (purity >70%), dissolved in DMSO and pooled as megapools (MP) (S_1-327_; S_317-647_; S_637-963_; S_953-1273_; O3a_1-275_; E_1-75_; M_1-115_; M_105-222_; N_1-211_; N_201-419_; 1 mg/mL per peptide).

#### Single peptides

Peptide selection was based on a) peptides identified by Mass-spectrometry and b) 9- and 10-mers predicted to display the highest binding affinity to the MHC according to NetMHCpan4.1 (BA_Rank <0.2%) and/or NetMHC4.0 (Rank <0.25%) (Suppl Table 5). One additional peptide was included based on early published reports^37^ . HLA-A*01:01, HLA-A*02:01, HLA-A*03:01, HLA-B*07:02, and HLA-B*08:01 were included to cover the most prevalent HLA alleles in the Caucasian population. In total, 132 peptides derived from five SARS-CoV-2 proteins (Spike, Nucleocapsid, Envelope, Membrane, and ORF3a) were successfully synthesized at Genscript (purity >70%) and dissolved in DMSO.

### Determination of *in vivo* activation status of PBMC

1 × 10^6^ freshly thawed PBMC of COVID-19 convalescent (n=96) and healthy control samples (pre-pandemic n=19 and pandemic n=14) were incubated with 10 μL undiluted Human Fc Receptor binding inhibitor (AH Diagnostics; Cat no 14-9161-73) for 10 min at room temperature. Subsequently, surface staining was performed by adding the following antibodies: PerCP/Cyanine5.5 anti-human CD3, Alexa Fluor® 700 anti-human CD4, FITC anti-human CD8a, BV605 anti-human HLA-DR, BV785 anti-human CD38, BV421 anti-human CD69 together with Live/Dear-near IR (1:1000 in PBS; LD-NIR; ThermoFisher Scientific; Cat no L10119) for exclusion of dead cells. After staining the samples for 20 min at 4°C, cells were washed two times with flow buffer (50 nM EDTA, 5% FBS in PBS) before acquiring them on the FACSymphony A5 (BD Biosciences). The gating strategy is outlined in Suppl Fig 1C.

### Functional T cell response (AIM assay)

AIM assays were performed using cryopreserved PBMC of COVID-19 convalescent (n = 96) and healthy control donors (19 pre-pandemic and 14 pandemic). On Day 0, 7.5 × 10^5^ PBMC were plated in 200 μL IMDM w/ L-Glutamine and HEPES (ThermoFisher Scientific; Cat no 12440061) supplemented with 1x P/S and 5% normal human serum (Trina Bioreactives AG; Cat no SN0300, Lot 12KSN27546) (referred to as complete medium) in polypropylene coated 96-well U-bottom plates in the presence of peptide megapools (each peptide at 150 ng/mL) covering the different proteins (1 well/pool/donor). DMSO was used as negative control (1-2 wells/donor). On Day 3, half-medium exchange was performed and 10 IU/mL IL-2 (R&D Systems; Cat no 202-IL-500) added. On Day 5, PBMC were restimulated by adding the relevant peptide megapools (each peptide at 375 ng/mL). 20-24 h post restimulation, PBMC were harvested, washed with PBS, blocked for 10 min with Human Fc Receptor binding inhibitor at room temperature and subsequently stained with PerCP/Cyanine5.5 anti-human CD3, Alexa Fluor® 700 anti-human CD4, FITC anti-human CD8a; Alexa Fluor 647 anti-human CD137; PE anti-human CD134, BV421 anti-Human CD69 and LD-NIR for exclusion of dead cells. Samples were acquired using a FACSymphony A5 (BD Biosciences). Responding T cells were gated as % CD137^+^CD134^+^ events among live CD4^+^ events, and % CD137^+^CD69^+^ among live CD8^+^ events. Specific activation responses were determined by subtracting background activation in DMSO from the response observed following peptide-stimulation, for each patient. Gating strategy is outlined in Suppl Fig. 2B.

### Combinatorial multimer staining

#### Monomer and Multimer production

Soluble biotinylated Class-I MHC monomers containing a UV-cleavable peptide in their binding groove were produced in-house according to published protocols^56, 57^. Stocks were frozen at −80°C until further use. Final concentrations of the UV-monomer and peptide diluted in PBS in the exchange reaction were 25 μg/mL and 50 μg/mL, respectively. The UV-dependent peptide exchange was performed at a wavelength of 366 nm for 1 h at 4°C. 24 h after UV-exchange, the following streptavidin-tagged fluorochromes were added at optimized ratios to the pMonomer solution: SA-PE (Invitrogen; Cat no S866; Lot no 2129894); SA-APC (Invitrogen; Cat no S868; Lot no 2105223), SA-BV421 (Biolegend; Cat no 405229; Lot no B202410), SA-BV605 (Biolegend; Cat no 405225; Lot no B267737), SA-BV421 (Biolegend; Cat no 405229; Lot no B202410), SA-PE Cy7 ( BD Biosciences; Cat no 557598; Lot no 7325636), SA-PE-CF594 (BD Biosciences; Cat no 562284; Lot no 7138598), SA-PE-Cy5 (BD Biosciences; Cat no 554062; Lot no 7066693), SA-APC-R700 ( BD Biosciences; Cat no 565144; Lot no 7179560), SA-BB790-P (BD Biosciences; prototype kindly provided by Bob Balderas from BD Biosciences; Lot no 8124567). To block unoccupied binding sites, D-biotin (20 μM; Avidity; Cat no I2011) was added 24 h after multimerization. Plates were stored in the dark at 4°C until use.

#### Multimer staining assay

Among the 96 convalescent individuals included in the functional T cell response analysis, 11 did not express the prevalent HLA alleles included in the multimer staining assays, and for two no HLA typing data were available. Thus, PBMC from 83 COVID-19 convalescents (Cohort 1+2) and 19 healthy pre-pandemic samples were included in the multimer staining assays. For 7 convalescent individuals two sampling time points were included (on average 56 [38-81] days between sampling). On Day 0 PBMC were thawed and resuspended at 1 × 10^6^ c/mL in IMDM w/ L-Glutamine and HEPES and loaded for 2 h at 37°C with the peptide master mix containing all 132 pre-selected peptides (each peptide at 100 ng/mL). Cells were washed and plated out in multiple wells at 7.5 × 10^5^ c/well (in 200 μL) in complete medium in a polypropylene coated 96-well U-bottom plate. On Day 3, half-medium exchange was performed and 10 IU/mL IL-2 added. On Day 5, medium was completely replenished. On Day 7, cells from each donor were pooled, washed and 2-3 × 10^6^ PBMC were resuspended in 50 μL PBS (polystyrene coated 96-well V-bottom plate) and stained with separate mixes of multimers complexed with 13-29 distinctive peptides, with each multimer conjugated to dual fluorochrome combinations. 1 μL/sample of PE-Cy7 / PE-CF594 / PE-Cy5 / APC-R700 and 2 μL/sample for BV421 / BV605 / APC / PE / BB790- tagged multimers were added. The sample was incubated for 10 min at room temperature in the dark. Thereafter the following antibodies and LD-NIR (1:1000) were added to the sample for 30 min at 4°C: FITC anti-human CD8a, BV785 anti-human CD19, BV785 anti-human CD56, BV785 anti-human CD14, BV785 anti-human CD4. The samples were washed three times with flow buffer before acquiring them on the BD Symphony A5. The gating strategy is outlined in Suppl Fig. 3. The sample was included in the analysis if at least 4000 live CD8 cells were acquired. An individual was classified as responder to a peptide if the tetramer population had 1) at least 5 clearly double-positive events (each multimer conjugated to two different fluorochromes), 2) constituted ≥0.005% of the live CD8, and 3) formed a tight cluster.

### Sorting of peptide-specific T cells as cell lines

PBMC were stimulated and stained with multimers and antibodies as described above (section “Multimer staining assay”). Multimer positive cells were sorted onto irradiated feeders (PBMC from 3 donors at equal ratios; 300 kV 10 mA 7 min in X-Vivo-20 (BioNordika; Cat no BE04-448Q) supplemented with 1x P/S, 5% HS, 1 μg/mL PHA (Remel, Cat no 30852801), 875 U/mL IL-2 and 2 ng/mL IL-15 (Peprotech; Cat no 200-15)) as bulk T cells on a FACS Aria II cell sorter (BD Biosciences), followed by expansion in X-Vivo-20 supplemented with 1x P/S, 5% HS, 437.5 U/mL IL-2 and 1 ng/mL IL-15 for generation of T-cell lines.

### Functional analysis of SARS-CoV-2-specific T-cell lines

SARS-CoV-2-specific T-cell lines (confirmed by multimer staining to recognize peptide of interest) were combined with mono-allelic B721.221 cells loaded or not with peptide (2 h; 100 ng/mL), or retrovirally transduced to express the relevant SARS-CoV-2 antigen (see section “HLA mono-allelic B721.221 cells”) (treated with doxycycline for 48 h to express the antigen) for 20 h (n=1, duplicate or triplicate, depending on available cell numbers; effector to target ratio (E:T) = 1:1). Harvested cells were incubated with Human Fc Receptor binding inhibitor for 10 min at room temperature. Subsequently, surface staining was performed by adding the following antibodies: PE anti-human-CD8a, Alexa Fluor 647 anti-human CD137, FITC anti-human CD20, together with LD-NIR. After staining the samples for 20 min at 4°C, cells were washed twice with flow buffer and acquired on the LSR II Yellow laser (BD Biosciences). Killing of target cells was quantified by adding CountBright™ Absolute Counting Beads (Invitrogen, Cat no C36950) following the manufacturer’s instructions before acquisition of sample. A fixed number of beads was acquired for each well. The number of acquired target cells was normalized to the matching mock control (w/o AG) and displayed as % remaining viable cells. The gating strategy is outlined in Suppl Fig. 6a.

### Peptide sequence alignment with human common cold coronaviruses

Homology calculations were performed by aligning the sequence of SARS-CoV-2 epitopes with the corresponding protein sequences of common cold coronaviruses (OC43, NL63, HKU1,229E) using the Clustal Omega tool with default parameters. The following sequences were used: OC43 (Spike: P36334, Nucleocapsid: P33469, Membrane: Q01455), NL63 (Spike: Q6Q1S2, Nucleocapsid: Q6Q1R8, Membrane: Q6Q1R9, ORF3a: Q6Q1S1), HKU1 (Spike: Q0ZME7, Nucleocapsid: Q0ZME3, Membrane: Q0ZME4), 229E (Spike: P15423, Nucleocapsid: P15130, Membrane: P15422). The percent homology between SARS-Cov-2 epitopes and common cold corona viruses was calculated by the number of matching amino acid residues.

### Immune Epitope Database analysis

We performed an extensive search for previously published data on T-cell responses to SARS-CoV-2 epitopes in the Immune Epitope Database IEDB (http://www.iedb.org). The data cut-off date was 27^th^ of July 2021. We used the following search criteria in the IEDB search engine:

Epitope: Linear peptide
Assay: T cell
Outcome: positive and negative
Organism: SARS-CoV-2
MHC restriction: class 1
Host: Human
Disease: Any

IEDB entries were removed from further analysis if immunogenicity data could not be mapped unambiguously to a single peptide and HLA allele, and if there was missing data regarding the number of donors tested or the number of donors with an immune response. Furthermore, we included only entries where disease state was specified as COVID-19 and disease stage as “post” or “active/recent onset”. One study was missing from IEDB, reporting *ex vivo* validation of an epitope identified by MS^12^. Data for this single epitope was manually added to IEDB output for downstream analysis. We first generated summary statistics for each peptide, HLA allele and study. If a given peptide/HLA combination was tested by more than one assay in a single study, we moved forward with the highest number of donors tested and the highest number of donors with an immune response. To generate pooled data for each peptide/HLA, data from individual studies were summarized. For quality control purposes we also performed analysis considering each individual assay performed, e.g., calculating separate statistics considering only data from direct *ex vivo* HLA multimer staining. When evaluating the impact of immunoprevalence on estimated population coverage in random sets of epitopes from IEDB, we included only epitopes with data from >20 individuals and/or >1 study.

### Estimating population coverage adjusted for immunoprevalence

Suppose a vaccine is made using the epitope set *E_K_* = {*e*_1_, *e*_2_ … *e_K_*}. Every individual has *H* different loci, and two alleles at each locus. An epitope may be independently presented on one or more of these alleles, at any given locus, and we assume that one response-generating hit (hit = allele-epitope match) is sufficient for the vaccine to generate protection. Given this, we can calculate the probability of response *P*(*R*) from at least one locus as:

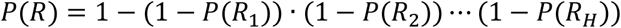

where *P*(*R*_ℎ_) is the probability of response to an epitope presented on locus ℎ. We can also write it as:

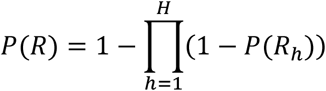

To calculate *P*(*R*_ℎ_), we need to know the relative population frequencies of every possible allele combination for that locus, as well as the probability that each allele combination generates an immune response to at least one epitope. HLA allele frequencies were obtained from the Allele Frequency database^58^ and scaled to 1 for each locus. If we let 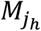 denote one of all the *J*_ℎ_ possible allele combinations occurring in the population for locus ℎ, and let 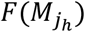 denote the population percentage with this combination, then since allele combinations are mutually exclusive events, we can write the probability of response with a given locus as:

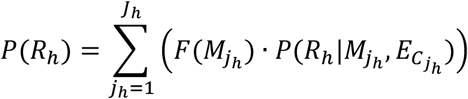

where 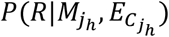 is the probability of an immune response from at least one epitope-allele pair, given the allele combination 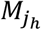 and its set of compatible epitopes 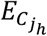, where 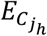 is a subset of *E_K_*. To estimate the probability that a presented epitope results in a response, we used immunoprevalence data from convalescent individuals. If we let *τ*(*a*, *e_k_*) be the immunoprevalence for a given allele-epitope pair, we can write the probability that an individual with the allele combination 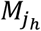 generates an immune response to the epitope *e_k_* at locus ℎ as:

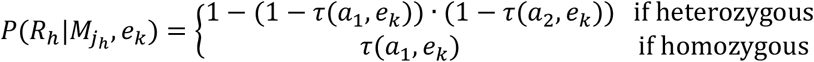

where *a*_1_ and *a*_2_ are the first and second alleles found at locus ℎ, respectively. Note that if an individual is heterozygous, and both alleles can present the epitope, both alleles have an independent chance of generating a response. If an allele does not bind with *e_k_*, then *τ*(*a*, *e_k_*) = 0. Since each allele combination 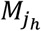 can have multiple compatible epitopes in *E_K_*, every allele combination can get multiple attempts at generating a response, where the number of attempts is 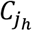. We can then write the probability that an individual with the allele combination 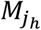 generates a response from *at least one* of the epitopes in 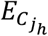 as:

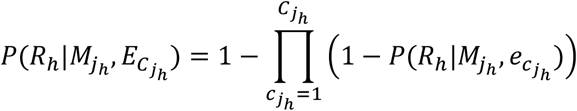

This is assuming that 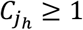. If there are no compatible epitopes 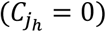, then 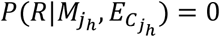. Once all the weights have been calculated for all the loci, we can substitute this into the first equation to get a point estimate for the predicted population coverage, given a vector of alleles, their population frequencies, an epitope set, and their associated immunoprevalences for different alleles. To compute a confidence interval for the predicted coverage, we use parametric bootstrapping to resample new immunoprevalence values for each epitope-allele pair. If we assume that the number of patients with a compatible allele who generate a response to an immunogenic epitope is binomially distributed, we can model the distribution of *τ*(*a*, *e_k_*) with the beta distribution:

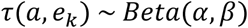

where (α) and (β − 1) are the numbers of responders and non-responders, respectively, from which immunoprevalence was calculated for that allele-epitope combination. Because the beta distribution is not defined when *τ*(*a*, *e_k_*) = 1, we added 1 to the number of non-responders for every epitope. By repeatedly resampling new immunoprevalences from a beta distribution with shape parameters determined by the immunoprevalence data, this will generate a distribution of population coverages. The 95 % confidence interval is then given by the 2.5^th^ and 97.5^th^ percentiles of this distribution.

### Data analysis and statistics

Flow cytometry data acquired on different BD Biosciences instruments were analyzed using FlowJo (TreeStar) version 10.6.2 software. Statistical analysis was performed in R version 4.0.4 and GraphPad Prism version 8.3.0. The use of statistical tests including p-value thresholds and adjustment for multiple testing is specified in the text and figure legends. In general, we used Fisher’s exact test for contingency tables and Wilcoxon rank-sum test for non-parametric continuous variables. For correlation analysis we used the Pearson method whenever its assumptions were met, otherwise we used Spearman rank correlation. We reasoned that the proportion of donors showing an immune response to a particular epitope could be modeled by the binomial distribution, allowing us to estimate binomial 95% confidence intervals for immunoprevalence in our data as well as studies reported in IEDB. Moreover, we used binomial tests for the hypothesis that the true immunoprevalence of each epitope is higher than given thresholds (i.e., 50% and 70%). Adjustment for multiple comparisons was done using the Benjamini-Hochberg method when specified in text or figure legends.

## Supporting information

R code Population coverage

## Supplementary Materials

Fig. S1. Antibody responses and lack of recent *in vivo* activation of T cells.

Fig. S2. Functional CD4 and CD8 T-cell responses to peptide megapools.

Fig. S3. Gating strategy for combinatorial multimer staining approach.

Fig. S4. Representative flow plots of immunogenic SARS-CoV-2 peptides presented by HLA-A*01:01, HLA-A*02:01, HLA-A*03:01, HLA-B*07:02 or HLA-B*08:01 identified by multimer staining.

Fig. S5. Immunoprevalence evaluated by binomial tests.

Fig. S6. Gating strategy for functional validation of T-cell lines (A), cross-recognition of T-cell lines of length variants (B), and T-cell epitope alignment (C).

Fig. S7. Uncertainty in immunoprevalence estimates and population coverage.

Table S1. Information for all samples (COVID-19 convalescent individuals and healthy controls) included in current study.

Table S2. Description of cohort included in T-cell analysis using a functional readout (AIM assay).

Table S3. Mono-allelic B721.221 cell lines transduced to express SARS-CoV-2 proteins that were analyzed by MS in this study.

Table S4. Peptides that were identified by MS analysis of mono-allelic B721.221 cell lines expressing the SARS-CoV-2 proteins.

Table S5. Complete list of all pHLA combinations analyzed in the current study along with the criteria employed in selecting these peptides for further immunogenicity testing.

Table S6. Summary of immunogenicity data for all 137 pHLA combinations included in our study.

Table S7. List of lineage defining mutations identified in variants of concern (VOC) and variants under investigation (VUI) according to UK briefings.

File S1. MS/MS fragmentation spectra of synthetic and eluted HLA class I peptides identified from mono-allelic B721.221 cells expressing the SARS-CoV-2 proteins

Data S1. R code and example data to calculate adjusted population coverage

## Acknowledgements

We express our gratitude to all donors and health care personnel involved in this work. We thank Oslo University Hospital (OUH) flow cytometry core facility for excellent technical assistance.

## Funding

South-Eastern Regional Health Authority Norway (JO)

Research Council of Norway (JO)

University of Oslo (JO)

Oslo University Hospital (JO)

The Norwegian Cancer Society (FL-J and JO)

The Coalition for Epidemic Preparedness Innovations (CEPI) (JTV and FL-J)

## Author contributions

Conceptualization: SM, IB, RCB, EHR, JO

Methodology: SM, IB, RCB, TTT, FLJ, JO

Software: TSM, EHR

Resources: SM, MD-S, CK, AG, LN-M, AS

Investigation: SM, IB, RCB, TTT, K-ZD, WY, KD, ML, MMN, JTV, BT, FL-J, JO

Formal analysis: SM, RCB, TSM, EHR

Visualization: SM, RCB, EHR

Funding acquisition: JO

Supervision: JO

Writing – original draft: SM, IB, RCB, EHR, FL-J, JO

Writing – review & editing: SM, IB, RCB, TSM, EHR, TTT, WY, KD, ML, MMN, JTV, BT, MD-S, CK, AG, LN-M, AS, MD-S, FL-J, JO

## Data and materials availability

The data that support the findings of this study are available within the article and its supplementary information files. Raw data are available from the corresponding authors upon reasonable request. Public datasets were derived from . Accompanying this manuscript is an implementation in R of our novel algorithm to estimate population coverage of an epitope set adjusted for immunoprevalence. Other analysis code is available from the corresponding authors upon reasonable request.

