## Supplementary Figures for "Beyond Spike: Identification of nine highly prevalent SARS-CoV-2-specific CD8 T-cell epitopes in a large Norwegian cohort"

**Supplementray Figures**

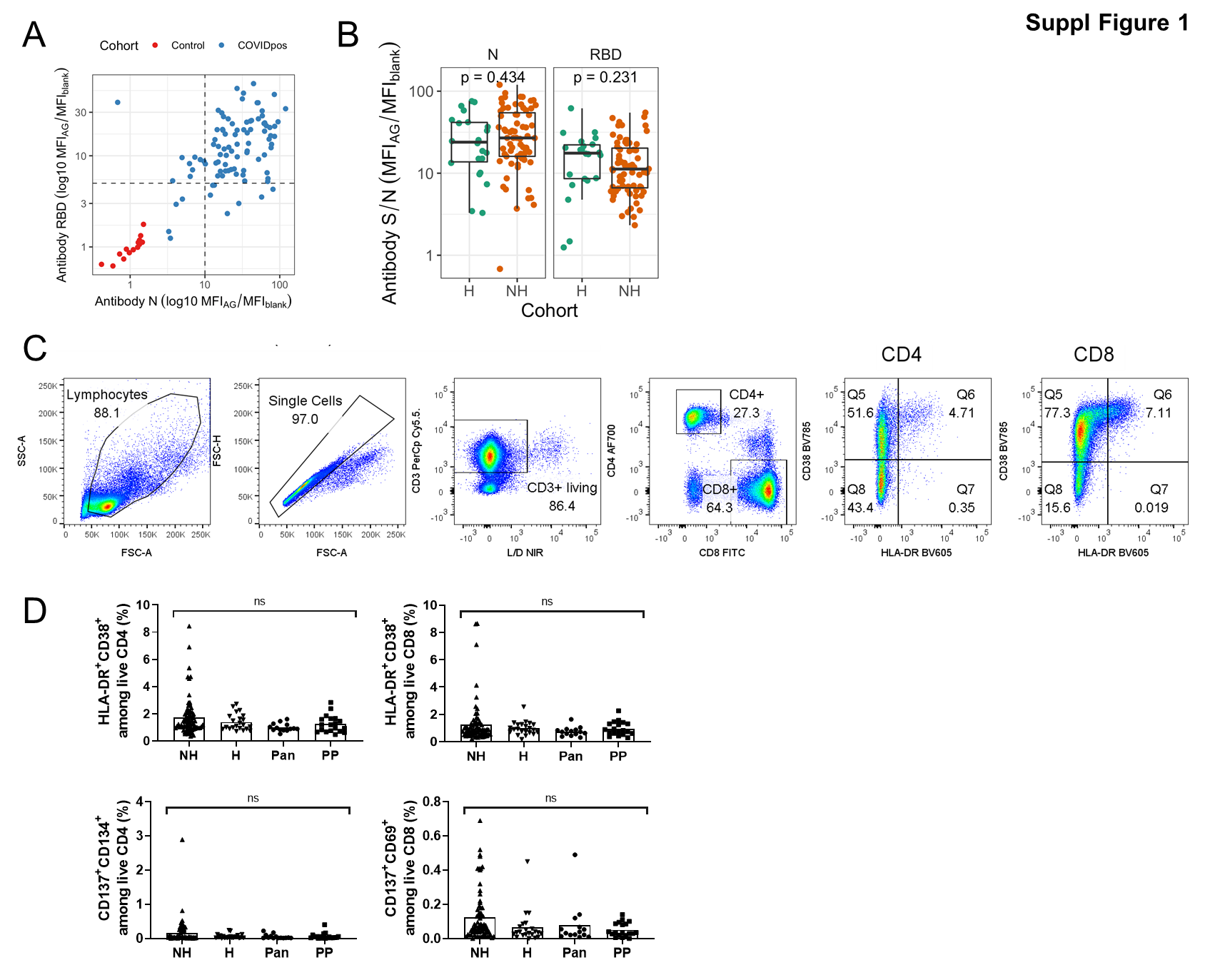

**Supplementary Figure 1.** **Antibody responses and lack of recent *in vivo* activation of T cells**

**(A)** Antibody responses to the receptor-binding domain (RBD) located in the Spike protein (y-axis) and to the Nucelocapsid (N) protein (x-axis). In contrast to individuals with negative SARS-CoV-2 test (pandemic controls; n=14; red), convalescent COVID-19 patients raise antibodies (n=93; blue) against RBD and/or Nucleocapsid. Dashed black lines represent thresholds used to designate positive tests for each of the antibodies. Responses are displayed as ratio of MFI values obtained for specific recognition (MFI_AG_) and blank (MFI_blank_). **(B)** Antibody responses to the receptor-binding domain located in the Spike protein and the Nucleocapsid protein in non-hospitalized (NH) and hospitalized (H) convalescent COVID-19 individuals. Responses are displayed as signal-to-noise ratio. Wilcoxon test was used to compare antibody levels between groups. **(C)** Representative gating strategy used for analysis of recent *in vivo* activation status of T cells (here convalescent individual sample ID 94). **(D)** Low or no recent T-cell activation *in vivo* in convalescent individuals and healthy controls, analyzed by staining of T cells from COVID-19 convalescents (n=96) and healthy controls (n=33; 19 pre-pandemic (PP) and 14 pandemic (Pan)) with T-cell markers (CD3, CD4, CD8) and anti-HLA-DR/-CD38 (top) or anti-CD137/-CD134 (CD4) and anti-CD137/-CD69 (CD8) T cells (bottom), showing no significant differences between cohorts (One-way ANOVA p>0.05).

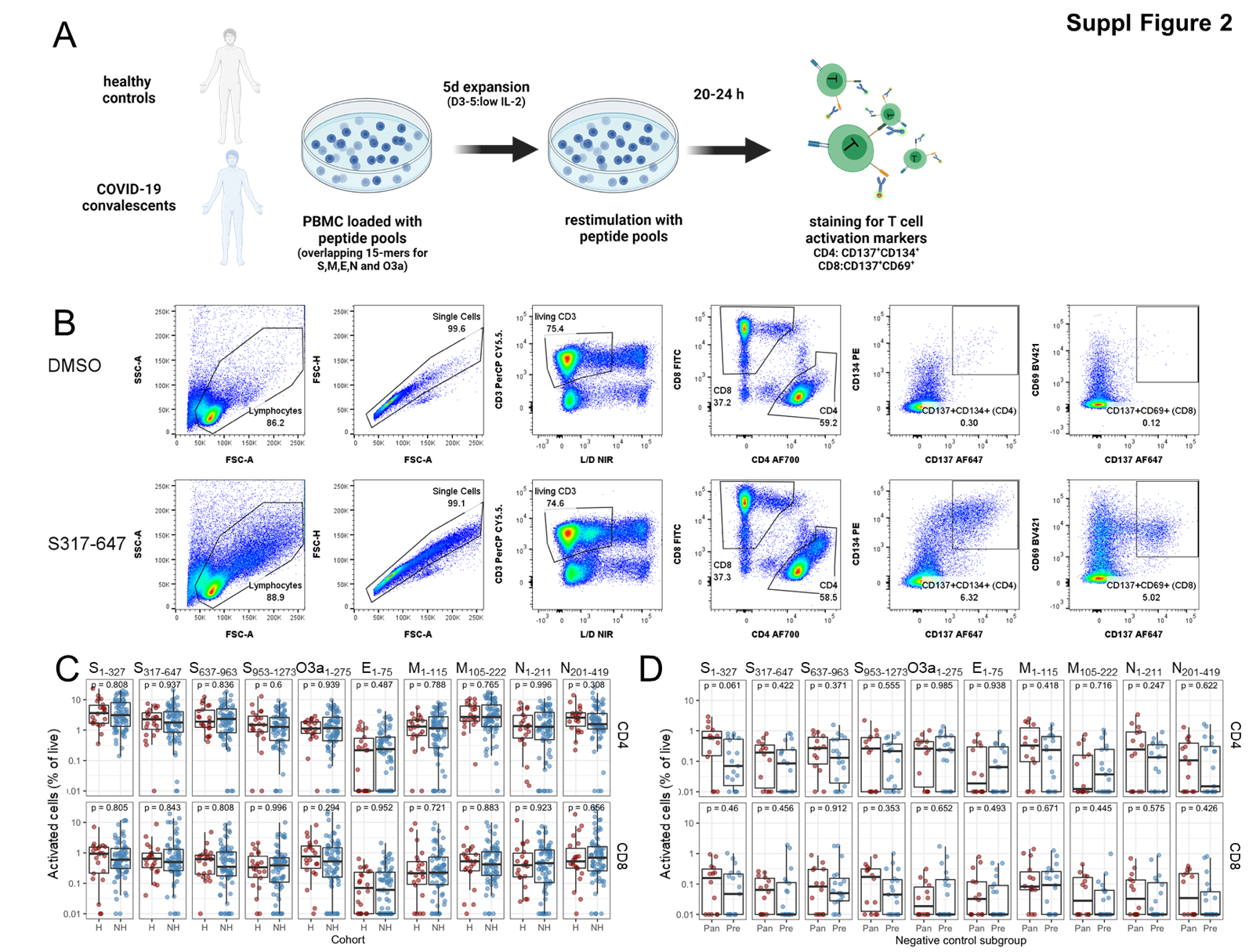

Supplementary Figure 2. Functional CD4 and CD8 T cell responses to peptide megapools

**(A)** Schematic outline of functional T-cell response analysis (AIM assay) to SARS-CoV-2 peptide pools. Thawed PBMC (96 convalescents, 19 pre-pandemic and 14 pandemic controls) were stimulated with peptide megapools (overlapping 15-mers; each peptide at 150 ng/mL) covering the SARS-CoV-2 proteins Spike, Envelope, Membrane, Nucleocapsid and ORF3a (1 well/pool) with a DMSO negative control (1-2 wells/donor). On Day 3, half-medium exchange was performed and 10 IU/mL IL-2 added. On Day 5, PBMC were restimulated by adding the relevant peptide megapools (each peptide at 375ng/mL). At 20-24 h post restimulation, PBMC were stained with T-cell markers CD3, CD4 and CD8 and activation was measured as CD134^+^CD137^+^ and CD69^+^CD137^+^ for live CD4 and CD8 T cells, respectively.
**(B)** Gating strategy for the functional T-cell response analysis. Representative flow plots for a convalescent individual (sample ID 48) stimulated with DMSO control or Spike pool S_317-647_.
**(C+D)** Comparison of functional CD4 and CD8 T-cell responses to peptide pools in hospitalized (H) versus non-hospitalized (NH) convalescent individuals **(C)** and pre-pandemic (Pre) versus pandemic (Pan) healthy controls **(D)**, respectively (Wilcoxon test).

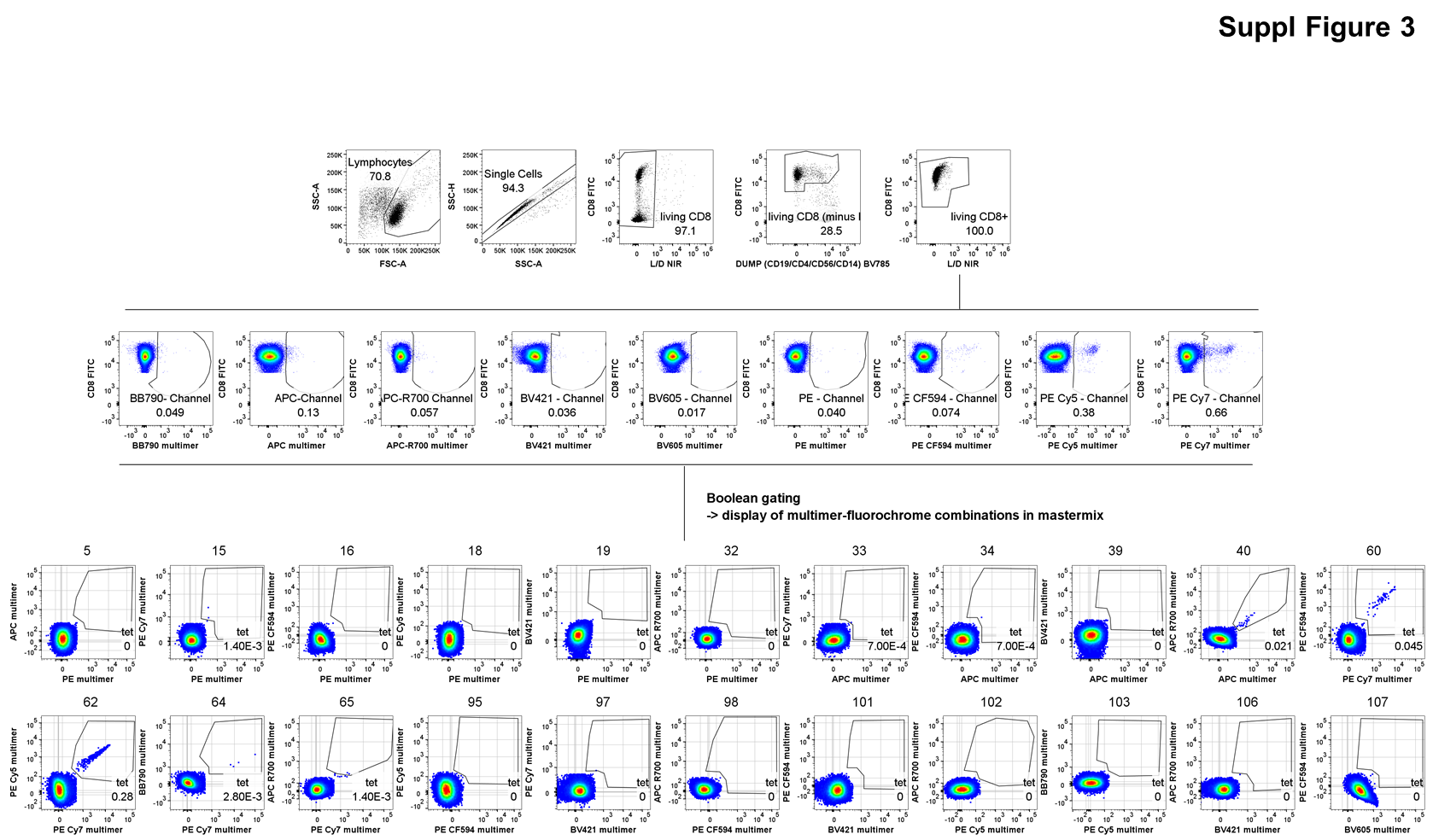

Supplementary Figure 3. Gating strategy for combinatorial multimer staining approach

Representative flow plots for a convalescent individual (sample ID 48) stained with multimers for a subset of the tested HLA-A*02:01 peptides.

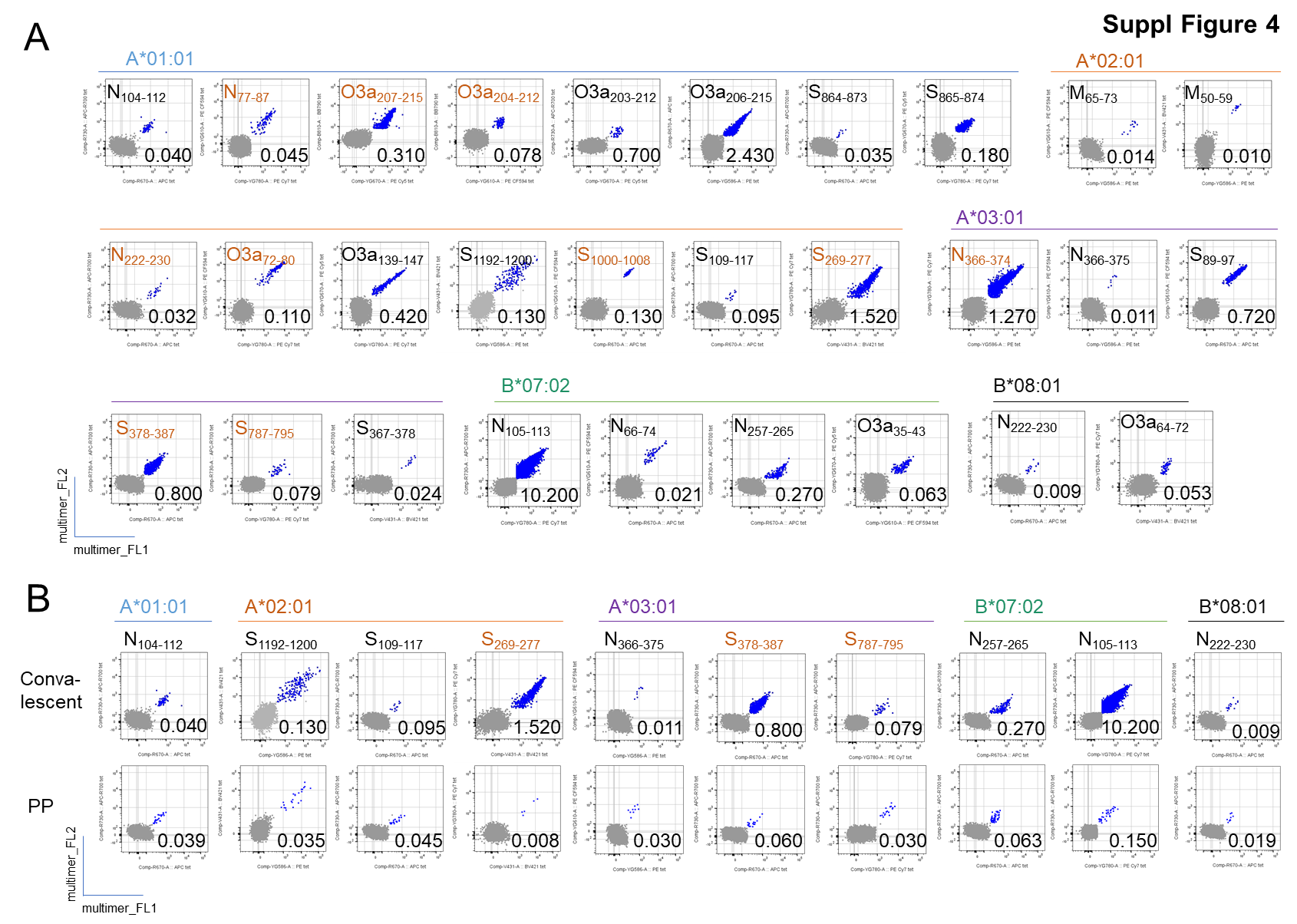

Supplementary Figure 4. **Representative flow plots of immunogenic SARS-CoV-2 peptides presented by HLA-A*01:01, HLA-A*02:01, HLA-A*03:01, HLA-B*07:02 or HLA-B*08:01 identified by multimer staining** **(A)** Representative plots for all immunogenic peptides (n=29) in convalescent COVID-19 individuals. **(B)** Plots for immunogenic peptides (n=10) identified in convalescent COVID-19 individuals (top; same as in **A**) and pre-pandemic (PP) controls (bottom). Peptides identified by MS are highlighted in orange.

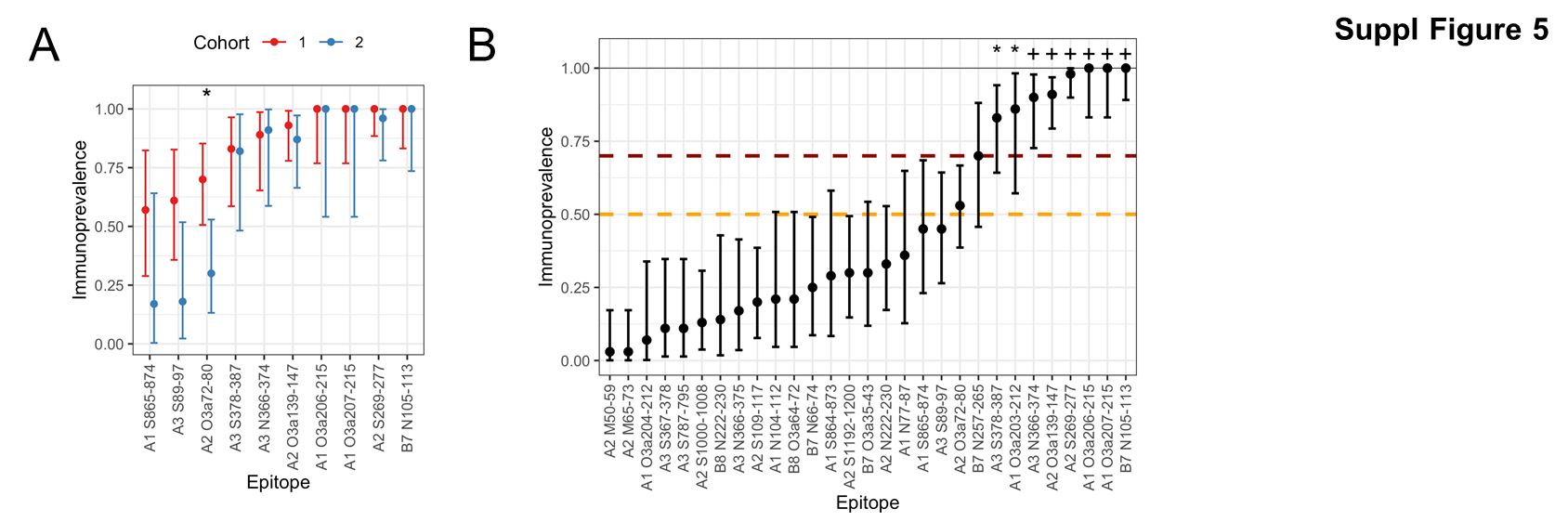

Supplementary Figure 5. Immunoprevalence evaluated by binomial tests

**(A)** Immunoprevalence with exact binomial 95 % confidence intervals estimated separately for the two independent cohorts of convalescent individuals analyzed for multimer responses (Cohort 1 and Cohort 2). Shows all epitopes tested in Cohort 2. Differences in immunoprevalence between the cohorts were assessed using Fisher’s exact test (*: p<0.05). **(B)** Immunoprevalence with exact binomial 95 % confidence intervals for all immunogenic epitopes identified in our study, estimated based on pooled data from cohorts 1 and 2. Binomial tests were performed to assess whether the immunoprevalence for each epitope was significantly higher than 50% (orange line) and 70% (red line), respectively (*: p<0.05 for immunoprevalence > 50%; +: p<0.05 for immunoprevalence > 70%).

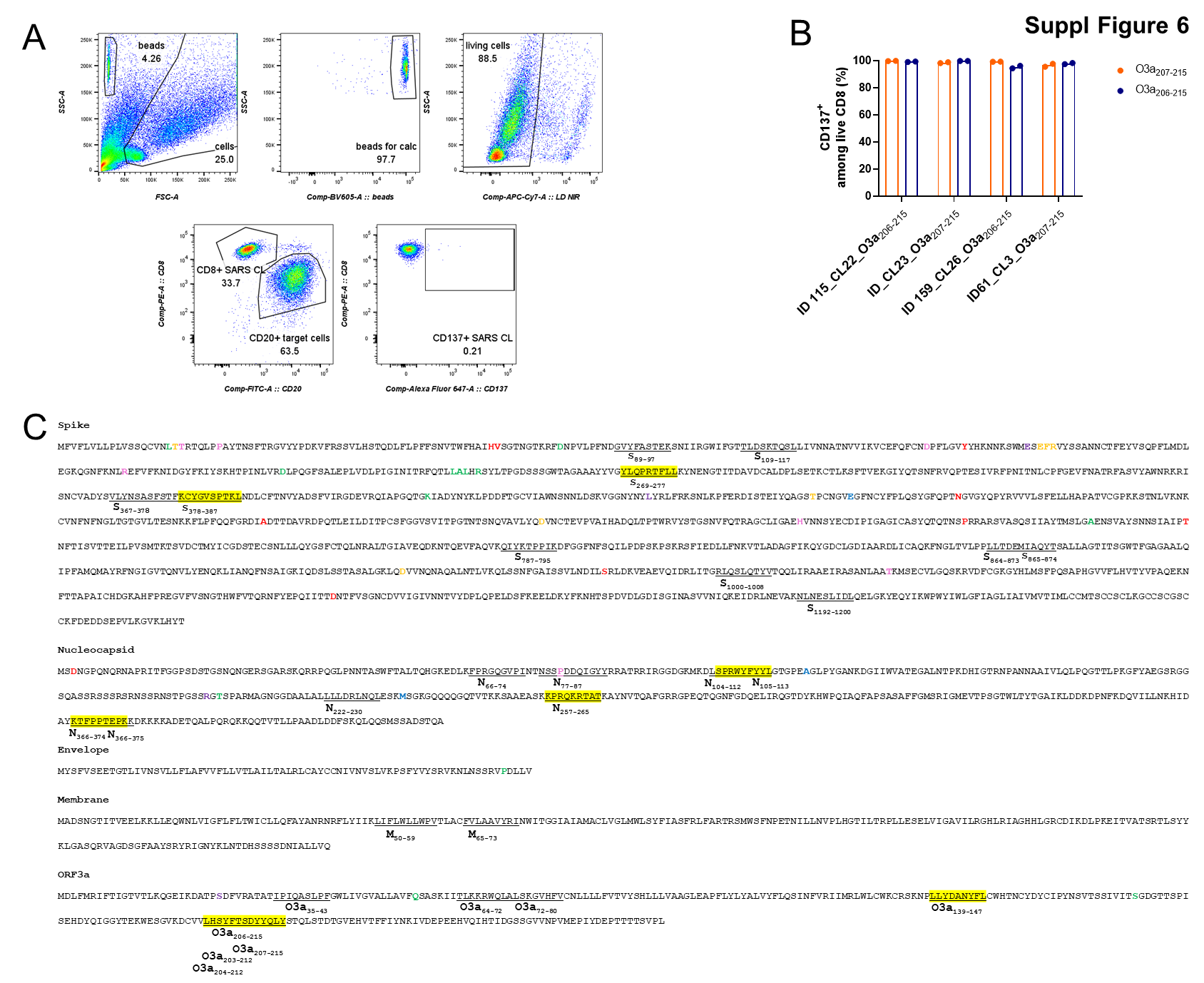

Supplementary Figure 6. Gating strategy for functional validation of T-cell lines (A), cross-recognition of T-cell lines of length variants (B), and T-cell epitope alignment (C)

**(A)** Gating strategy for functional validation of SARS-CoV-2 specific T-cell lines. Here, flow plots for B721.221-HLA-A*02:01 cells w/o antigen co-cultured with a T-cell line recognizing an HLA-A*02:01 specific peptide are shown. Beads (middle top plot) are added only to the samples used to quantitate absolute numbers of events in the target cell gate (lower left plot) to determine killing by CD8 T-cell lines. In the lower right plot the gate used to determine the fraction of activated CD137^+^ T cells is shown.

**(B)** T-cell lines sorted for binding of multimers complexed with ORF3a-derived peptides O3a_207-215_ (FTSDYYQLY; 9-mer) and O3a_206-215_ (YFTSDYYQLY; 10-mer) from several convalescent individuals recognize the 9- and 10-mer equally well, as determined by activation marker expression (CD137) after 20 h co-culture with peptide-loaded (100 nM) mono-allelic B721.221 cells.

**(C)** Alignment of identified 29 immunogenic peptides with the SARS-CoV-2 Wuhan-Hu-1 amino acid sequence (GenBank: MN908947.3). Amino acids mutated in variants of concern (alpha in red, beta in green, gamma in pink) and variants under observation (zeta in blue, kappa in purple) as defined by Public Health England (according to Technical briefings released before July 9, 2021) are highlighted. Mutations found in several strains are highlighted in color of first occurring strain (detailed overview see Suppl Table 7). Sequences recognized by immunoprevalent peptides (70% or more) are highlighted in yellow.

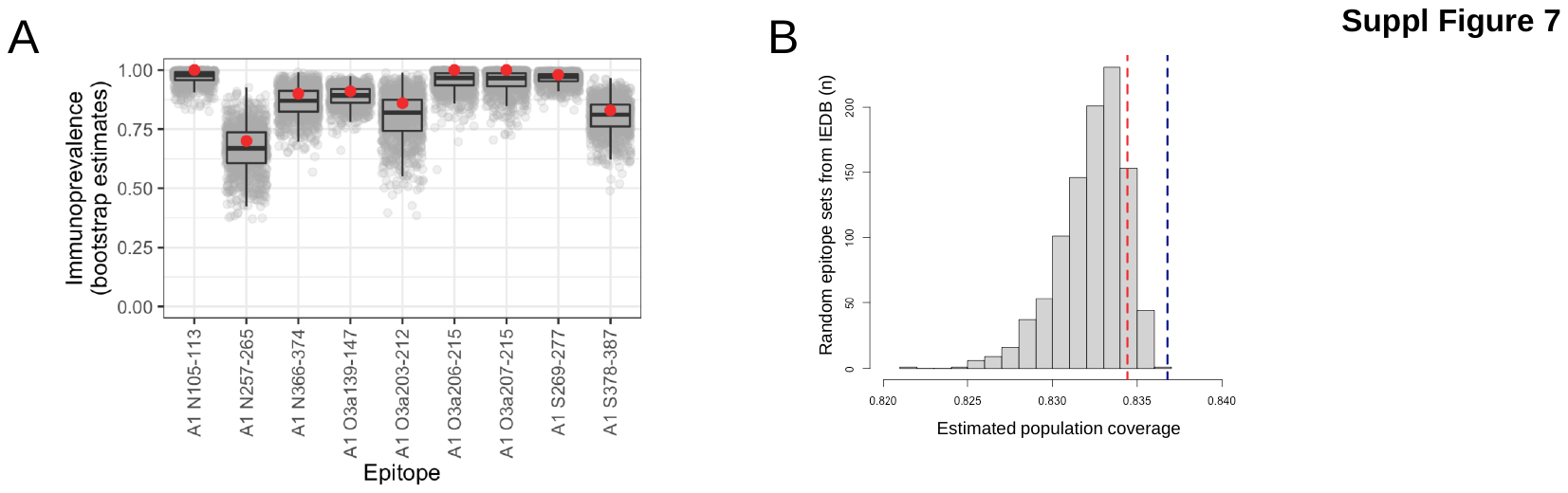

Supplementary Figure 7. Uncertainty in immunoprevalence estimates and population coverage

**(A)** Uncertainty in immunoprevalence estimates for our top 9 immunoprevalent epitopes based on bootstrapping from the beta distribution. Each of 1000 iterations per epitope are represented as shaded gray dots. Boxplots show median and inter-quartile range (IQR); whiskers extend to ± 1.5 times the IQR. The observed immunoprevalence in experimental data is drawn as a red dot. **(B)** Coverage in the European Caucasian population with immunoprevalent peptides identified in this study. The dashed vertical blue line represents population coverage with peptides recognizing HLA-A*01:01, HLA-A*02:01, HLA-A*03:01 and HLA-B*07:02, based solely on HLA allele frequencies (not considering immunoprevalence). Estimated population coverage with our top 9 epitopes (prevalence 70% or higher) after adjusting for immunoprevalence is shown as a dashed vertical red line. Histogram in gray shows population coverage for the same 9 epitopes when considering uncertainty in immunoprevalence estimates, as represented by bootstrapped data shown in **(A).**
