## Supplementary File for "Beyond Spike: Identification of nine highly prevalent SARS-CoV-2-specific CD8 T-cell epitopes in a large Norwegian cohort"

**Supplementary file 1:** MS/MS fragmentation spectra of synthetic and eluted HLA class I peptides identified from mono-allelic B721.221 cells expressing the SARS-CoV-2 proteins. 49 out of the 50 peptides reported in the study were validated with the corresponding synthetic peptides. Peptides reported as immunogenic in the current study are highlighted in yellow background.

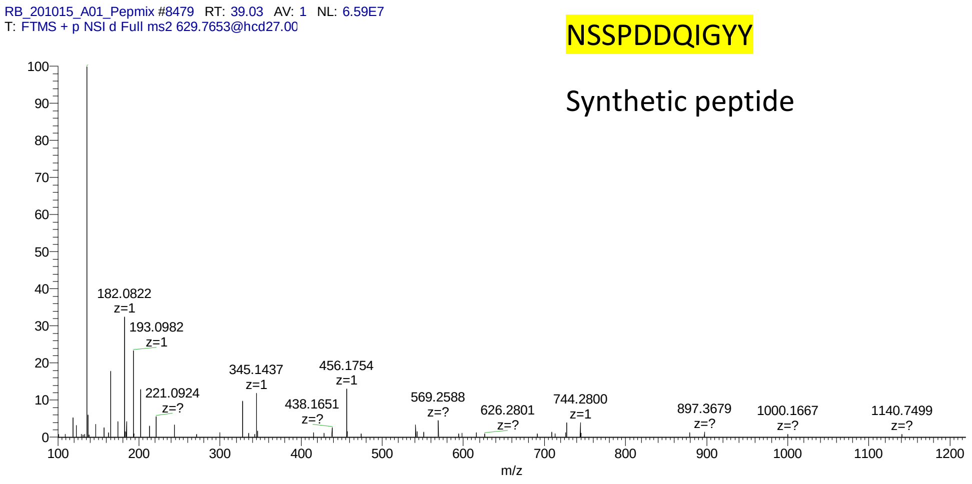


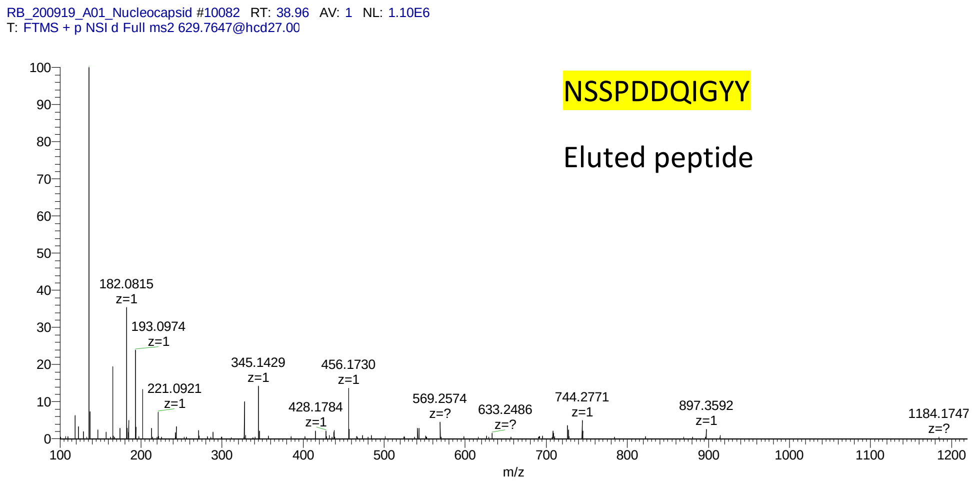

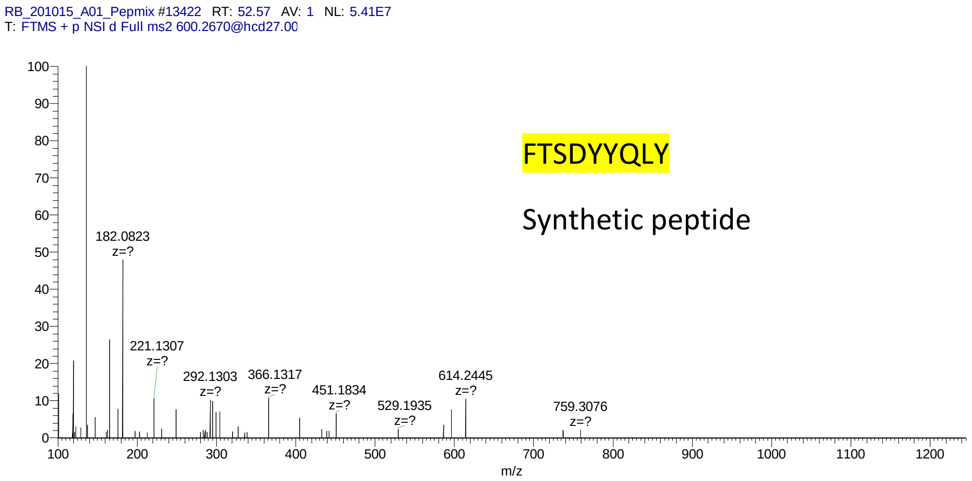
RB_201015_A01_Pepmix #13422 RT: 52.57 AV: 1 NL: 5.41E7
T: FTMS + p NSI d Full ms2 600.2670@hcd27.00
FTSDYYQLY
Synthetic peptide
182.0823 z=?
221.1307 z=?
292.1303 z=?
366.1317 z=?
451.1834 z=?
529.1935 z=?
614.2445 z=?
759.3076 z=?
m/z

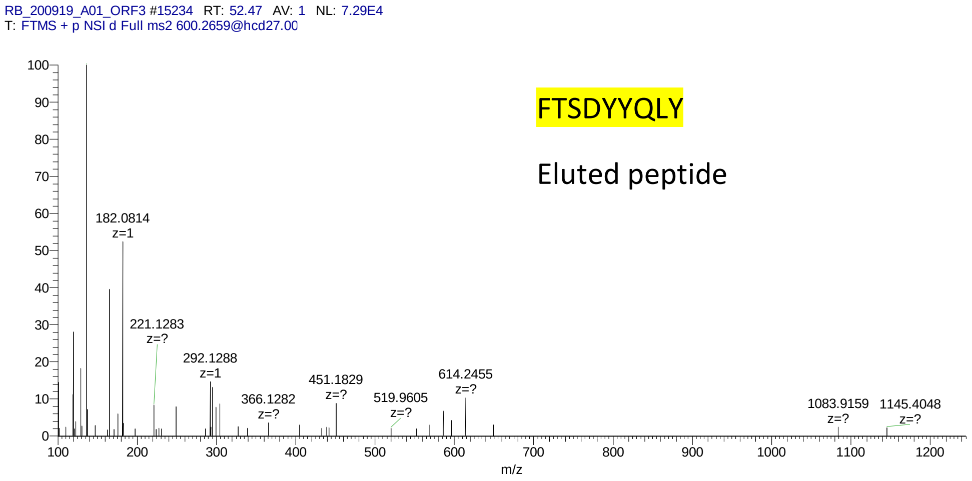
RB_200919_A01_ORF3 #15234 RT: 52.47 AV: 1 NL: 7.29E4
T: FTMS + p NSI d Full ms2 600.2659@hcd27.00
FTSDYYQLY
Eluted peptide
182.0814 z=1
221.1283 z=?
292.1288 z=1
366.1282 z=?
451.1829 z=?
519.9605 z=?
614.2455 z=?
1083.9159 z=?
1145.4048 z=?
m/z

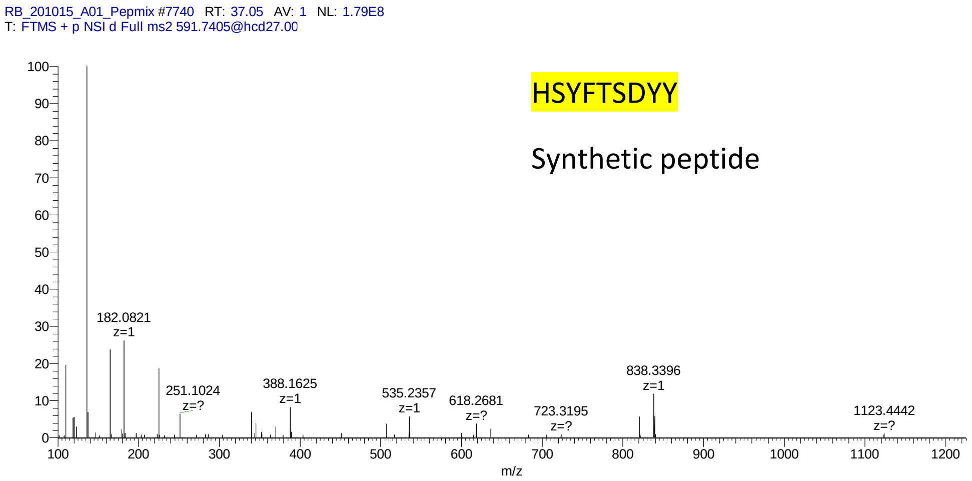
RB_201015_A01_Pepmix #7740 RT: 37.05 AV: 1 NL: 1.79E8
T: FTMS + p NSI d Full ms2 591.7405@hcd27.00
HSYFTSDYY
Synthetic peptide
182.0821 z=1
251.1024 z=?
388.1625 z=1
535.2357 z=1
618.2681 z=?
723.3195 z=?
838.3396 z=1
1123.4442 z=?
m/z

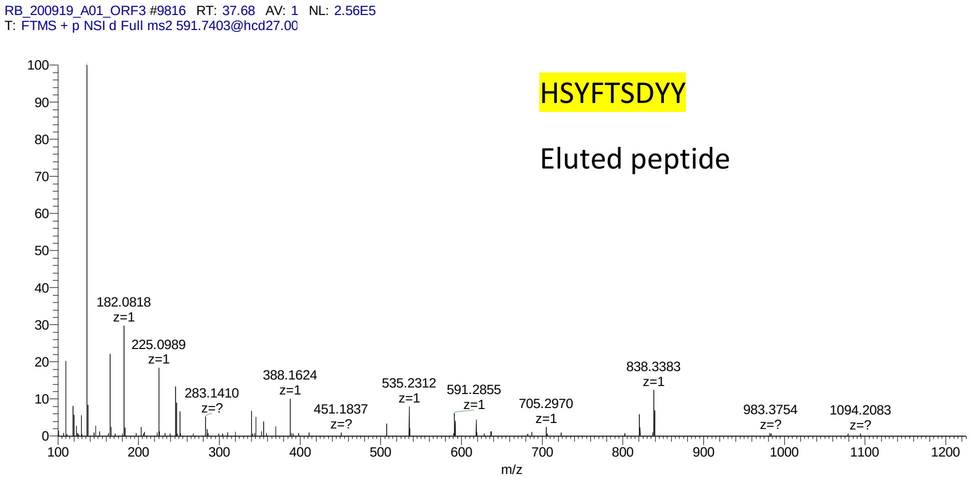
RB_200919_A01_ORF3 #9816 RT: 37.68 AV: 1 NL: 2.56E5
T: FTMS + p NSI d Full ms2 591.7403@hcd27.00
HSYFTSDYY
Eluted peptide
182.0818 z=1
225.0989 z=1
283.1410 z=?
388.1624 z=1
451.1837 z=?
535.2312 z=1
591.2855 z=1
705.2970 z=1
838.3383 z=1
983.3754 z=?
1094.2083 z=?
m/z

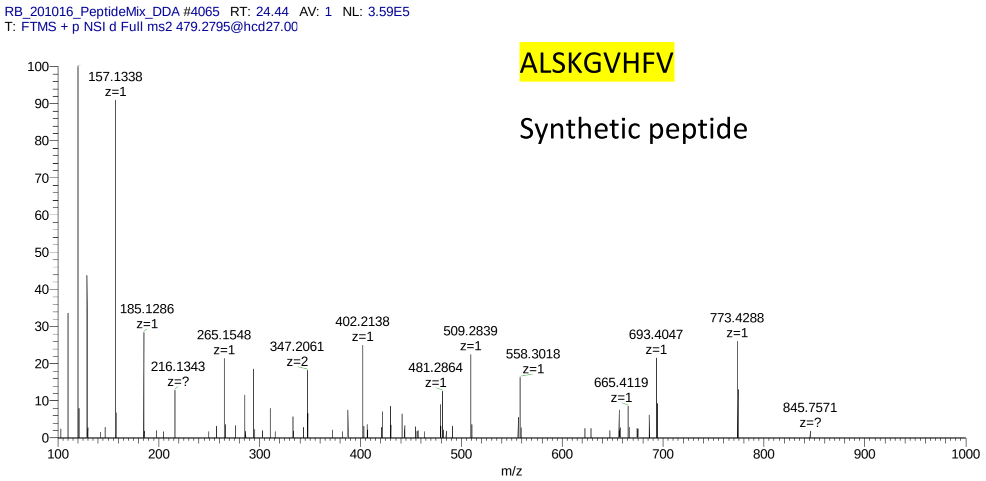
RB_201016_PeptideMix_DDA #4065 RT: 24.44 AV: 1 NL: 3.59E5
T: FTMS + p NSI d Full ms2 479.2795@hcd27.00
ALSKGVHFV
Synthetic peptide
157.1338 z=1
185.1286 z=1
216.1343 z=?
265.1548 z=1
347.2061 z=2
402.2138 z=1
481.2864 z=1
509.2839 z=1
558.3018 z=1
665.4119 z=1
693.4047 z=1
773.4288 z=1
845.7571 z=?
m/z

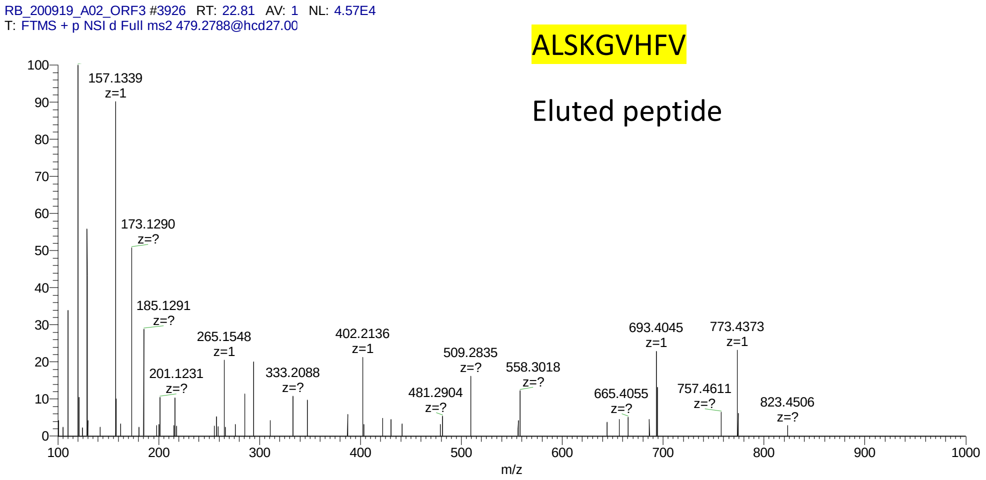
RB_200919_A02_ORF3 #3926 RT: 22.81 AV: 1 NL: 4.57E4
T: FTMS + p NSI d Full ms2 479.2788@hcd27.00
ALSKGVHFV
Eluted peptide
157.1339 z=1
173.1290 z=?
185.1291 z=?
201.1231 z=?
265.1548 z=1
333.2088 z=?
402.2136 z=1
481.2904 z=?
509.2835 z=?
558.3018 z=?
665.4055 z=?
693.4045 z=1
757.4611 z=?
773.4373 z=1
823.4506 z=?
m/z

RB_201016_PeptideMix_DDA #13724 RT: 49.70 AV: 1 NL: 2.61E5
T: FTMS + p NSI d Full ms2 549.3371@hcd27.00 [100.0000-1140.0000]

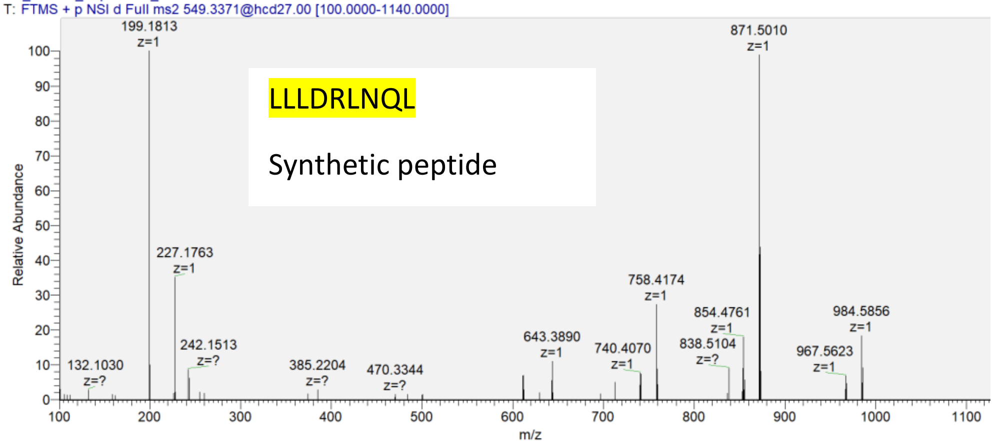


RB_200919_A02_Nucleocapsid #13809 RT: 49.75 AV: 1 NL: 1.54E5
T: FTMS + p NSI d Full ms2 549.3374@hcd27.00

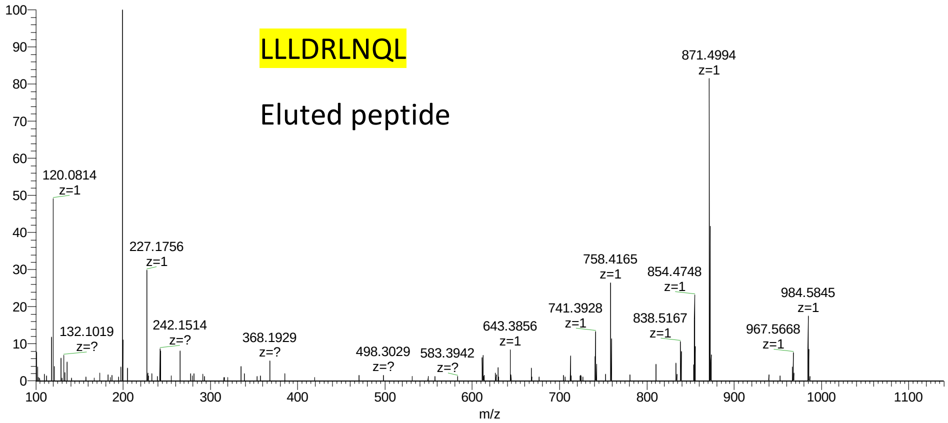

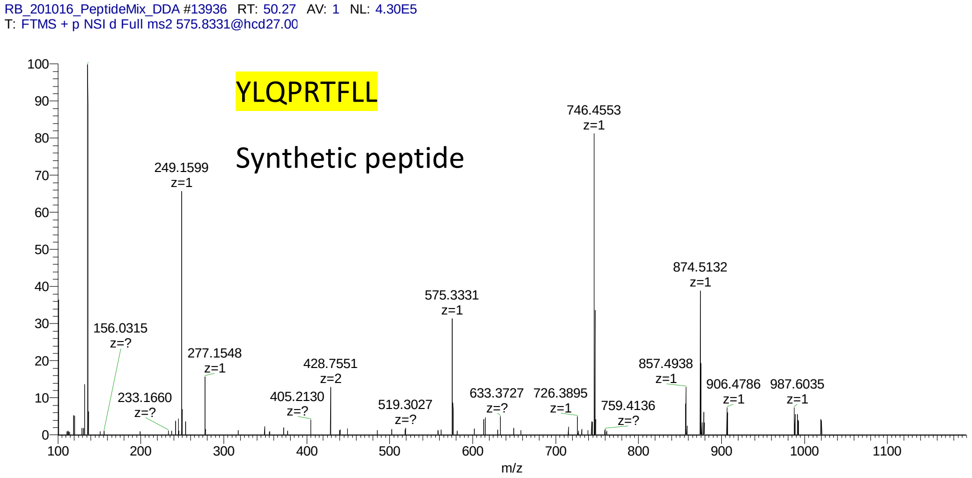
RB_201016_PeptideMix_DDA #13936 RT: 50.27 AV: 1 NL: 4.30E5
T: FTMS + p NSI d Full ms2 575.8331@hcd27.00
YLQPRTFLL
Synthetic peptide
746.4553 z=1
249.1599 z=1
874.5132 z=1
575.3331 z=1
156.0315 z=?
277.1548 z=1
428.7551 z=2
857.4938 z=1
233.1660 z=?
405.2130 z=?
519.3027 z=?
633.3727 z=?
726.3895 z=1
759.4136 z=?
906.4786 z=1
987.6035 z=1
m/z

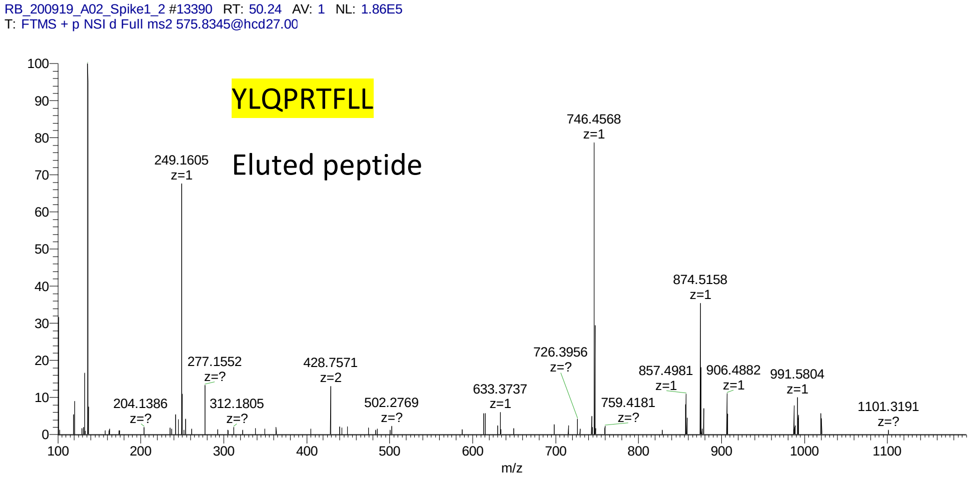
RB_200919_A02_Spike1_2 #13390 RT: 50.24 AV: 1 NL: 1.86E5
T: FTMS + p NSI d Full ms2 575.8345@hcd27.00
YLQPRTFLL
Eluted peptide
746.4568 z=1
249.1605 z=1
874.5158 z=1
277.1552 z=?
428.7571 z=2
726.3956 z=?
204.1386 z=?
312.1805 z=?
502.2769 z=?
633.3737 z=1
759.4181 z=?
857.4981 z=1
906.4882 z=1
991.5804 z=1
1101.3191 z=?
m/z

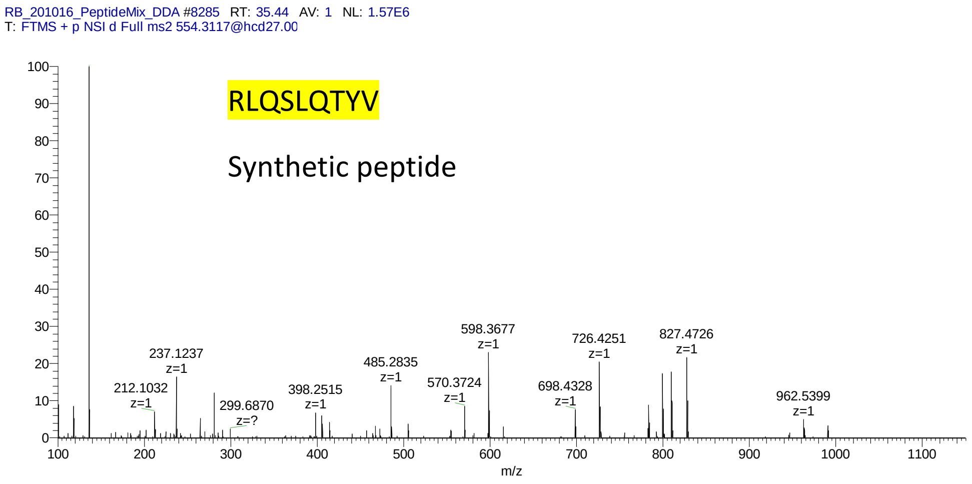
RB_201016_PeptideMix_DDA #8285 RT: 35.44 AV: 1 NL: 1.57E6
T: FTMS + p NSI d Full ms2 554.3117@hcd27.00
RLQSLQTYV
Synthetic peptide
212.1032 z=1
237.1237 z=1
299.6870 z=?
398.2515 z=1
485.2835 z=1
570.3724 z=1
598.3677 z=1
698.4328 z=1
726.4251 z=1
827.4726 z=1
962.5399 z=1
m/z

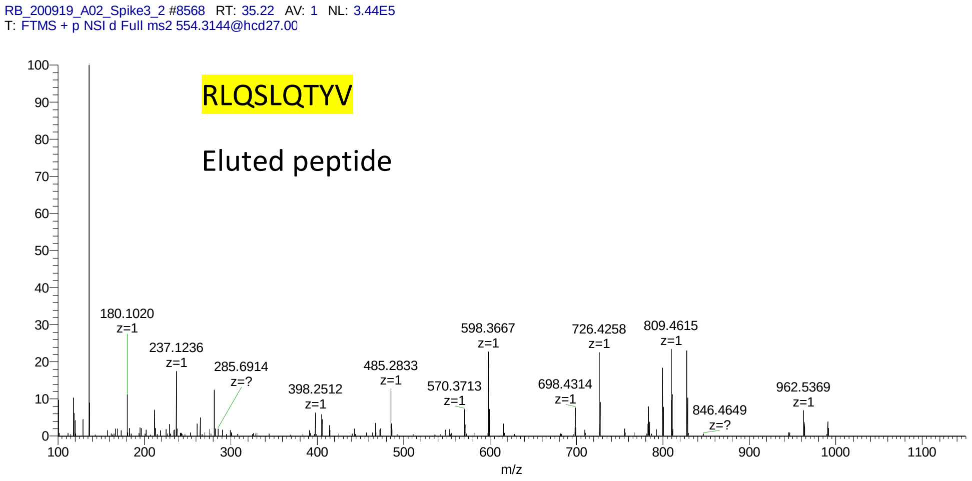
RB_200919_A02_Spike3_2 #8568 RT: 35.22 AV: 1 NL: 3.44E5
T: FTMS + p NSI d Full ms2 554.3144@hcd27.00
RLQSLQTYV
Eluted peptide
180.1020 z=1
237.1236 z=1
285.6914 z=?
398.2512 z=1
485.2833 z=1
570.3713 z=1
598.3667 z=1
698.4314 z=1
726.4258 z=1
809.4615 z=1
846.4649 z=?
962.5369 z=1
m/z

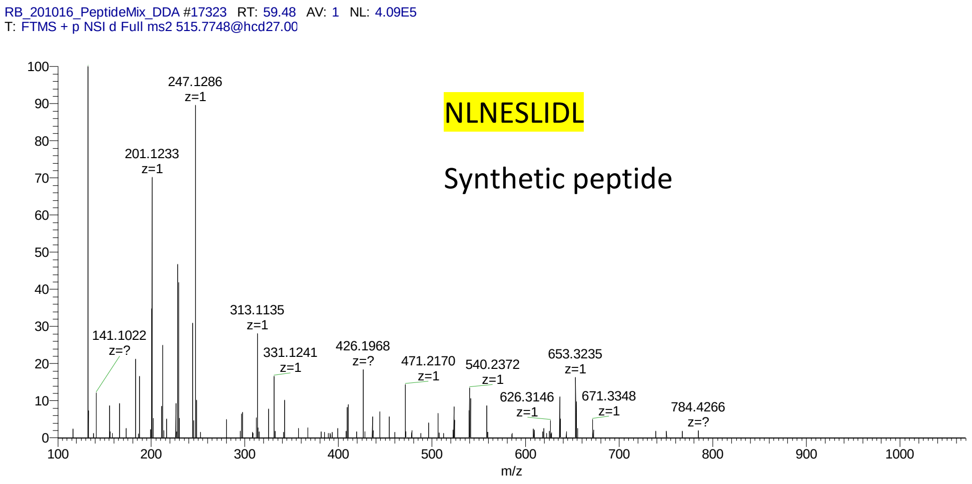
RB_201016_PeptideMix_DDA #17323 RT: 59.48 AV: 1 NL: 4.09E5
T: FTMS + p NSI d Full ms2 515.7748@hcd27.00
NLNESLIDL
Synthetic peptide
247.1286 z=1
201.1233 z=1
141.1022 z=?
313.1135 z=1
331.1241 z=1
426.1968 z=?
471.2170 z=1
540.2372 z=1
626.3146 z=1
653.3235 z=1
671.3348 z=1
784.4266 z=?
m/z

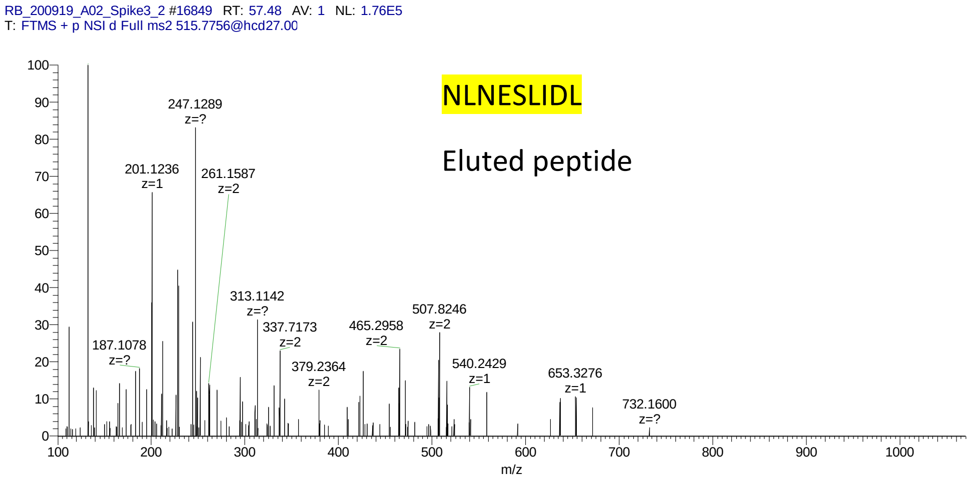
RB_200919_A02_Spike3_2 #16849 RT: 57.48 AV: 1 NL: 1.76E5
T: FTMS + p NSI d Full ms2 515.7756@hcd27.00
NLNESLIDL
Eluted peptide
247.1289 z=?
201.1236 z=1
261.1587 z=2
187.1078 z=?
313.1142 z=?
337.7173 z=2
379.2364 z=2
465.2958 z=2
507.8246 z=2
540.2429 z=1
653.3276 z=1
732.1600 z=?
m/z

RB_201016_PeptideMix_DDA #1213 RT: 16.06 AV: 1 NL: 3.44E5
T: FTMS + p NSI d Full ms2 522.8040@hcd27.00

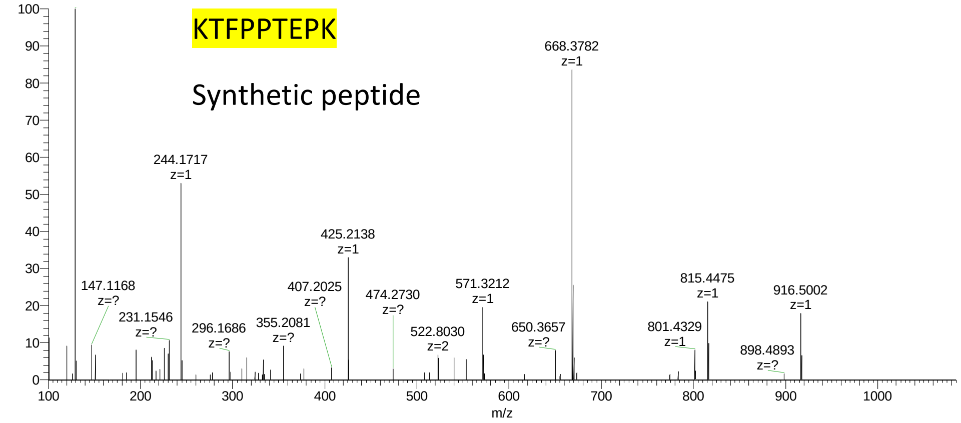


RB_200919_A03_Nucleocapsid #2845 RT: 19.05 AV: 1 NL: 9.73E4
T: FTMS + p NSI d Full ms2 522.7906@hcd27.00

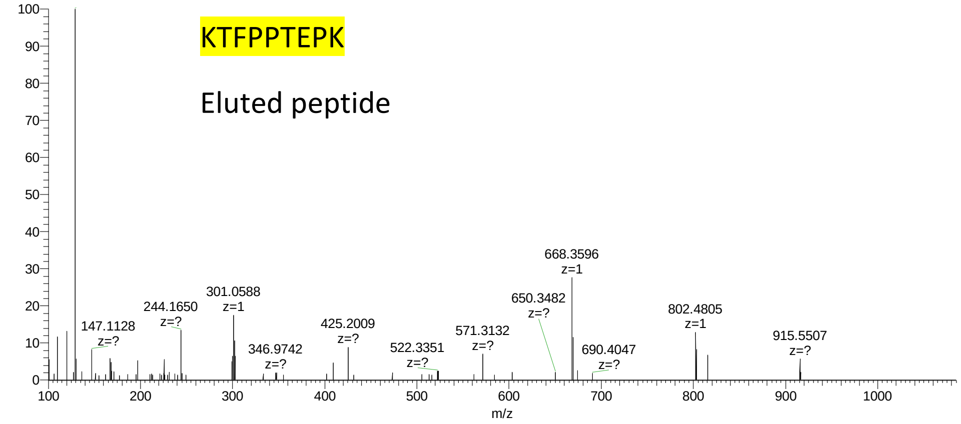

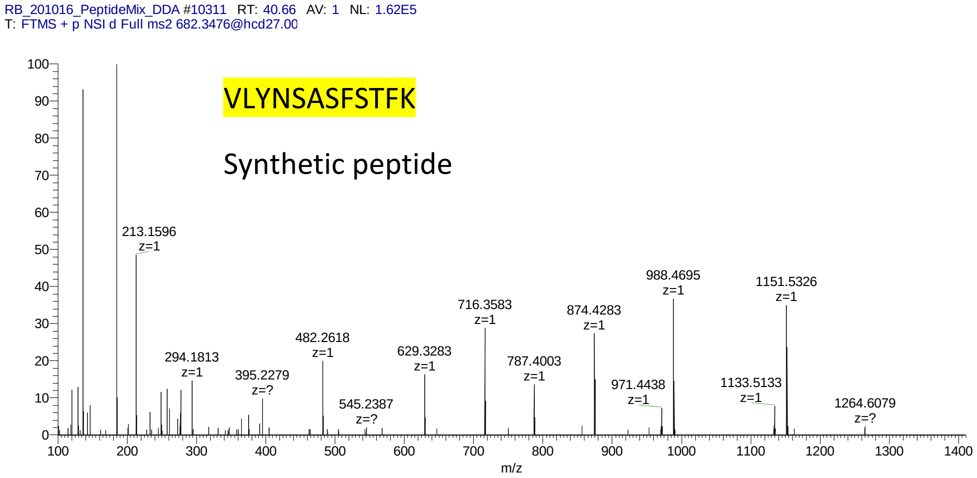
RB_201016_PeptideMix_DDA #10311 RT: 40.66 AV: 1 NL: 1.62E5
T: FTMS + p NSI d Full ms2 682.3476@hcd27.00
VLYNSASFSTFK
Synthetic peptide
213.1596 z=1
294.1813 z=1
395.2279 z=?
482.2618 z=1
545.2387 z=?
629.3283 z=1
716.3583 z=1
787.4003 z=1
874.4283 z=1
971.4438 z=1
988.4695 z=1
1133.5133 z=1
1151.5326 z=1
1264.6079 z=?
m/z

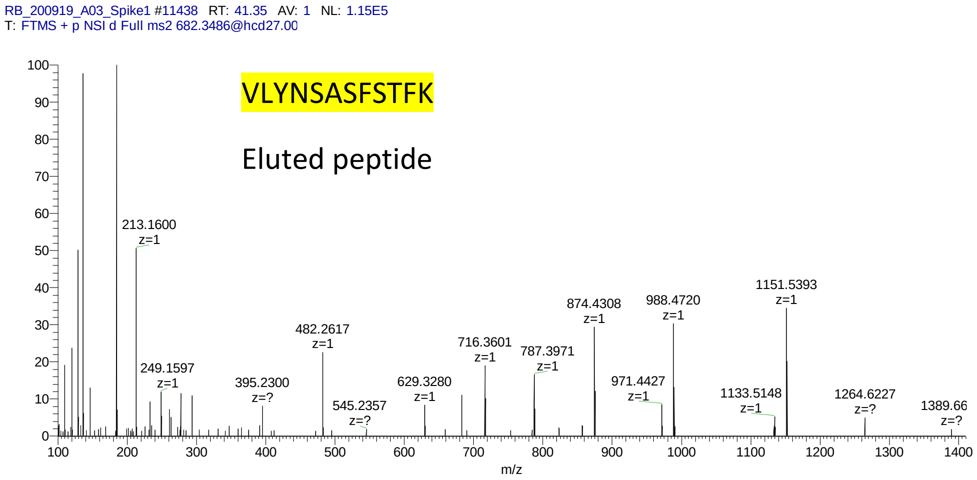
RB_200919_A03_Spike1 #11438 RT: 41.35 AV: 1 NL: 1.15E5
T: FTMS + p NSI d Full ms2 682.3486@hcd27.00
VLYNSASFSTFK
Eluted peptide
213.1600 z=1
249.1597 z=1
395.2300 z=?
482.2617 z=1
545.2357 z=?
629.3280 z=1
716.3601 z=1
787.3971 z=1
874.4308 z=1
971.4427 z=1
988.4720 z=1
1133.5148 z=1
1151.5393 z=1
1264.6227 z=?
1389.66 z=?
m/z

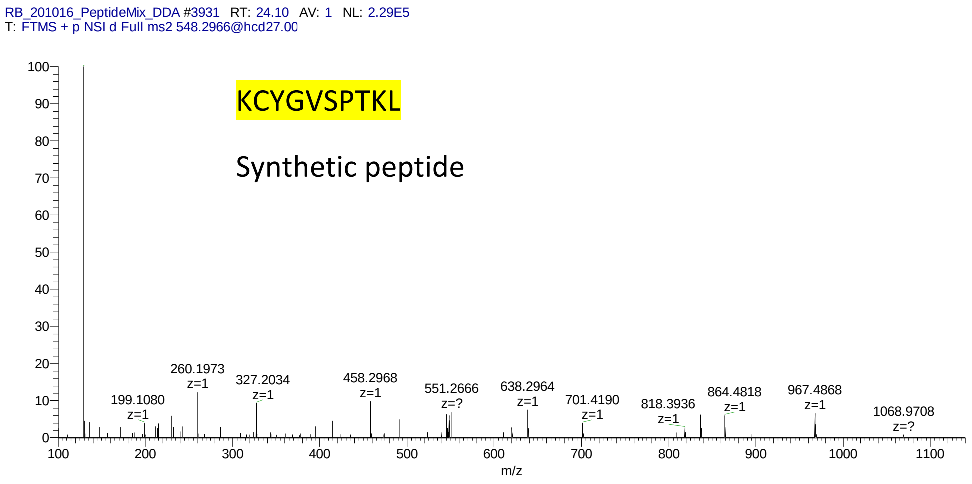
RB_201016_PeptideMix_DDA #3931 RT: 24.10 AV: 1 NL: 2.29E5
T: FTMS + p NSI d Full ms2 548.2966@hcd27.00
KCYGVSPTKL
Synthetic peptide
199.1080 z=1
260.1973 z=1
327.2034 z=1
458.2968 z=1
551.2666 z=?
638.2964 z=1
701.4190 z=1
818.3936 z=1
864.4818 z=1
967.4868 z=1
1068.9708 z=?
m/z

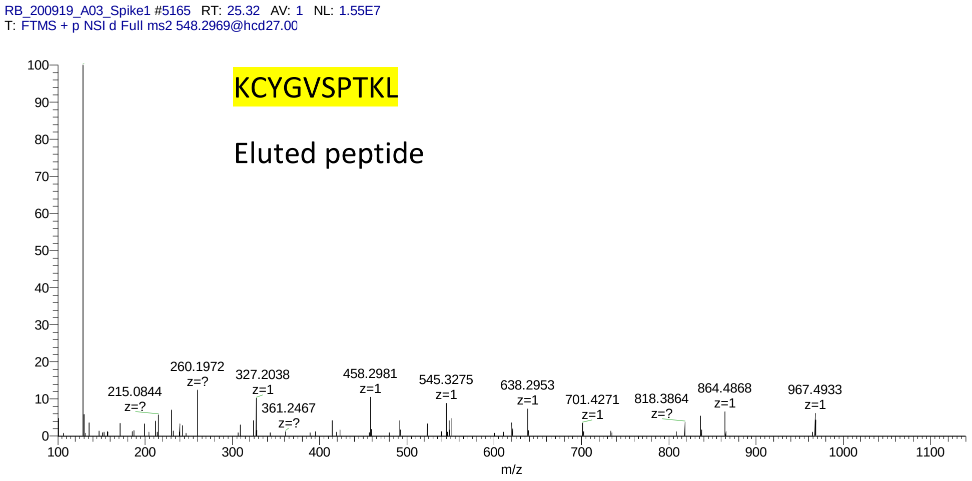
RB_200919_A03_Spike1 #5165 RT: 25.32 AV: 1 NL: 1.55E7
T: FTMS + p NSI d Full ms2 548.2969@hcd27.00
KCYGVSPTKL
Eluted peptide
215.0844 z=?
260.1972 z=?
327.2038 z=1
361.2467 z=?
458.2981 z=1
545.3275 z=1
638.2953 z=1
701.4271 z=1
818.3864 z=?
864.4868 z=1
967.4933 z=1
m/z

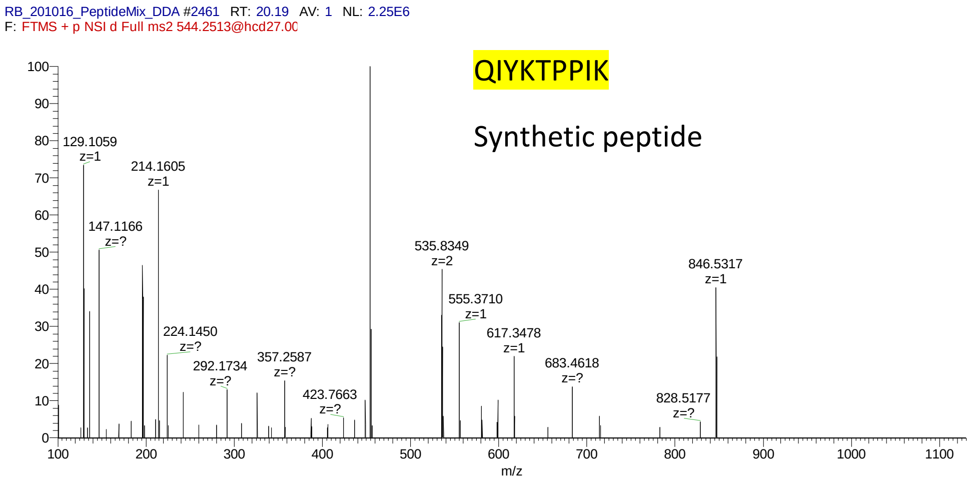
RB_201016_PeptideMix_DDA #2461 RT: 20.19 AV: 1 NL: 2.25E6
F: FTMS + p NSI d Full ms2 544.2513@hcd27.00
QIYKTPPIK
Synthetic peptide
129.1059 z=1
214.1605 z=1
147.1166 z=?
224.1450 z=?
292.1734 z=?
357.2587 z=?
423.7663 z=?
535.8349 z=2
555.3710 z=1
617.3478 z=1
683.4618 z=?
828.5177 z=?
846.5317 z=1
m/z

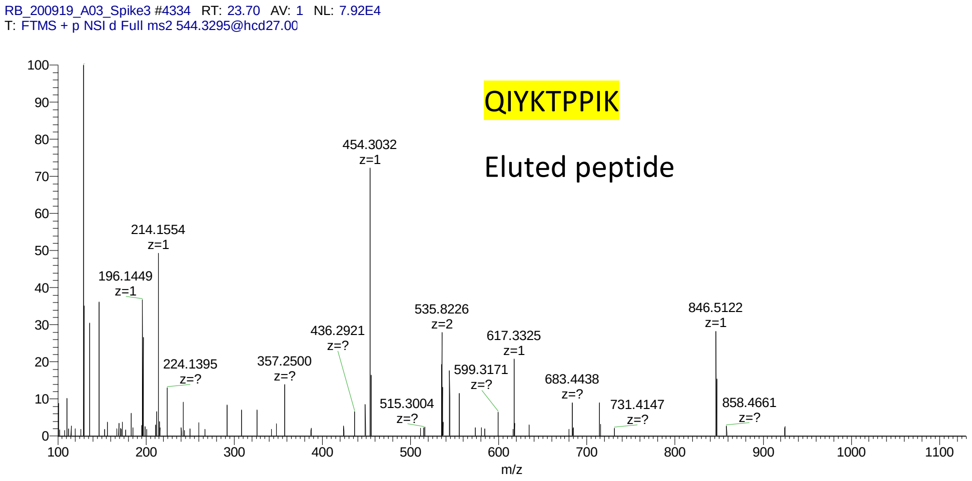
RB_200919_A03_Spike3 #4334 RT: 23.70 AV: 1 NL: 7.92E4
T: FTMS + p NSI d Full ms2 544.3295@hcd27.00
QIYKTPPIK
Eluted peptide
454.3032 z=1
214.1554 z=1
196.1449 z=1
224.1395 z=?
357.2500 z=?
436.2921 z=?
515.3004 z=?
535.8226 z=2
599.3171 z=?
617.3325 z=1
683.4438 z=?
731.4147 z=?
846.5122 z=1
858.4661 z=?
m/z

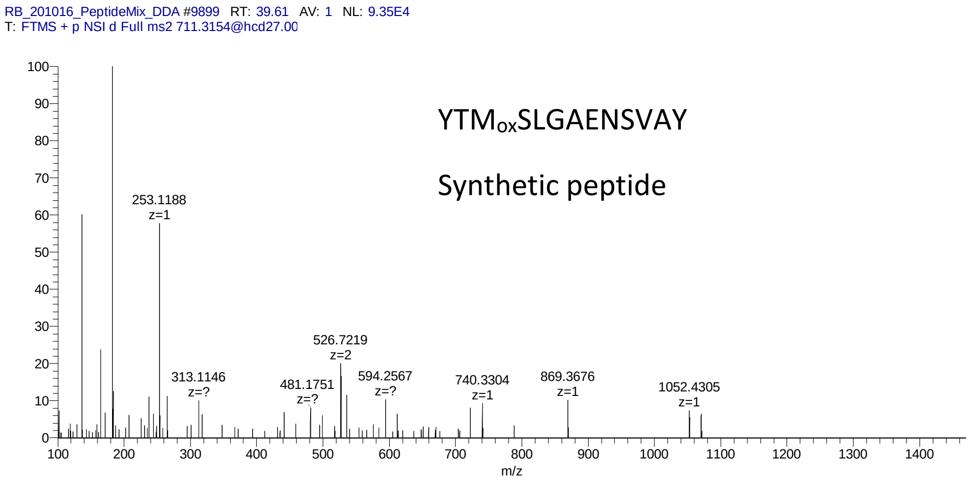
RB_201016_PeptideMix_DDA #9899 RT: 39.61 AV: 1 NL: 9.35E4
T: FTMS + p NSI d Full ms2 711.3154@hcd27.00
YTM$_{ox}$SLGAENSVAY
Synthetic peptide
253.1188 z=1
313.1146 z=?
481.1751 z=?
526.7219 z=2
594.2567 z=?
740.3304 z=1
869.3676 z=1
1052.4305 z=1
m/z

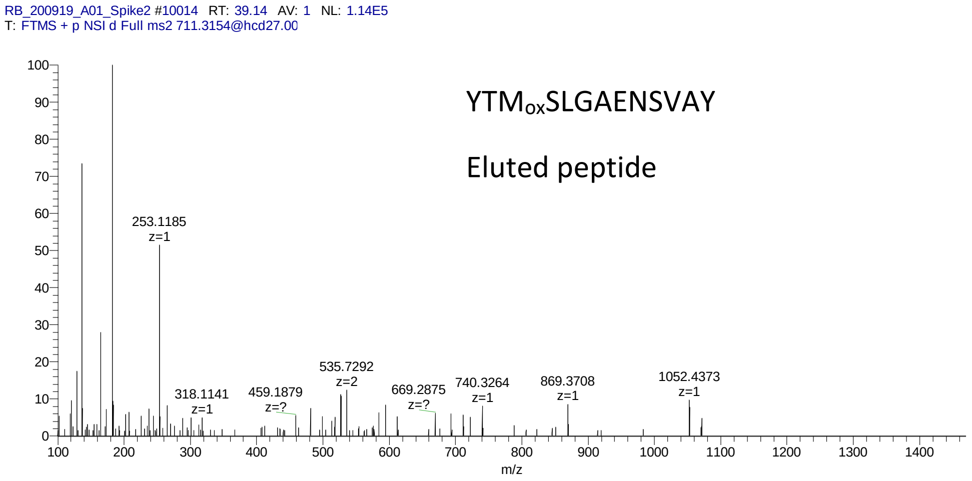
RB_200919_A01_Spike2 #10014 RT: 39.14 AV: 1 NL: 1.14E5
T: FTMS + p NSI d Full ms2 711.3154@hcd27.00
YTM$_{ox}$SLGAENSVAY
Eluted peptide
253.1185 z=1
318.1141 z=1
459.1879 z=?
535.7292 z=2
669.2875 z=?
740.3264 z=1
869.3708 z=1
1052.4373 z=1
m/z

RB_201016_PeptideMix_DDA #9162 RT: 37.70 AV: 1 NL: 3.73E5
T: FTMS + p NSI d Full ms2 550.2524@hcd27.00

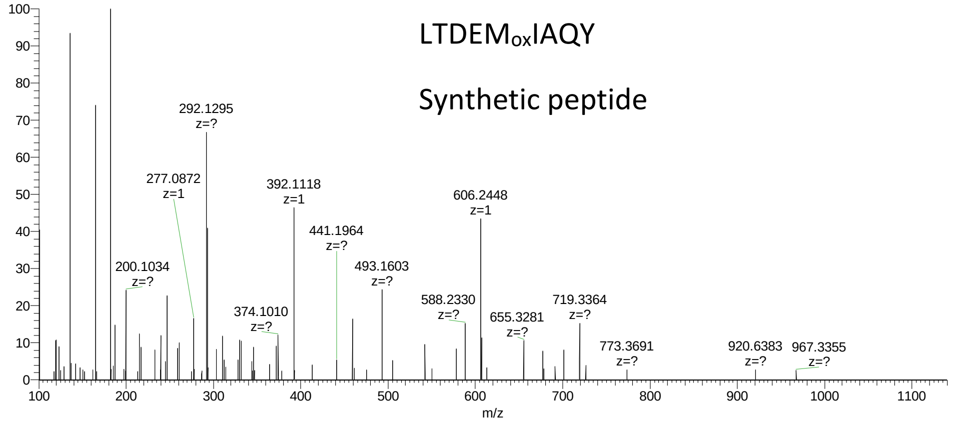


RB_200919_A01_Spike2 #9400 RT: 37.50 AV: 1 NL: 1.33E6
T: FTMS + p NSI d Full ms2 550.2529@hcd27.00

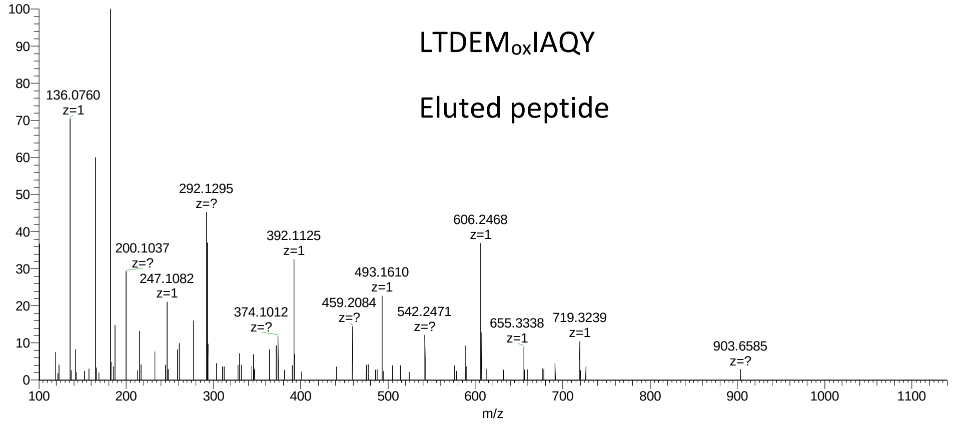

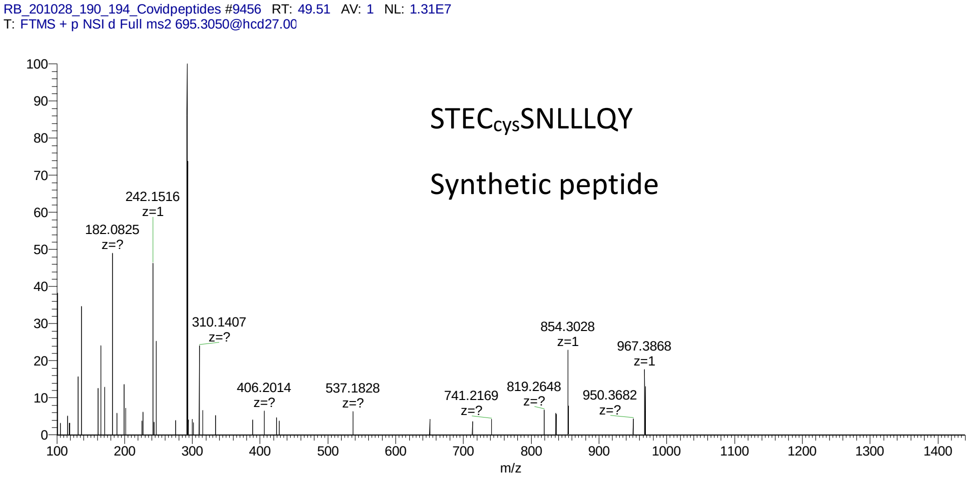
RB_201028_190_194_Covidpeptides #9456 RT: 49.51 AV: 1 NL: 1.31E7
T: FTMS + p NSI d Full ms2 695.3050@hcd27.00
STECcysSNLLLQY
Synthetic peptide
182.0825 z=?
242.1516 z=1
310.1407 z=?
406.2014 z=?
537.1828 z=?
741.2169 z=?
819.2648 z=?
854.3028 z=1
950.3682 z=?
967.3868 z=1
m/z

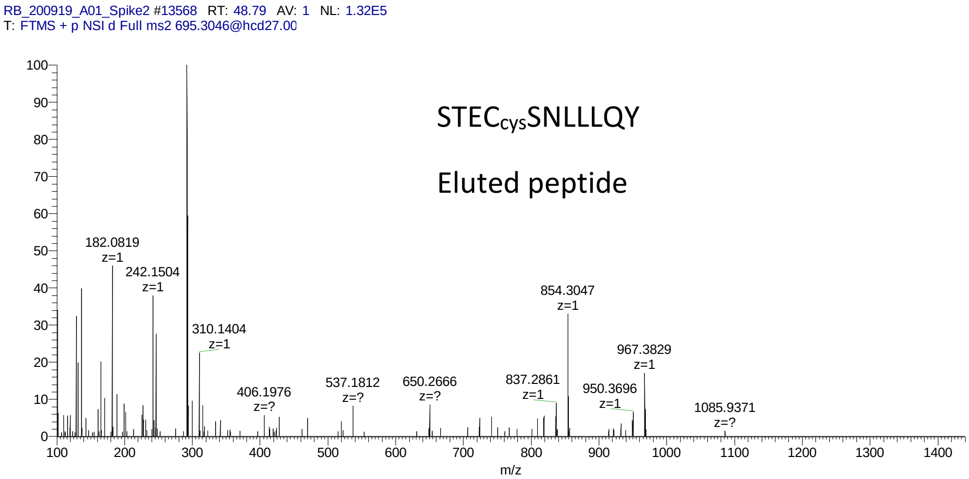
RB_200919_A01_Spike2 #13568 RT: 48.79 AV: 1 NL: 1.32E5
T: FTMS + p NSI d Full ms2 695.3046@hcd27.00
STECcysSNLLLQY
Eluted peptide
182.0819 z=1
242.1504 z=1
310.1404 z=1
406.1976 z=?
537.1812 z=?
650.2666 z=?
837.2861 z=1
854.3047 z=1
950.3696 z=1
967.3829 z=1
1085.9371 z=?
m/z

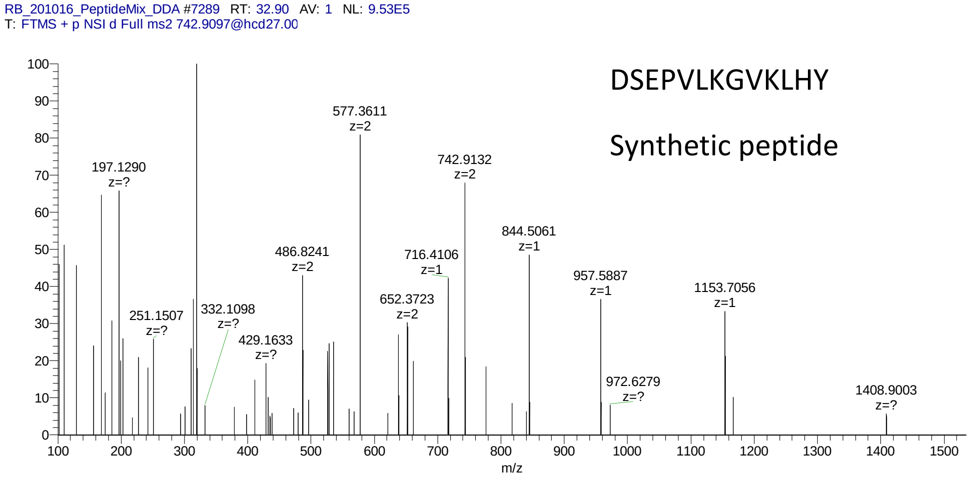
RB_201016_PeptideMix_DDA #7289 RT: 32.90 AV: 1 NL: 9.53E5
T: FTMS + p NSI d Full ms2 742.9097@hcd27.00
DSEPVLKGVKLHY
Synthetic peptide
577.3611 z=2
197.1290 z=?
742.9132 z=2
844.5061 z=1
486.8241 z=2
716.4106 z=1
957.5887 z=1
1153.7056 z=1
251.1507 z=?
332.1098 z=?
652.3723 z=2
429.1633 z=?
972.6279 z=?
1408.9003 z=?
m/z

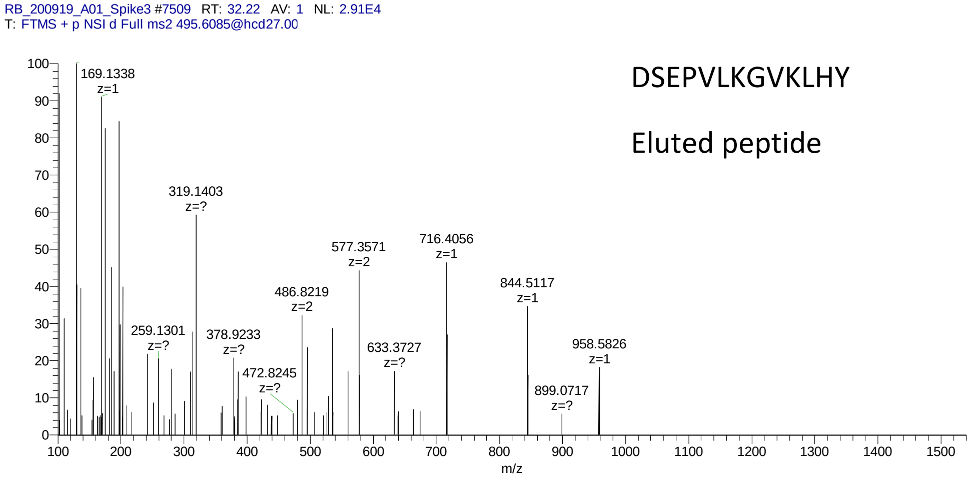
RB_200919_A01_Spike3 #7509 RT: 32.22 AV: 1 NL: 2.91E4
T: FTMS + p NSI d Full ms2 495.6085@hcd27.00
DSEPVLKGVKLHY
Eluted peptide
169.1338 z=1
319.1403 z=?
577.3571 z=2
716.4056 z=1
844.5117 z=1
486.8219 z=2
259.1301 z=?
378.9233 z=?
633.3727 z=?
958.5826 z=1
472.8245 z=?
899.0717 z=?
m/z

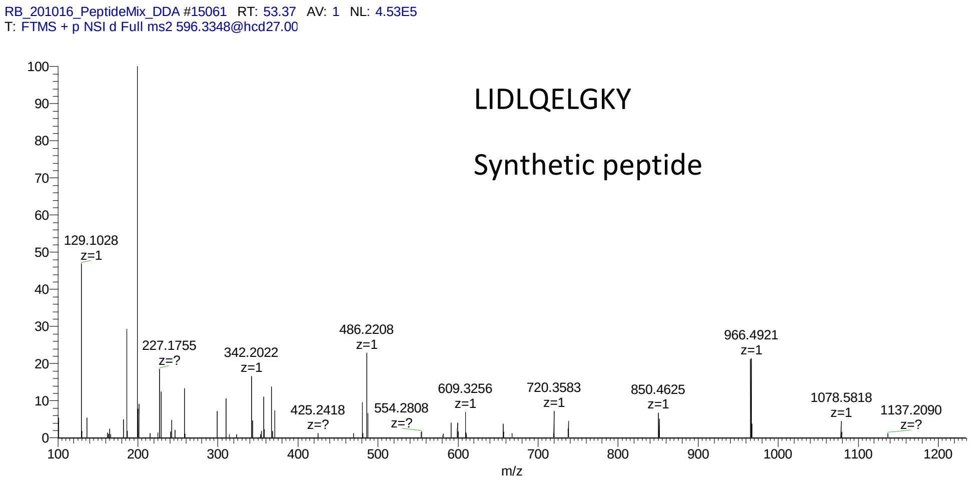
RB_201016_PeptideMix_DDA #15061 RT: 53.37 AV: 1 NL: 4.53E5
T: FTMS + p NSI d Full ms2 596.3348@hcd27.00
LIDLQELGKY
Synthetic peptide
129.1028 z=1
227.1755 z=?
342.2022 z=1
425.2418 z=?
486.2208 z=1
554.2808 z=?
609.3256 z=1
720.3583 z=1
850.4625 z=1
966.4921 z=1
1078.5818 z=1
1137.2090 z=?
m/z

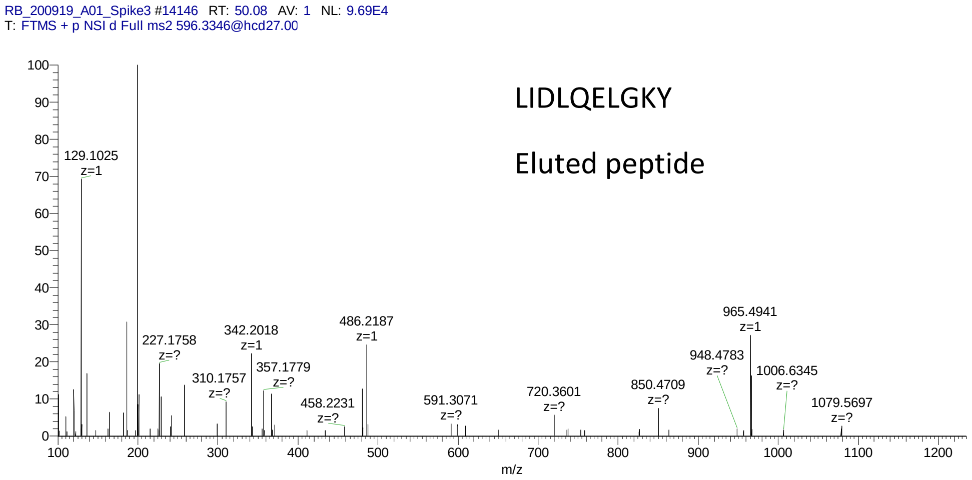
RB_200919_A01_Spike3 #14146 RT: 50.08 AV: 1 NL: 9.69E4
T: FTMS + p NSI d Full ms2 596.3346@hcd27.00
LIDLQELGKY
Eluted peptide
129.1025 z=1
227.1758 z=?
310.1757 z=?
342.2018 z=1
357.1779 z=?
458.2231 z=?
486.2187 z=1
591.3071 z=?
720.3601 z=?
850.4709 z=?
948.4783 z=?
965.4941 z=1
1006.6345 z=?
1079.5697 z=?
m/z

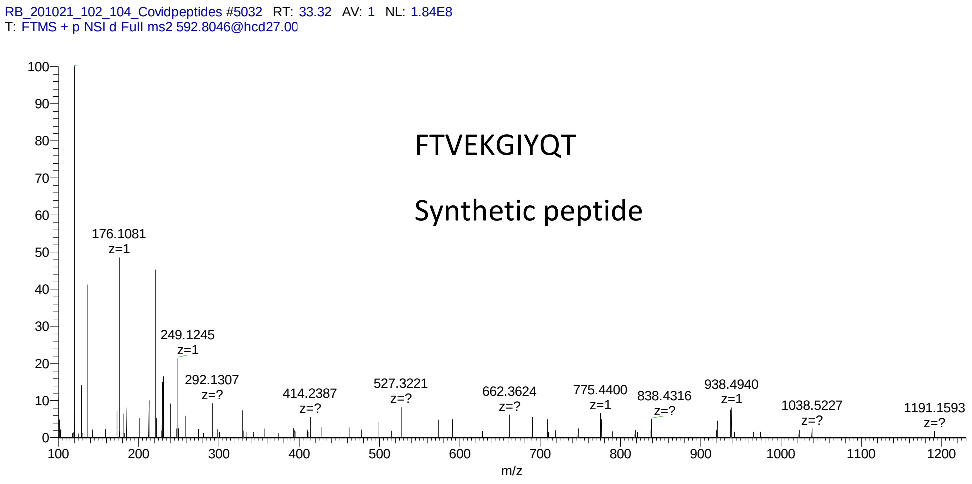
RB_201021_102_104_Covidpeptides #5032 RT: 33.32 AV: 1 NL: 1.84E8
T: FTMS + p NSI d Full ms2 592.8046@hcd27.00
FTVEKGIYQT
Synthetic peptide
176.1081 z=1
249.1245 z=1
292.1307 z=?
414.2387 z=?
527.3221 z=?
662.3624 z=?
775.4400 z=1
838.4316 z=?
938.4940 z=1
1038.5227 z=?
1191.1593 z=?
m/z

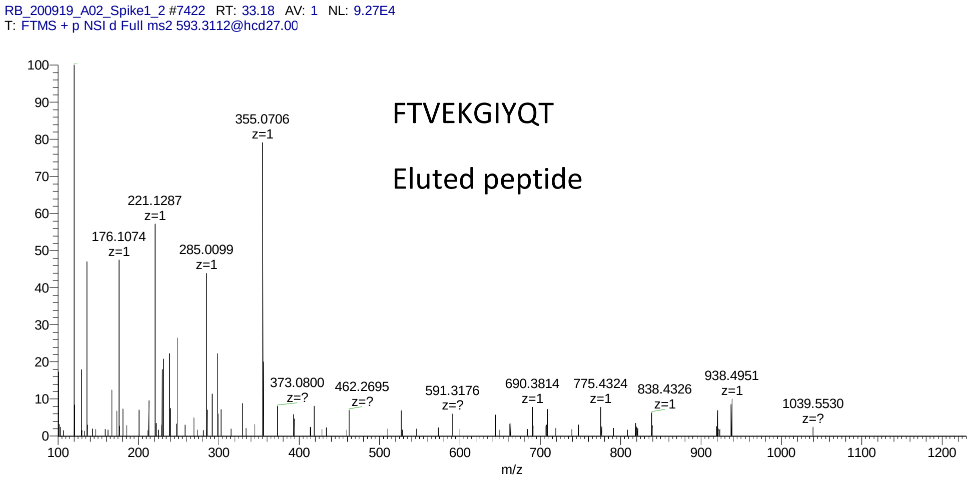
RB_200919_A02_Spike1_2 #7422 RT: 33.18 AV: 1 NL: 9.27E4
T: FTMS + p NSI d Full ms2 593.3112@hcd27.00
FTVEKGIYQT
Eluted peptide
355.0706 z=1
221.1287 z=1
176.1074 z=1
285.0099 z=1
373.0800 z=?
462.2695 z=?
591.3176 z=?
690.3814 z=1
775.4324 z=1
838.4326 z=1
938.4951 z=1
1039.5530 z=?
m/z

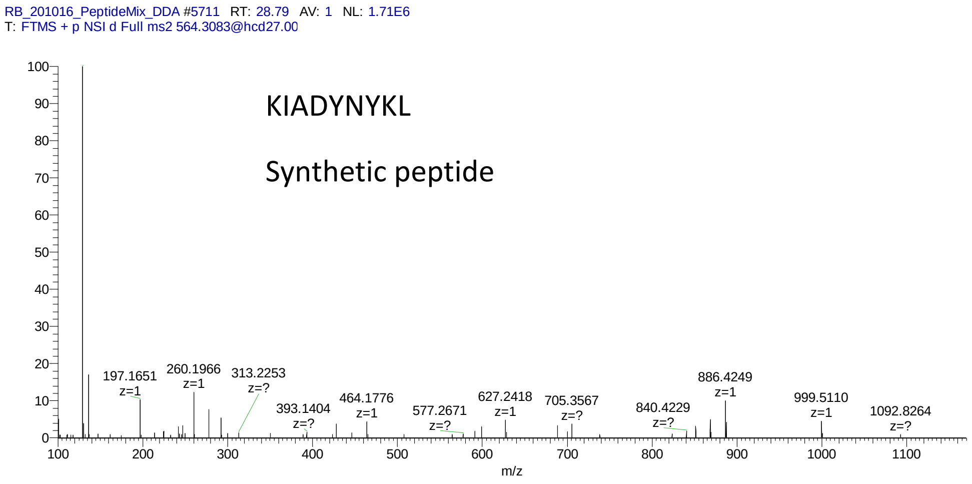
RB_201016_PeptideMix_DDA #5711 RT: 28.79 AV: 1 NL: 1.71E6
T: FTMS + p NSI d Full ms2 564.3083@hcd27.00
KIADYNYKL
Synthetic peptide
197.1651 z=1
260.1966 z=1
313.2253 z=?
393.1404 z=?
464.1776 z=1
577.2671 z=?
627.2418 z=1
705.3567 z=?
840.4229 z=?
886.4249 z=1
999.5110 z=1
1092.8264 z=?
m/z

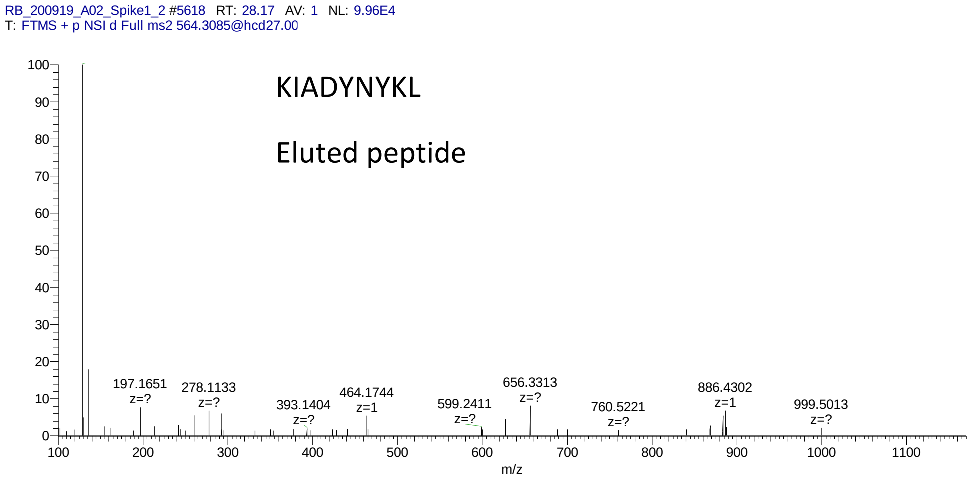
RB_200919_A02_Spike1_2 #5618 RT: 28.17 AV: 1 NL: 9.96E4
T: FTMS + p NSI d Full ms2 564.3085@hcd27.00
KIADYNYKL
Eluted peptide
197.1651 z=?
278.1133 z=?
393.1404 z=?
464.1744 z=1
599.2411 z=?
656.3313 z=?
760.5221 z=?
886.4302 z=1
999.5013 z=?
m/z

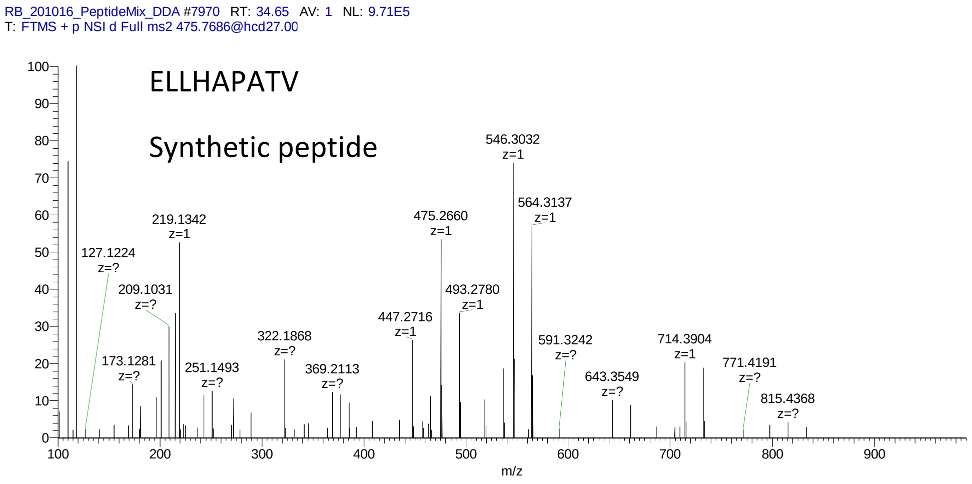

RB_201016_PeptideMix_DDA #7970 RT: 34.65 AV: 1 NL: 9.71E5
T: FTMS + p NSI d Full ms2 475.7686@hcd27.00
ELLHAPATV
Synthetic peptide
546.3032 z=1
564.3137 z=1
475.2660 z=1
219.1342 z=1
127.1224 z=?
209.1031 z=?
493.2780 z=1
447.2716 z=1
322.1868 z=?
591.3242 z=?
714.3904 z=1
173.1281 z=?
251.1493 z=?
369.2113 z=?
771.4191 z=?
643.3549 z=?
815.4368 z=?
m/z


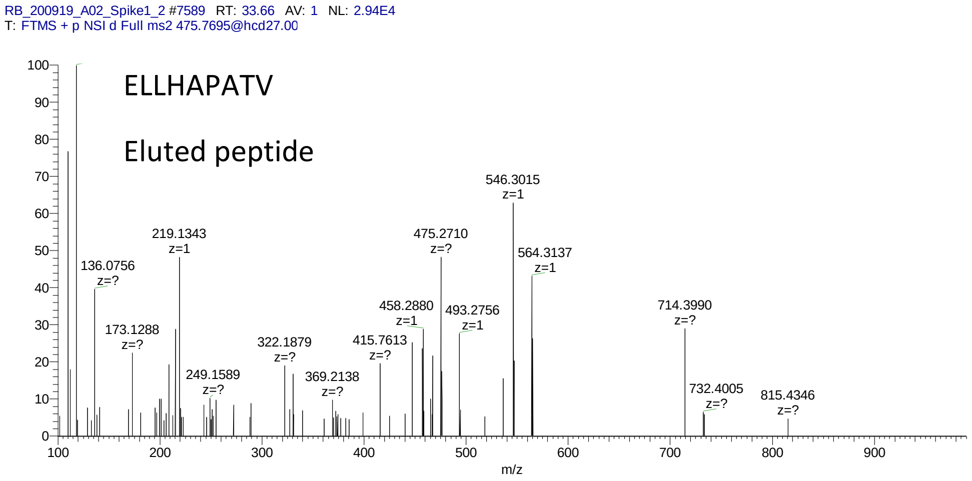

RB_200919_A02_Spike1_2 #7589 RT: 33.66 AV: 1 NL: 2.94E4
T: FTMS + p NSI d Full ms2 475.7695@hcd27.00
ELLHAPATV
Eluted peptide
546.3015 z=1
219.1343 z=1
475.2710 z=?
564.3137 z=1
136.0756 z=?
458.2880 z=1
493.2756 z=1
714.3990 z=?
173.1288 z=?
322.1879 z=?
415.7613 z=?
249.1589 z=?
369.2138 z=?
732.4005 z=?
815.4346 z=?
m/z

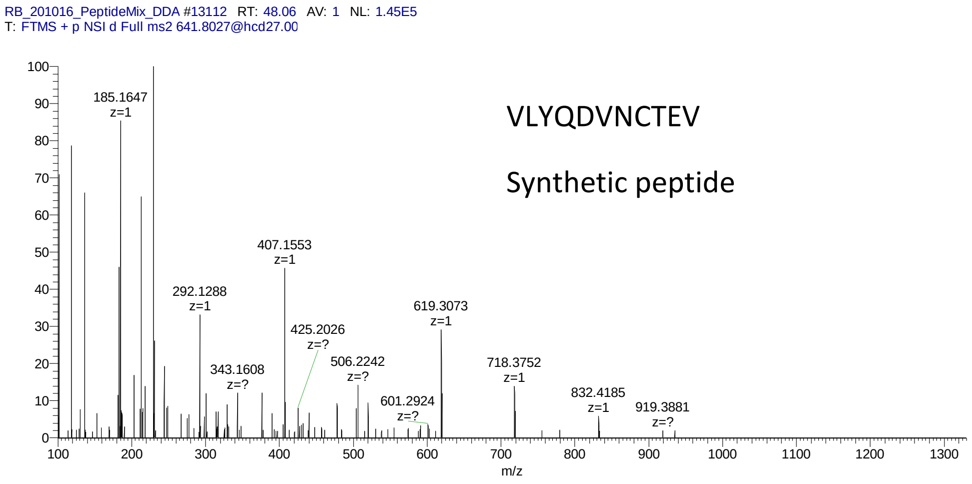
RB_201016_PeptideMix_DDA #13112 RT: 48.06 AV: 1 NL: 1.45E5
T: FTMS + p NSI d Full ms2 641.8027@hcd27.00
VLYQDVNCTEV
Synthetic peptide
185.1647 z=1
292.1288 z=1
343.1608 z=?
407.1553 z=1
425.2026 z=?
506.2242 z=?
601.2924 z=?
619.3073 z=1
718.3752 z=1
832.4185 z=1
919.3881 z=?
m/z

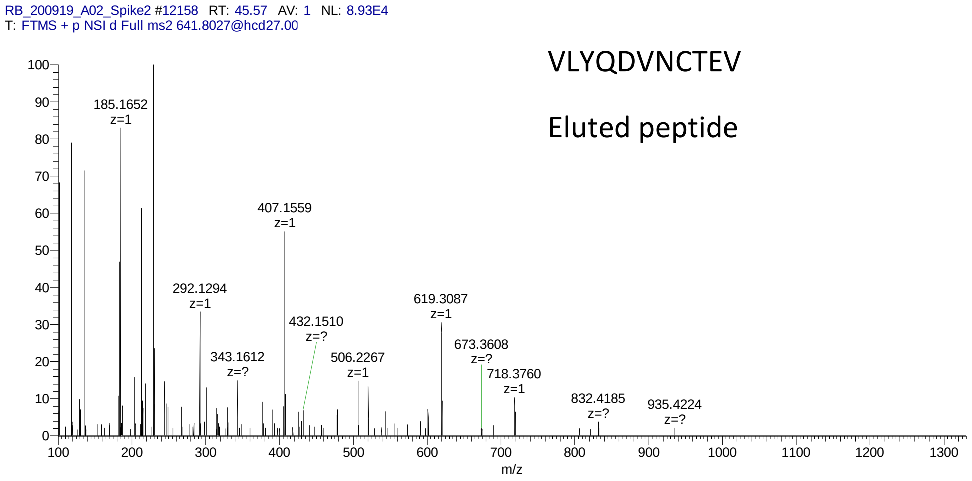
RB_200919_A02_Spike2 #12158 RT: 45.57 AV: 1 NL: 8.93E4
T: FTMS + p NSI d Full ms2 641.8027@hcd27.00
VLYQDVNCTEV
Eluted peptide
185.1652 z=1
292.1294 z=1
343.1612 z=?
407.1559 z=1
432.1510 z=?
506.2267 z=1
619.3087 z=1
673.3608 z=?
718.3760 z=1
832.4185 z=?
935.4224 z=?
m/z

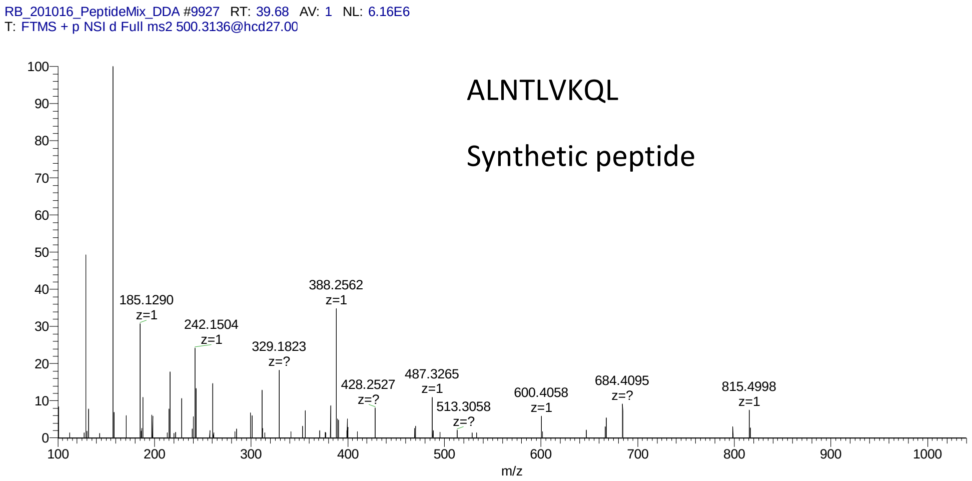

RB_201016_PeptideMix_DDA #9927 RT: 39.68 AV: 1 NL: 6.16E6
T: FTMS + p NSI d Full ms2 500.3136@hcd27.00
ALNTLVKQL
Synthetic peptide
185.1290
z=1
242.1504
z=1
329.1823
z=?
388.2562
z=1
428.2527
z=?
487.3265
z=1
513.3058
z=?
600.4058
z=1
684.4095
z=?
815.4998
z=1
m/z


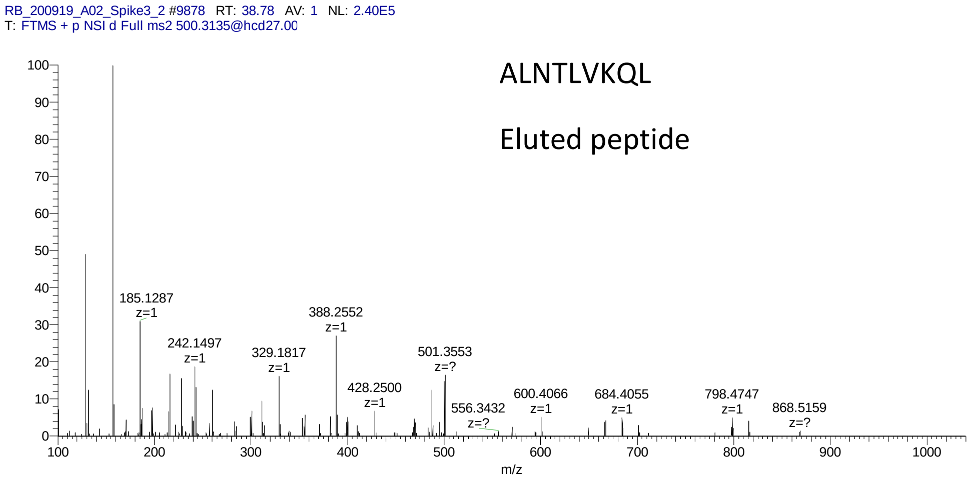

RB_200919_A02_Spike3_2 #9878 RT: 38.78 AV: 1 NL: 2.40E5
T: FTMS + p NSI d Full ms2 500.3135@hcd27.00
ALNTLVKQL
Eluted peptide
185.1287
z=1
242.1497
z=1
329.1817
z=1
388.2552
z=1
428.2500
z=1
501.3553
z=?
556.3432
z=?
600.4066
z=1
684.4055
z=1
798.4747
z=1
868.5159
z=?
m/z

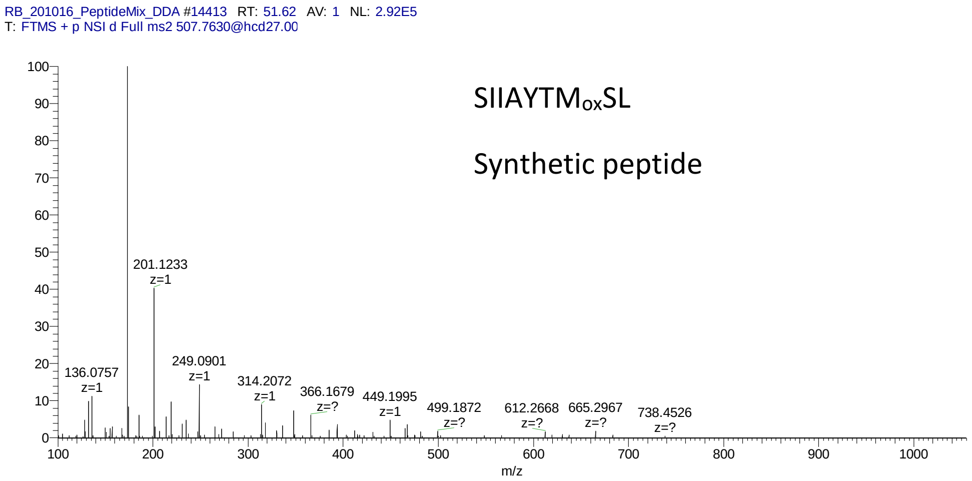
RB_201016_PeptideMix_DDA #14413 RT: 51.62 AV: 1 NL: 2.92E5
T: FTMS + p NSI d Full ms2 507.7630@hcd27.00
SIIAYTMoxSL
Synthetic peptide
201.1233
z=1
136.0757
z=1
249.0901
z=1
314.2072
z=1
366.1679
z=?
449.1995
z=1
499.1872
z=?
612.2668
z=?
665.2967
z=?
738.4526
z=?
m/z

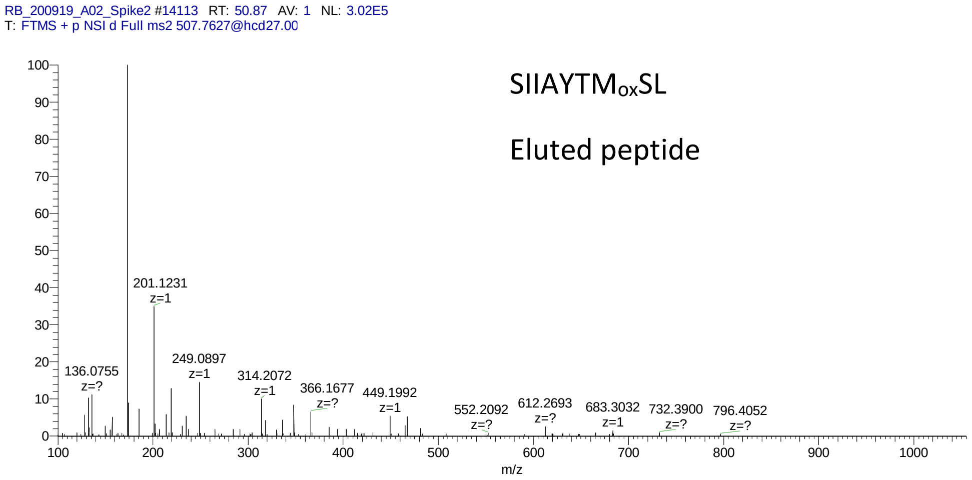
RB_200919_A02_Spike2 #14113 RT: 50.87 AV: 1 NL: 3.02E5
T: FTMS + p NSI d Full ms2 507.7627@hcd27.00
SIIAYTMoxSL
Eluted peptide
201.1231
z=1
136.0755
z=?
249.0897
z=1
314.2072
z=1
366.1677
z=?
449.1992
z=1
552.2092
z=?
612.2693
z=?
683.3032
z=1
732.3900
z=?
796.4052
z=?
m/z

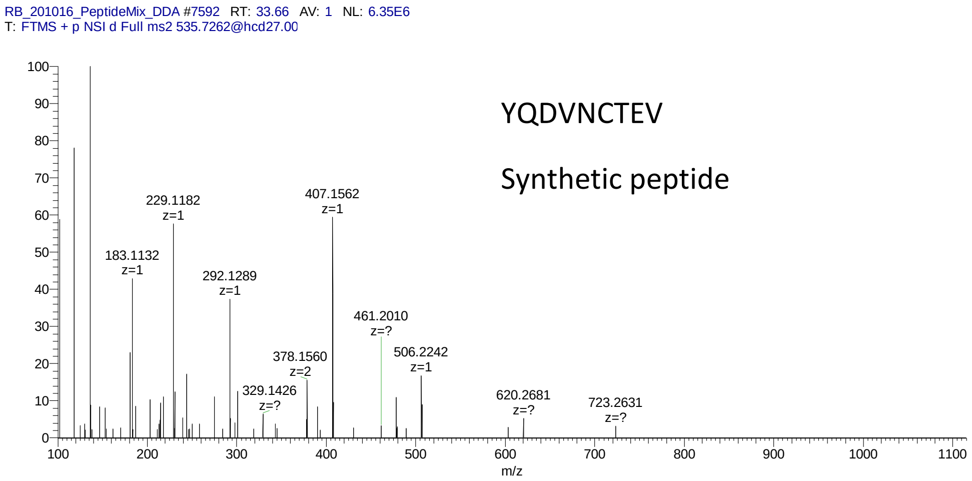
RB_201016_PeptideMix_DDA #7592 RT: 33.66 AV: 1 NL: 6.35E6
T: FTMS + p NSI d Full ms2 535.7262@hcd27.00
YQDVNCTEV
Synthetic peptide
229.1182 z=1
183.1132 z=1
292.1289 z=1
407.1562 z=1
461.2010 z=?
378.1560 z=2
506.2242 z=1
329.1426 z=?
620.2681 z=?
723.2631 z=?
m/z

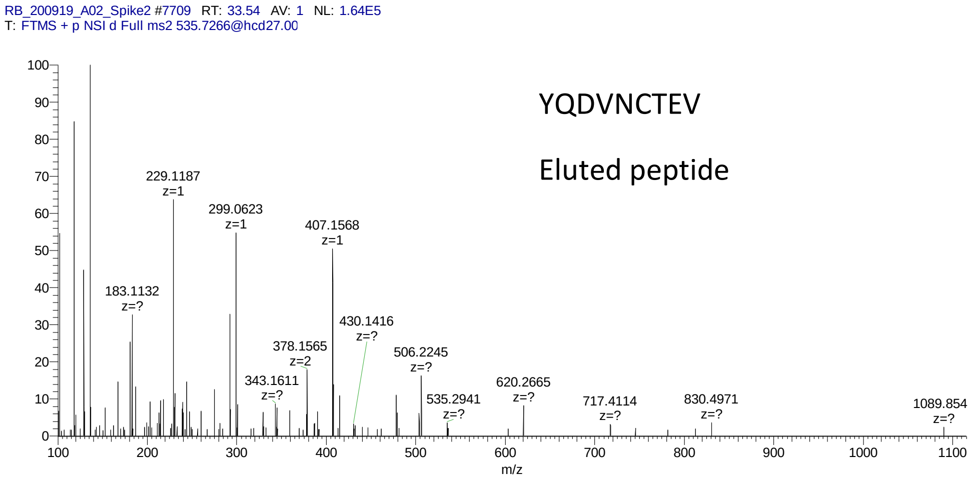
RB_200919_A02_Spike2 #7709 RT: 33.54 AV: 1 NL: 1.64E5
T: FTMS + p NSI d Full ms2 535.7266@hcd27.00
YQDVNCTEV
Eluted peptide
229.1187 z=1
299.0623 z=1
407.1568 z=1
183.1132 z=?
430.1416 z=?
378.1565 z=2
506.2245 z=?
343.1611 z=?
535.2941 z=?
620.2665 z=?
717.4114 z=?
830.4971 z=?
1089.854 z=?
m/z

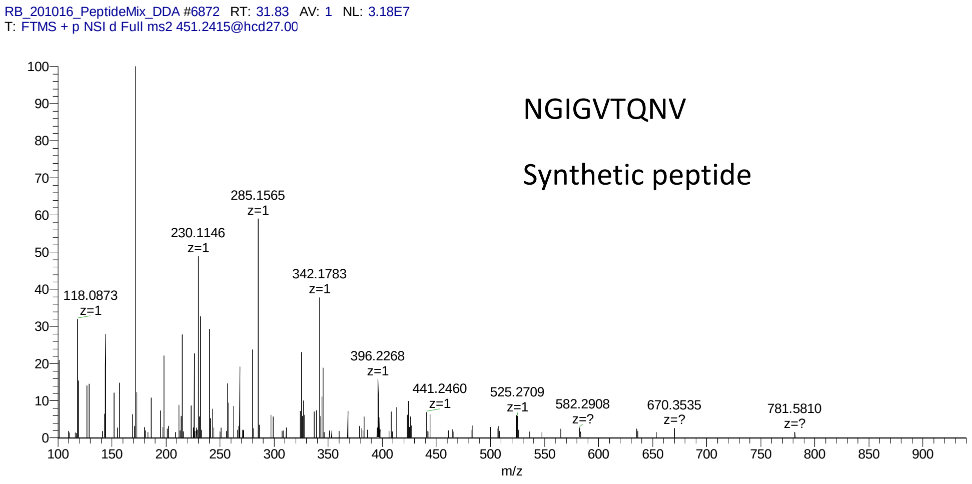
RB_201016_PeptideMix_DDA #6872 RT: 31.83 AV: 1 NL: 3.18E7
T: FTMS + p NSI d Full ms2 451.2415@hcd27.00
NGIGVTQNV
Synthetic peptide
118.0873 z=1
230.1146 z=1
285.1565 z=1
342.1783 z=1
396.2268 z=1
441.2460 z=1
525.2709 z=1
582.2908 z=?
670.3535 z=?
781.5810 z=?
m/z

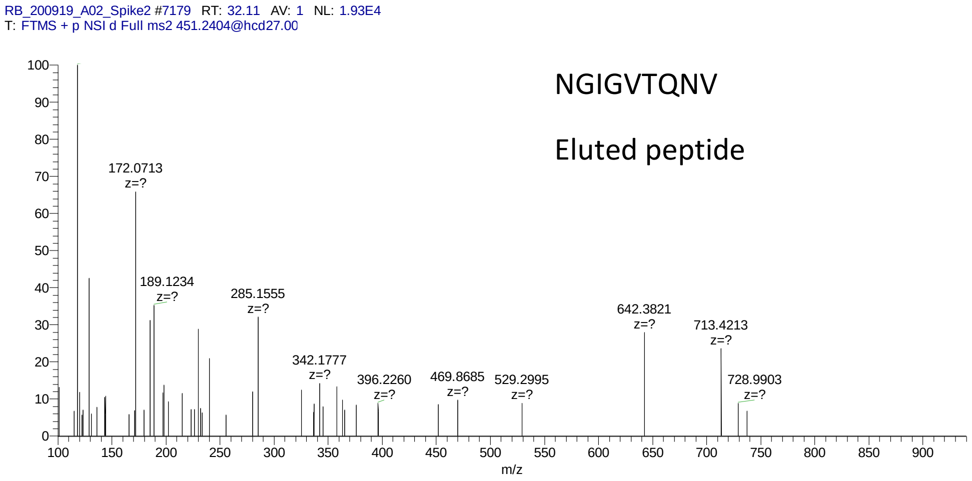
RB_200919_A02_Spike2 #7179 RT: 32.11 AV: 1 NL: 1.93E4
T: FTMS + p NSI d Full ms2 451.2404@hcd27.00
NGIGVTQNV
Eluted peptide
172.0713 z=?
189.1234 z=?
285.1555 z=?
342.1777 z=?
396.2260 z=?
469.8685 z=?
529.2995 z=?
642.3821 z=?
713.4213 z=?
728.9903 z=?
m/z

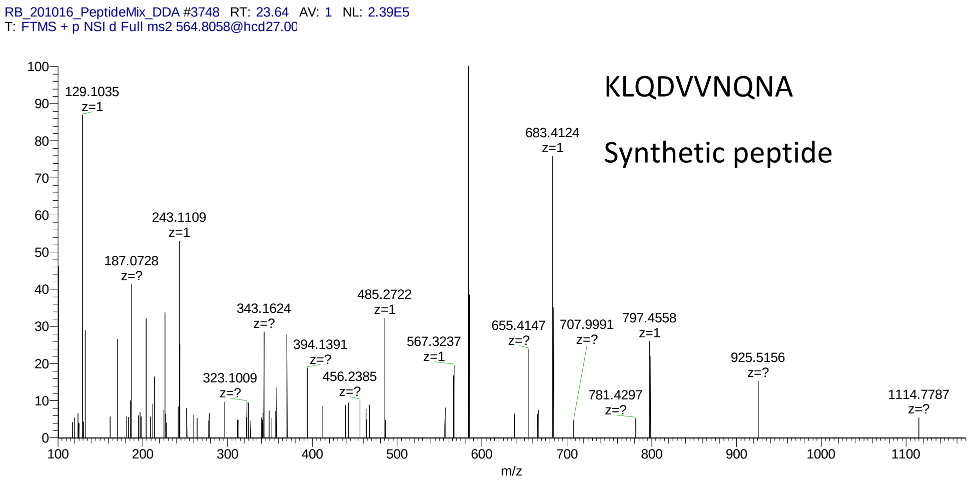
RB_201016_PeptideMix_DDA #3748 RT: 23.64 AV: 1 NL: 2.39E5
T: FTMS + p NSI d Full ms2 564.8058@hcd27.00
KLQDVVNQNA
Synthetic peptide
129.1035
z=1
683.4124
z=1
243.1109
z=1
187.0728
z=?
485.2722
z=1
343.1624
z=?
394.1391
z=?
567.3237
z=1
655.4147
z=?
707.9991
z=?
797.4558
z=1
925.5156
z=?
323.1009
z=?
456.2385
z=?
781.4297
z=?
1114.7787
z=?
m/z

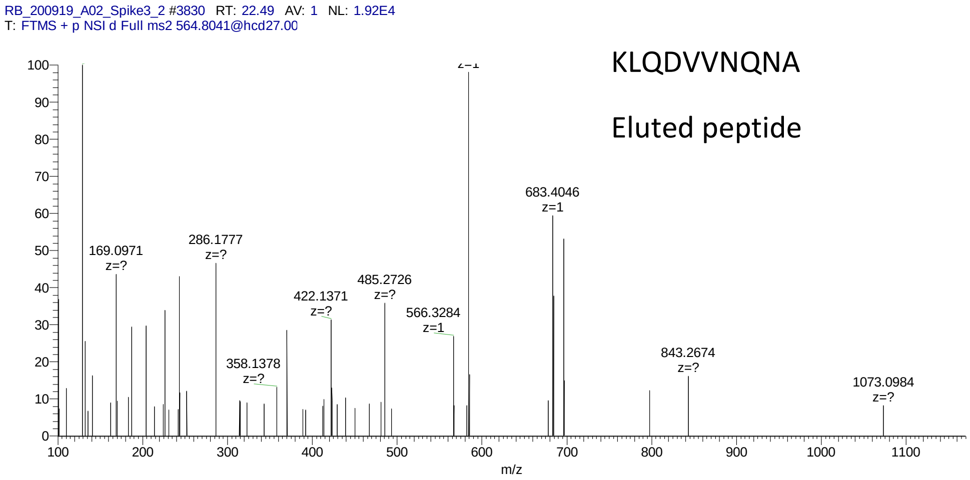
RB_200919_A02_Spike3_2 #3830 RT: 22.49 AV: 1 NL: 1.92E4
T: FTMS + p NSI d Full ms2 564.8041@hcd27.00
KLQDVVNQNA
Eluted peptide
683.4046
z=1
169.0971
z=?
286.1777
z=?
485.2726
z=?
422.1371
z=?
566.3284
z=1
358.1378
z=?
843.2674
z=?
1073.0984
z=?
m/z

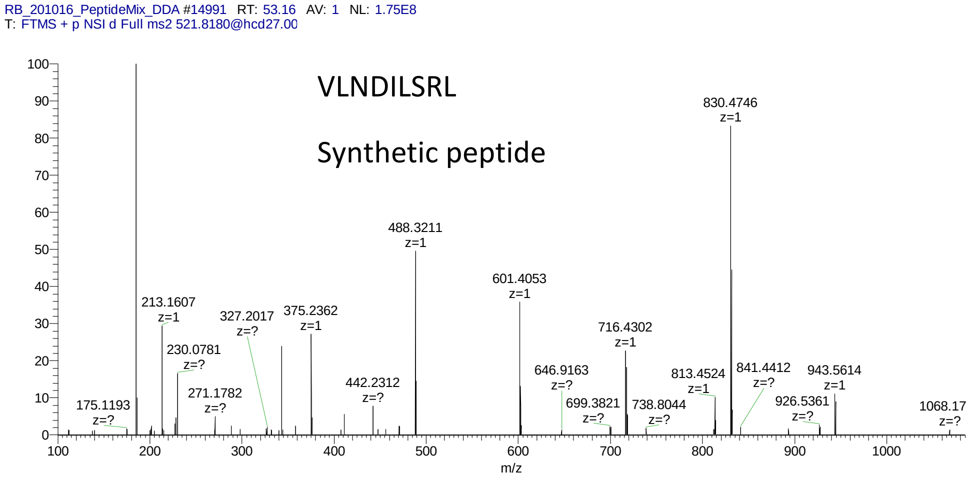
RB_201016_PeptideMix_DDA #14991 RT: 53.16 AV: 1 NL: 1.75E8
T: FTMS + p NSI d Full ms2 521.8180@hcd27.00
VLNDILSRL
Synthetic peptide
830.4746 z=1
488.3211 z=1
601.4053 z=1
213.1607 z=1
327.2017 z=?
375.2362 z=1
716.4302 z=1
230.0781 z=?
646.9163 z=?
813.4524 z=1
841.4412 z=?
943.5614 z=1
442.2312 z=?
271.1782 z=?
175.1193 z=?
699.3821 z=?
738.8044 z=?
926.5361 z=?
1068.17 z=?
m/z

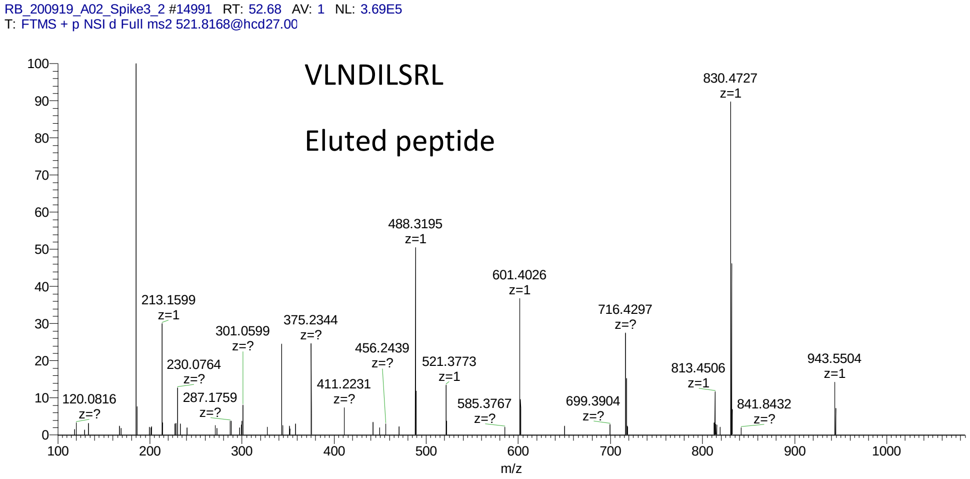
RB_200919_A02_Spike3_2 #14991 RT: 52.68 AV: 1 NL: 3.69E5
T: FTMS + p NSI d Full ms2 521.8168@hcd27.00
VLNDILSRL
Eluted peptide
830.4727 z=1
488.3195 z=1
601.4026 z=1
213.1599 z=1
716.4297 z=?
301.0599 z=?
375.2344 z=?
456.2439 z=?
521.3773 z=1
230.0764 z=?
813.4506 z=1
943.5504 z=1
411.2231 z=?
120.0816 z=?
287.1759 z=?
585.3767 z=?
699.3904 z=?
841.8432 z=?
m/z

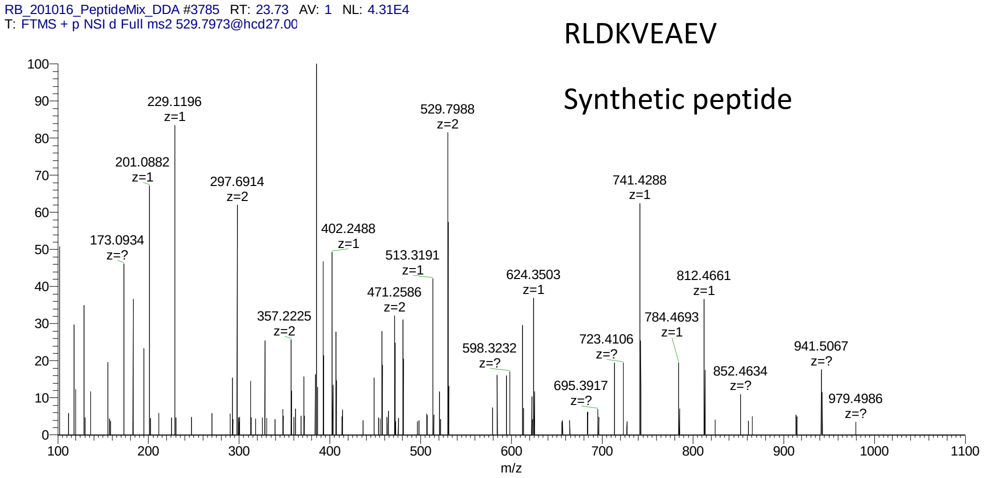
RB_201016_PeptideMix_DDA #3785 RT: 23.73 AV: 1 NL: 4.31E4
T: FTMS + p NSI d Full ms2 529.7973@hcd27.00
RLDKVEAEV
Synthetic peptide
229.1196 z=1
201.0882 z=1
173.0934 z=?
297.6914 z=2
357.2225 z=2
402.2488 z=1
471.2586 z=2
513.3191 z=1
529.7988 z=2
598.3232 z=?
624.3503 z=1
695.3917 z=?
723.4106 z=?
741.4288 z=1
784.4693 z=1
812.4661 z=1
852.4634 z=?
941.5067 z=?
979.4986 z=?
m/z

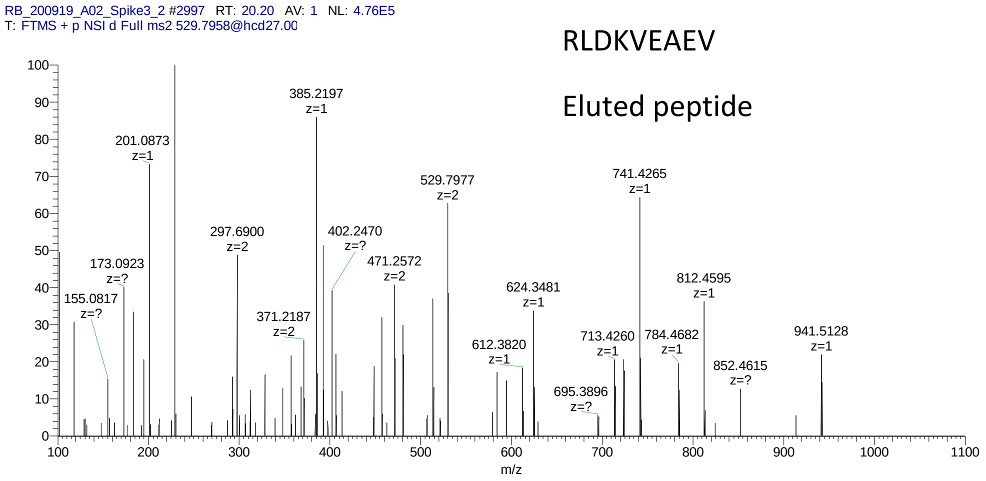
RB_200919_A02_Spike3_2 #2997 RT: 20.20 AV: 1 NL: 4.76E5
T: FTMS + p NSI d Full ms2 529.7958@hcd27.00
RLDKVEAEV
Eluted peptide
385.2197 z=1
201.0873 z=1
173.0923 z=?
155.0817 z=?
297.6900 z=2
371.2187 z=2
402.2470 z=?
471.2572 z=2
529.7977 z=2
612.3820 z=1
624.3481 z=1
695.3896 z=?
713.4260 z=1
741.4265 z=1
784.4682 z=1
812.4595 z=1
852.4615 z=?
941.5128 z=1
m/z

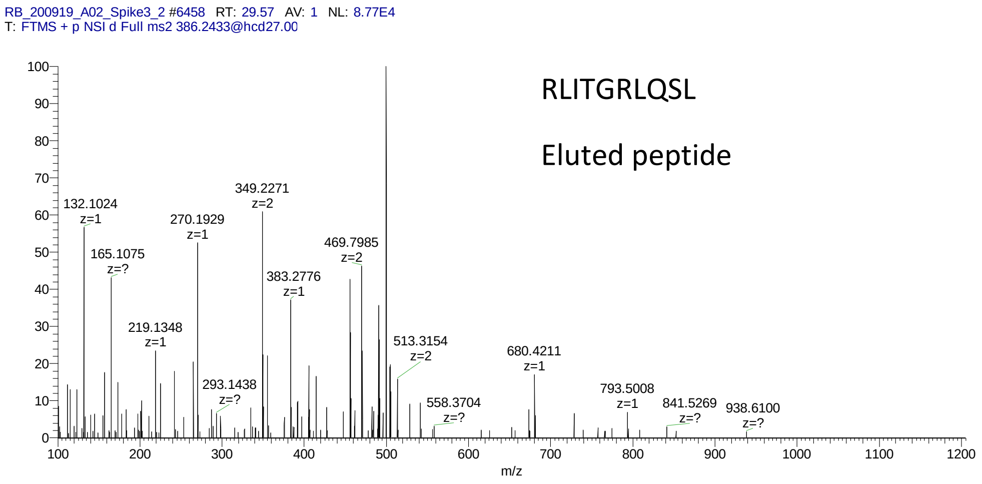
RB_200919_A02_Spike3_2 #6458 RT: 29.57 AV: 1 NL: 8.77E4
T: FTMS + p NSI d Full ms2 386.2433@hcd27.00
RLITGRLQSL
Eluted peptide
132.1024 z=1
165.1075 z=?
219.1348 z=1
270.1929 z=1
293.1438 z=?
349.2271 z=2
383.2776 z=1
469.7985 z=2
513.3154 z=2
558.3704 z=?
680.4211 z=1
793.5008 z=1
841.5269 z=?
938.6100 z=?
m/z

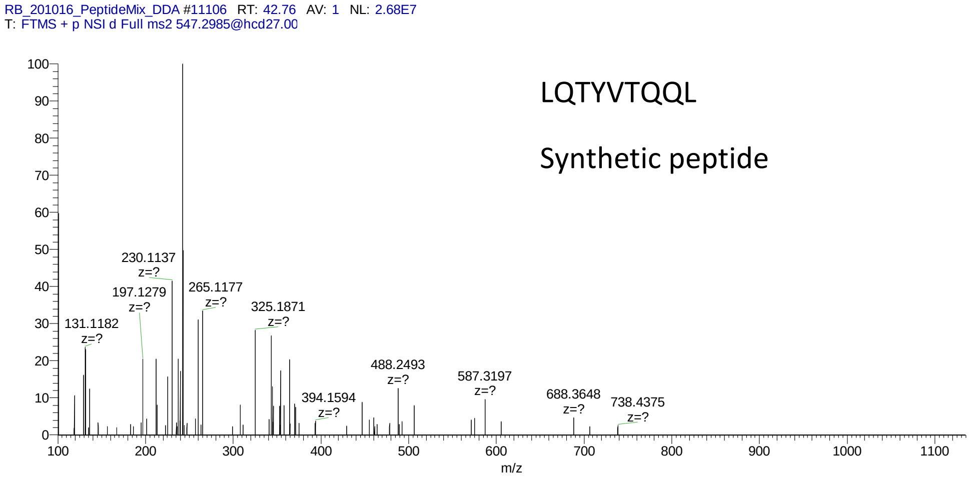
RB_201016_PeptideMix_DDA #11106 RT: 42.76 AV: 1 NL: 2.68E7
T: FTMS + p NSI d Full ms2 547.2985@hcd27.00
LQTYVTQQL
Synthetic peptide
230.1137 z=?
197.1279 z=?
265.1177 z=?
325.1871 z=?
131.1182 z=?
488.2493 z=?
587.3197 z=?
394.1594 z=?
688.3648 z=?
738.4375 z=?
m/z

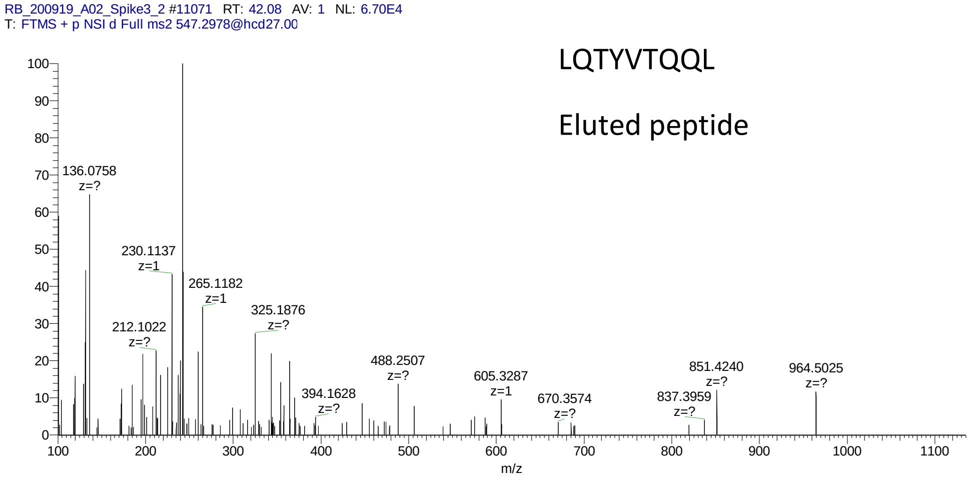
RB_200919_A02_Spike3_2 #11071 RT: 42.08 AV: 1 NL: 6.70E4
T: FTMS + p NSI d Full ms2 547.2978@hcd27.00
LQTYVTQQL
Eluted peptide
136.0758 z=?
230.1137 z=1
265.1182 z=1
325.1876 z=?
212.1022 z=?
488.2507 z=?
605.3287 z=1
394.1628 z=?
670.3574 z=?
851.4240 z=?
837.3959 z=?
964.5025 z=?
m/z

RB_201016_PeptideMix_DDA #2218 RT: 19.48 AV: 1 NL: 5.14E6
T: FTMS + p NSI d Full ms2 528.8238@hcd27.00

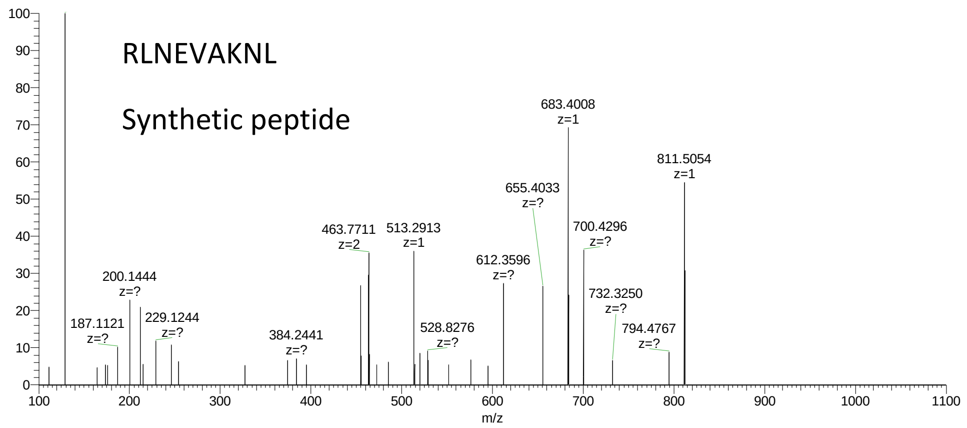


RB_200919_A02_Spike3_2 #2499 RT: 18.80 AV: 1 NL: 2.68E4
T: FTMS + p NSI d Full ms2 528.8110@hcd27.00

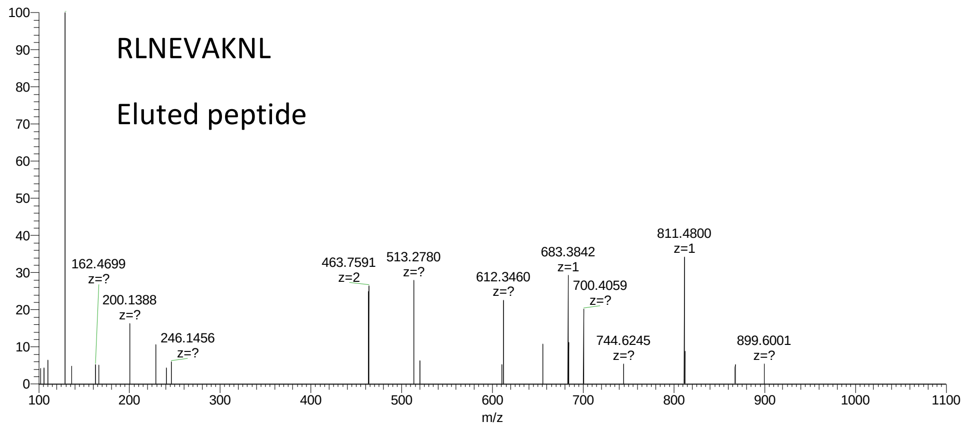

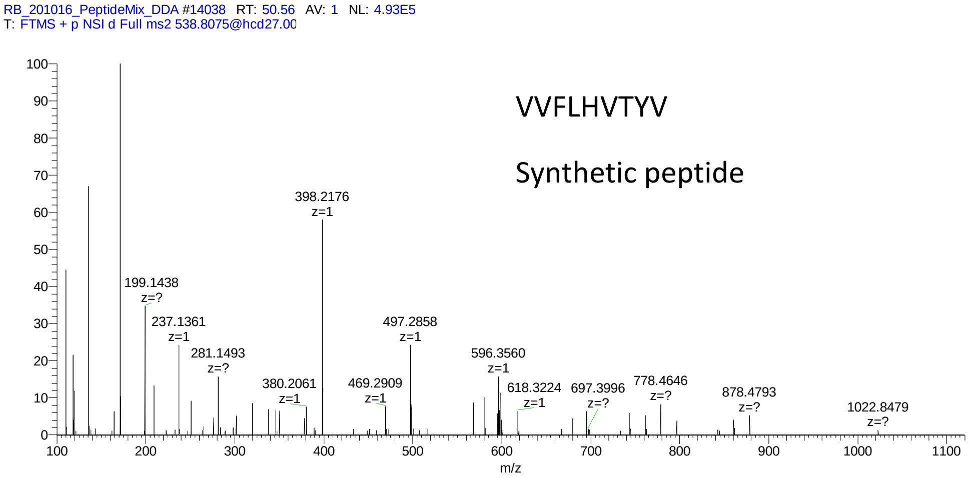
RB_201016_PeptideMix_DDA #14038 RT: 50.56 AV: 1 NL: 4.93E5
T: FTMS + p NSI d Full ms2 538.8075@hcd27.00
VVFLHVTYV
Synthetic peptide
398.2176 z=1
199.1438 z=?
237.1361 z=1
281.1493 z=?
380.2061 z=1
469.2909 z=1
497.2858 z=1
596.3560 z=1
618.3224 z=1
697.3996 z=?
778.4646 z=?
878.4793 z=?
1022.8479 z=?
m/z

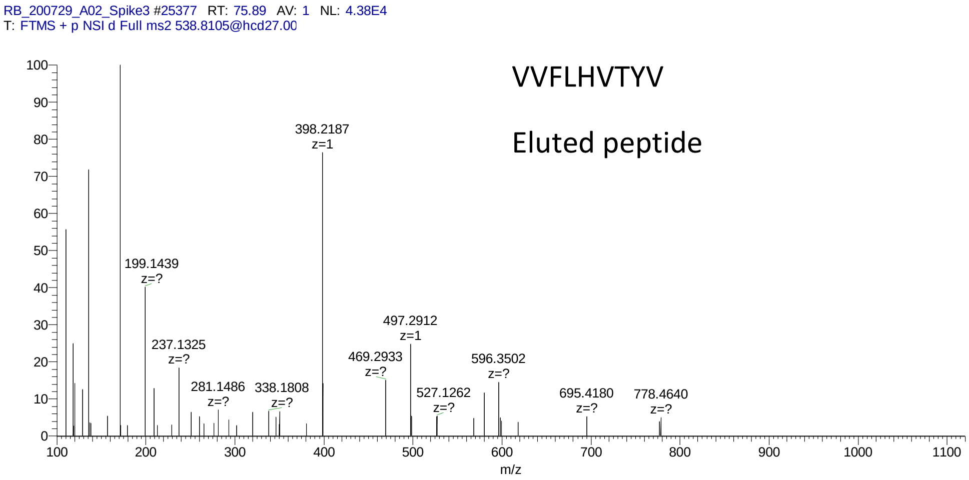
RB_200729_A02_Spike3 #25377 RT: 75.89 AV: 1 NL: 4.38E4
T: FTMS + p NSI d Full ms2 538.8105@hcd27.00
VVFLHVTYV
Eluted peptide
398.2187 z=1
199.1439 z=?
237.1325 z=?
281.1486 z=?
338.1808 z=?
469.2933 z=?
497.2912 z=1
527.1262 z=?
596.3502 z=?
695.4180 z=?
778.4640 z=?
m/z

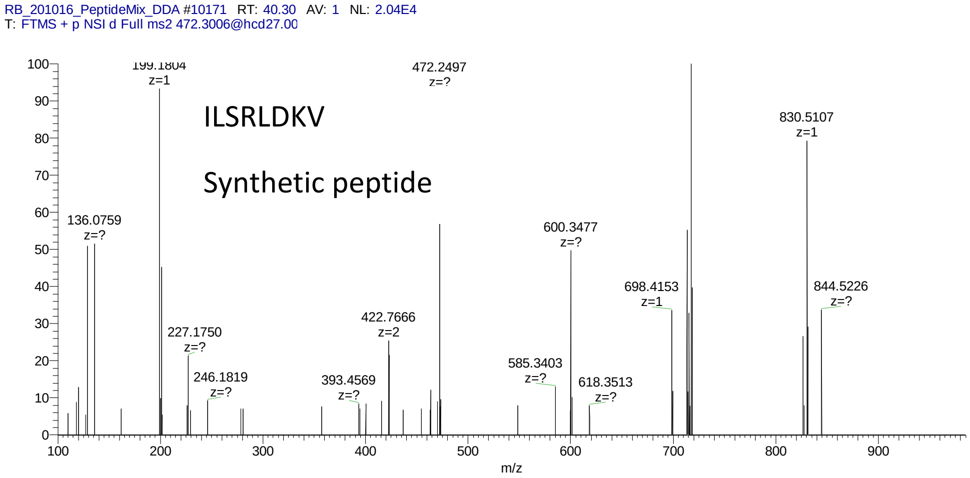
RB_201016_PeptideMix_DDA #10171 RT: 40.30 AV: 1 NL: 2.04E4
T: FTMS + p NSI d Full ms2 472.3006@hcd27.00
ILSRLDKV
Synthetic peptide
199.1804 z=1
472.2497 z=?
830.5107 z=1
136.0759 z=?
600.3477 z=?
698.4153 z=1
844.5226 z=?
422.7666 z=2
227.1750 z=?
246.1819 z=?
393.4569 z=?
585.3403 z=?
618.3513 z=?
m/z

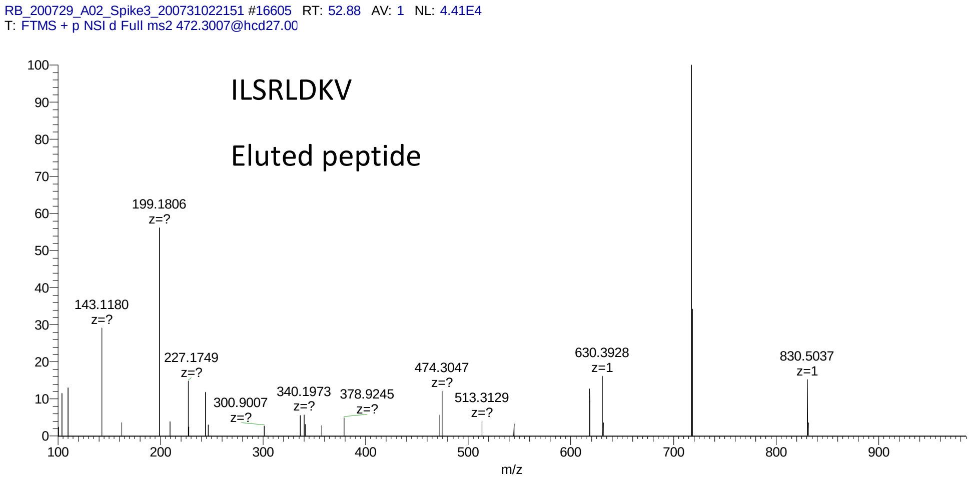
RB_200729_A02_Spike3_200731022151 #16605 RT: 52.88 AV: 1 NL: 4.41E4
T: FTMS + p NSI d Full ms2 472.3007@hcd27.00
ILSRLDKV
Eluted peptide
199.1806 z=?
143.1180 z=?
227.1749 z=?
300.9007 z=?
340.1973 z=?
378.9245 z=?
474.3047 z=?
513.3129 z=?
630.3928 z=1
830.5037 z=1
m/z

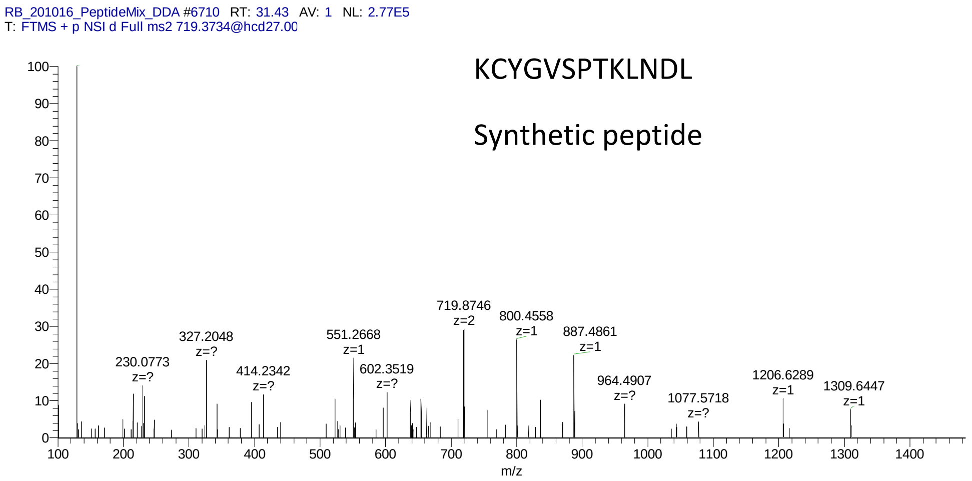
RB_201016_PeptideMix_DDA #6710 RT: 31.43 AV: 1 NL: 2.77E5
T: FTMS + p NSI d Full ms2 719.3734@hcd27.00
KCYGVSPTKLNDL
Synthetic peptide
230.0773 z=?
327.2048 z=?
414.2342 z=?
551.2668 z=1
602.3519 z=?
719.8746 z=2
800.4558 z=1
887.4861 z=1
964.4907 z=?
1077.5718 z=?
1206.6289 z=1
1309.6447 z=1
m/z

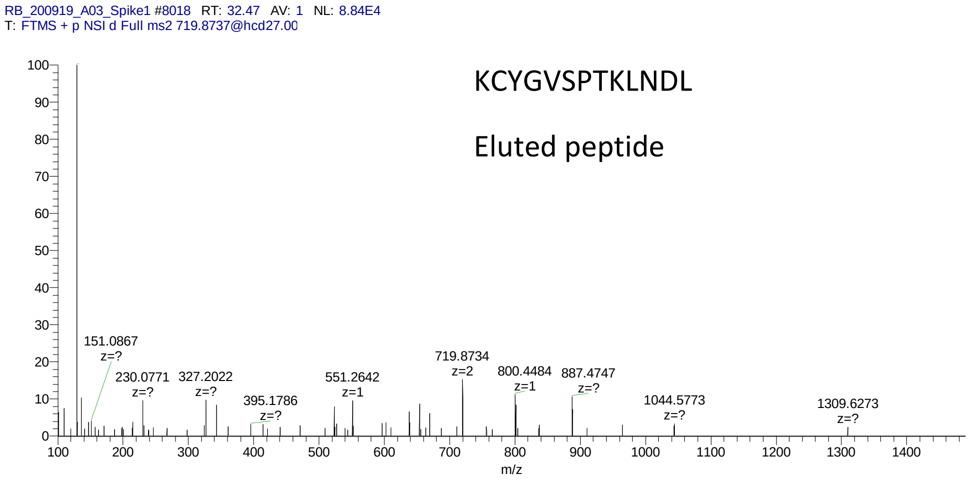
RB_200919_A03_Spike1 #8018 RT: 32.47 AV: 1 NL: 8.84E4
T: FTMS + p NSI d Full ms2 719.8737@hcd27.00
KCYGVSPTKLNDL
Eluted peptide
151.0867 z=?
230.0771 z=?
327.2022 z=?
395.1786 z=?
551.2642 z=1
719.8734 z=2
800.4484 z=1
887.4747 z=?
1044.5773 z=?
1309.6273 z=?
m/z

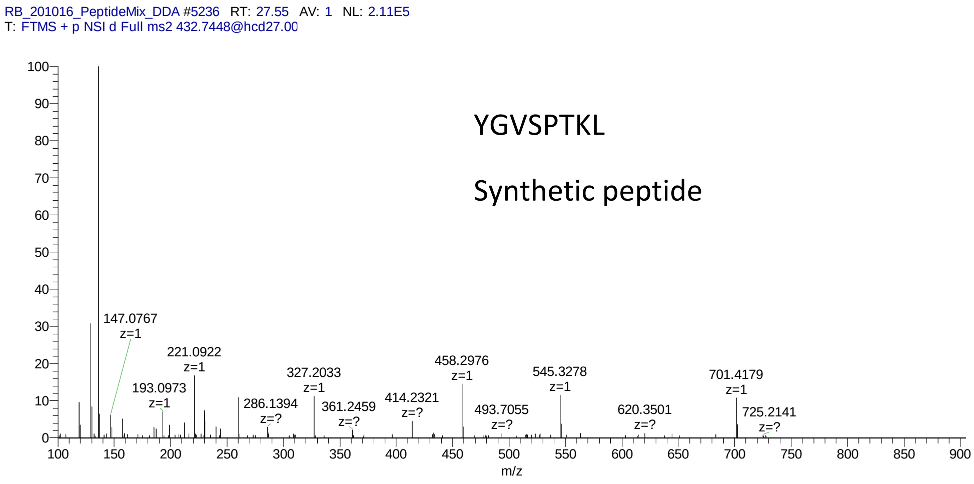
RB_201016_PeptideMix_DDA #5236 RT: 27.55 AV: 1 NL: 2.11E5
T: FTMS + p NSI d Full ms2 432.7448@hcd27.00
YGVSPTKL
Synthetic peptide
147.0767 z=1
221.0922 z=1
193.0973 z=1
286.1394 z=?
327.2033 z=1
361.2459 z=?
414.2321 z=?
458.2976 z=1
493.7055 z=?
545.3278 z=1
620.3501 z=?
701.4179 z=1
725.2141 z=?
m/z

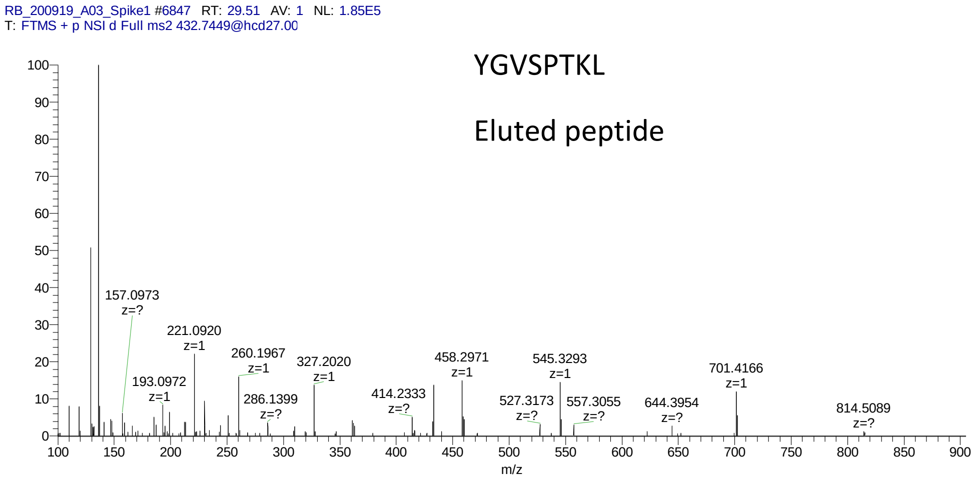
RB_200919_A03_Spike1 #6847 RT: 29.51 AV: 1 NL: 1.85E5
T: FTMS + p NSI d Full ms2 432.7449@hcd27.00
YGVSPTKL
Eluted peptide
157.0973 z=?
221.0920 z=1
193.0972 z=1
260.1967 z=1
286.1399 z=?
327.2020 z=1
414.2333 z=?
458.2971 z=1
527.3173 z=?
545.3293 z=1
557.3055 z=?
644.3954 z=?
701.4166 z=1
814.5089 z=?
m/z

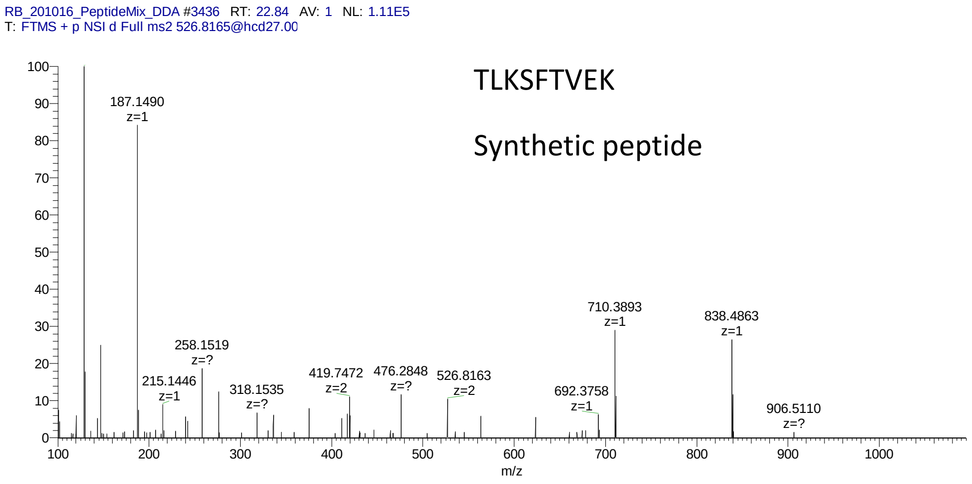
RB_201016_PeptideMix_DDA #3436 RT: 22.84 AV: 1 NL: 1.11E5
T: FTMS + p NSI d Full ms2 526.8165@hcd27.00
TLKSFTVEK
Synthetic peptide
187.1490
z=1
215.1446
z=1
258.1519
z=?
318.1535
z=?
419.7472
z=2
476.2848
z=?
526.8163
z=2
692.3758
z=1
710.3893
z=1
838.4863
z=1
906.5110
z=?
m/z

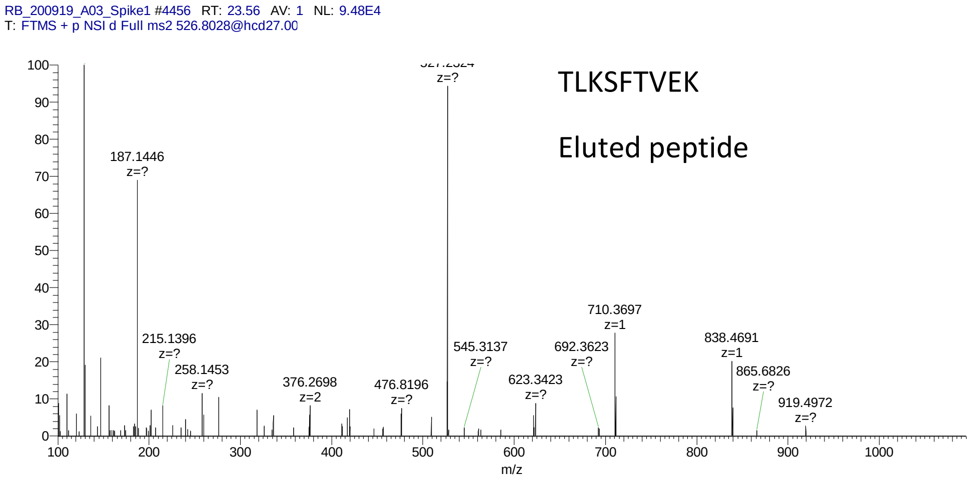
RB_200919_A03_Spike1 #4456 RT: 23.56 AV: 1 NL: 9.48E4
T: FTMS + p NSI d Full ms2 526.8028@hcd27.00
TLKSFTVEK
Eluted peptide
z=?
187.1446
z=?
215.1396
z=?
258.1453
z=?
376.2698
z=2
476.8196
z=?
545.3137
z=?
623.3423
z=?
692.3623
z=?
710.3697
z=1
838.4691
z=1
865.6826
z=?
919.4972
z=?
m/z

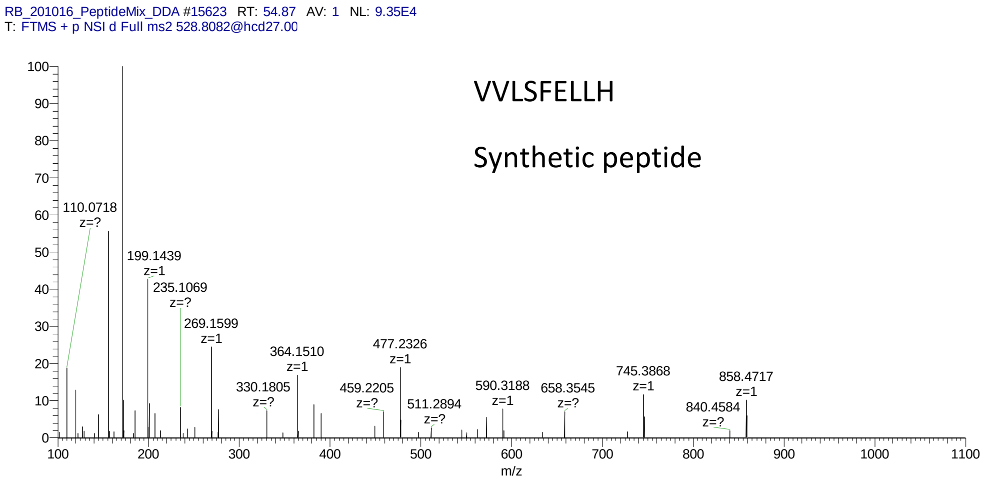
RB_201016_PeptideMix_DDA #15623 RT: 54.87 AV: 1 NL: 9.35E4
T: FTMS + p NSI d Full ms2 528.8082@hcd27.00
VVLSFELLH
Synthetic peptide
110.0718
z=?
199.1439
z=1
235.1069
z=?
269.1599
z=1
330.1805
z=?
364.1510
z=1
459.2205
z=?
477.2326
z=1
511.2894
z=?
590.3188
z=1
658.3545
z=?
745.3868
z=1
840.4584
z=?
858.4717
z=1
m/z

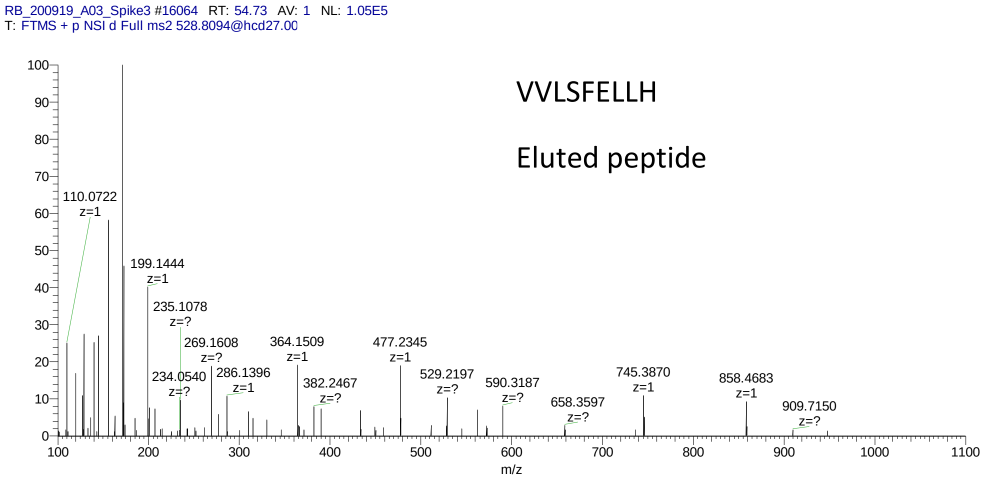
RB_200919_A03_Spike3 #16064 RT: 54.73 AV: 1 NL: 1.05E5
T: FTMS + p NSI d Full ms2 528.8094@hcd27.00
VVLSFELLH
Eluted peptide
110.0722
z=1
199.1444
z=1
235.1078
z=?
234.0540
z=?
269.1608
z=?
286.1396
z=1
364.1509
z=1
382.2467
z=?
477.2345
z=1
529.2197
z=?
590.3187
z=?
658.3597
z=?
745.3870
z=1
858.4683
z=1
909.7150
z=?
m/z

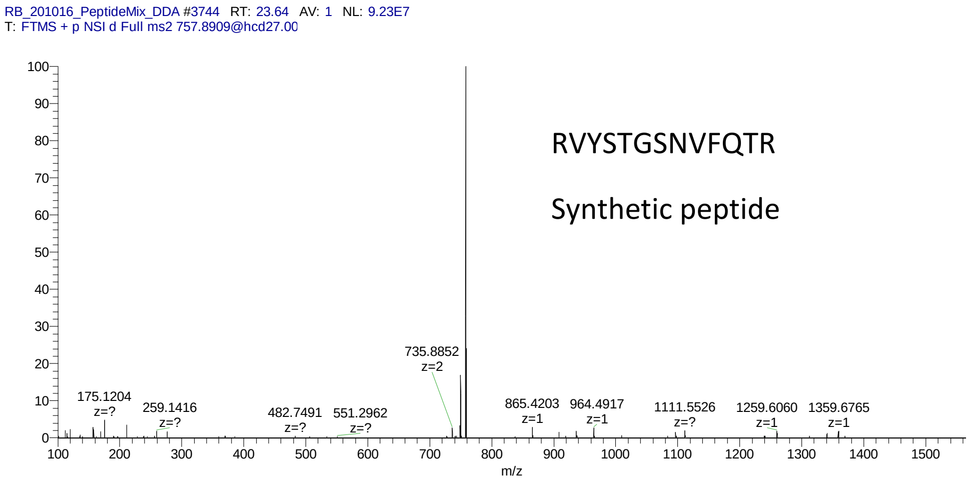

RB_201016_PeptideMix_DDA #3744 RT: 23.64 AV: 1 NL: 9.23E7
T: FTMS + p NSI d Full ms2 757.8909@hcd27.00
RVYSTGSNVFQTR
Synthetic peptide
175.1204 z=?
259.1416 z=?
482.7491 z=?
551.2962 z=?
735.8852 z=2
865.4203 z=1
964.4917 z=1
1111.5526 z=?
1259.6060 z=1
1359.6765 z=1
m/z


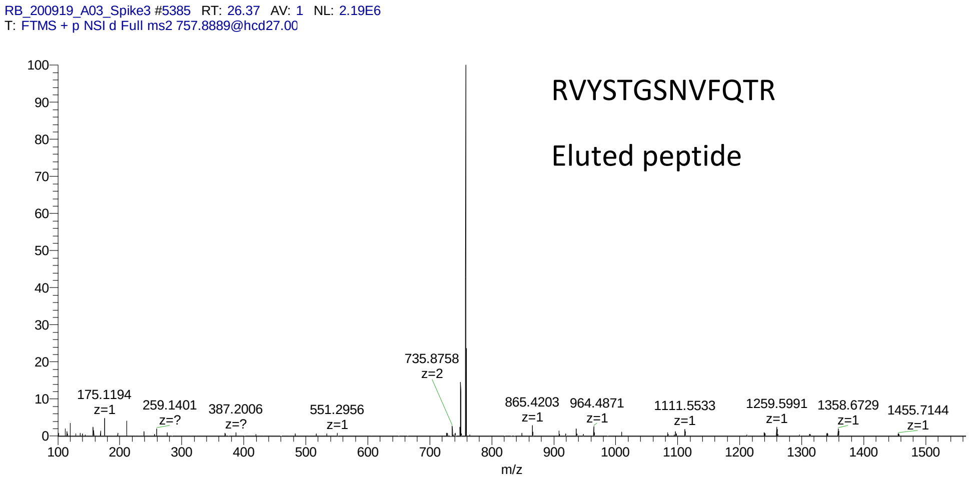

RB_200919_A03_Spike3 #5385 RT: 26.37 AV: 1 NL: 2.19E6
T: FTMS + p NSI d Full ms2 757.8889@hcd27.00
RVYSTGSNVFQTR
Eluted peptide
175.1194 z=1
259.1401 z=?
387.2006 z=?
551.2956 z=1
735.8758 z=2
865.4203 z=1
964.4871 z=1
1111.5533 z=1
1259.5991 z=1
1358.6729 z=1
1455.7144 z=1
m/z

RB_201016_PeptideMix_DDA #4969 RT: 26.83 AV: 1 NL: 4.13E4
T: FTMS + p NSI d Full ms2 679.8395@hcd27.00

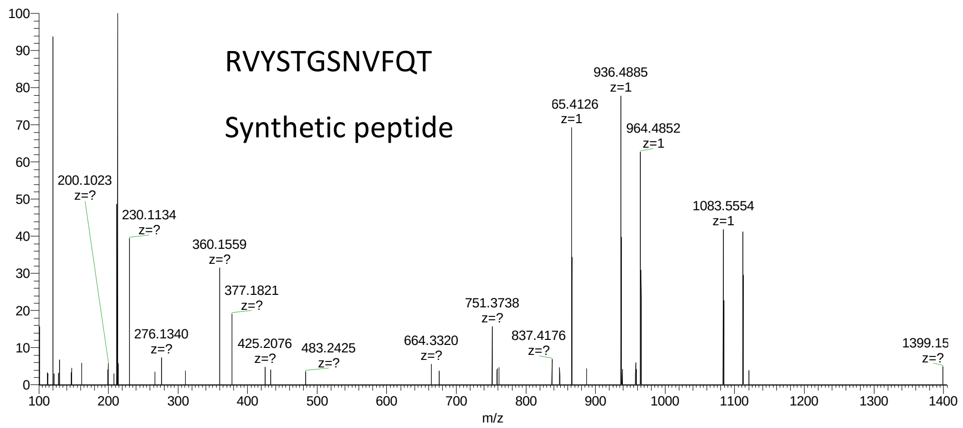


RB_200919_A03_Spike3 #7560 RT: 31.89 AV: 1 NL: 8.32E4
T: FTMS + p NSI d Full ms2 679.8375@hcd27.00

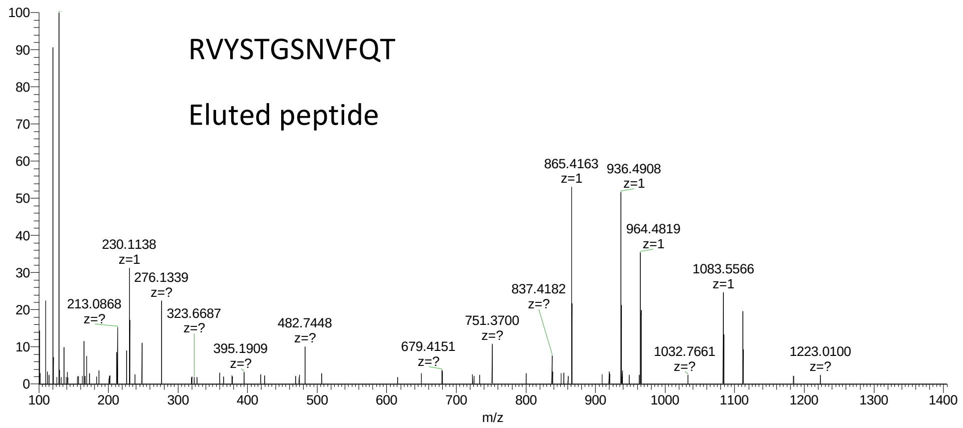

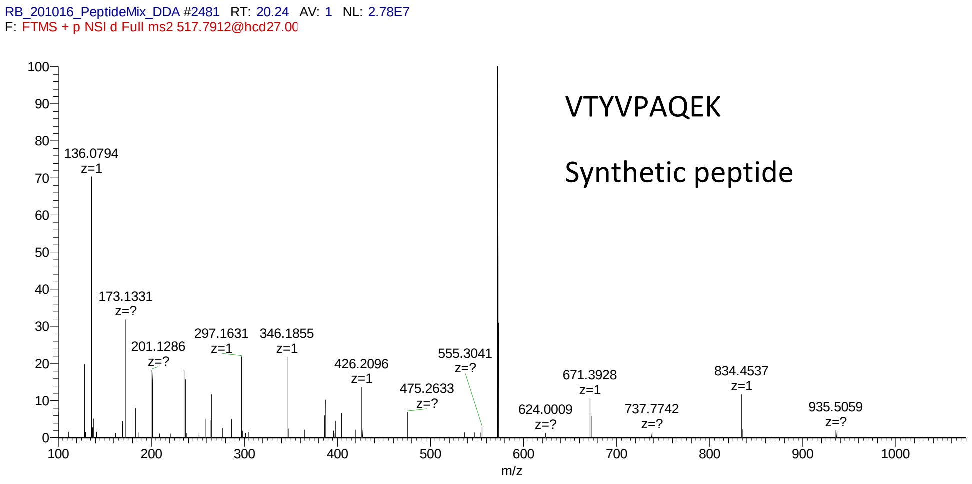
RB_201016_PeptideMix_DDA #2481 RT: 20.24 AV: 1 NL: 2.78E7
F: FTMS + p NSI d Full ms2 517.7912@hcd27.00
VTYVPAQEK
Synthetic peptide
136.0794 z=1
173.1331 z=?
201.1286 z=?
297.1631 z=1
346.1855 z=1
426.2096 z=1
475.2633 z=?
555.3041 z=?
624.0009 z=?
671.3928 z=1
737.7742 z=?
834.4537 z=1
935.5059 z=?
m/z

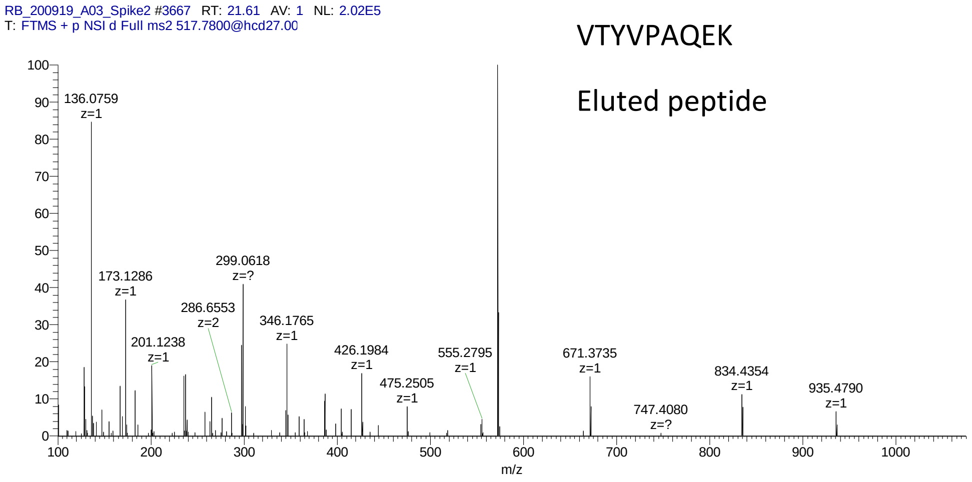
RB_200919_A03_Spike2 #3667 RT: 21.61 AV: 1 NL: 2.02E5
T: FTMS + p NSI d Full ms2 517.7800@hcd27.00
VTYVPAQEK
Eluted peptide
136.0759 z=1
173.1286 z=1
201.1238 z=1
286.6553 z=2
299.0618 z=?
346.1765 z=1
426.1984 z=1
475.2505 z=1
555.2795 z=1
671.3735 z=1
747.4080 z=?
834.4354 z=1
935.4790 z=1
m/z

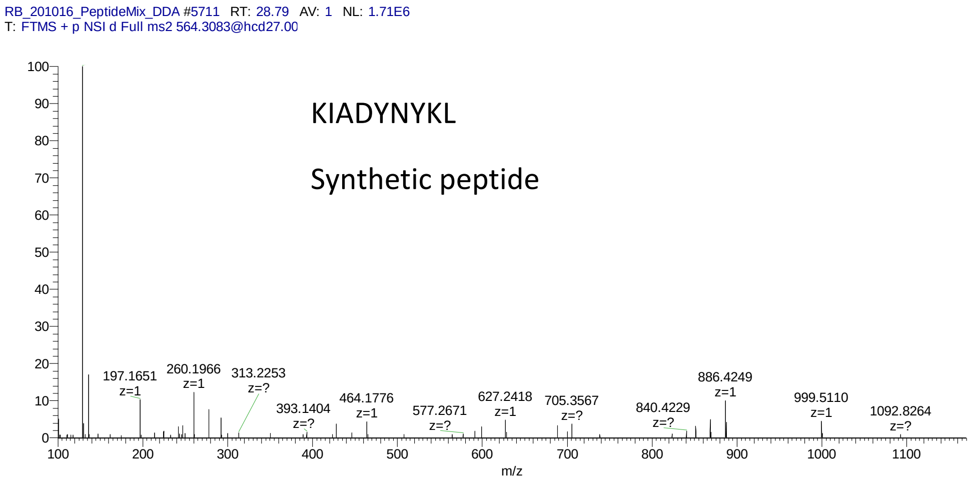
RB_201016_PeptideMix_DDA #5711 RT: 28.79 AV: 1 NL: 1.71E6
T: FTMS + p NSI d Full ms2 564.3083@hcd27.00
KIADYNYKL
Synthetic peptide
197.1651 z=1
260.1966 z=1
313.2253 z=?
393.1404 z=?
464.1776 z=1
577.2671 z=?
627.2418 z=1
705.3567 z=?
840.4229 z=?
886.4249 z=1
999.5110 z=1
1092.8264 z=?
m/z

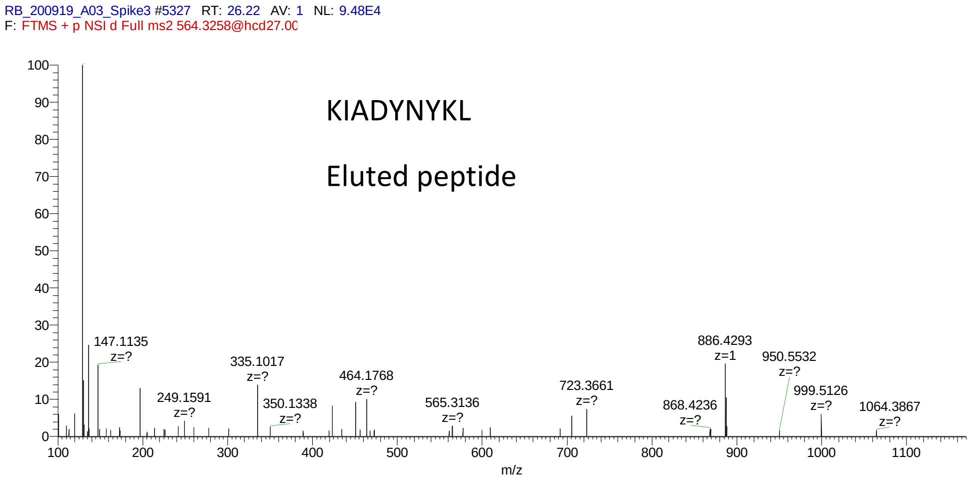
RB_200919_A03_Spike3 #5327 RT: 26.22 AV: 1 NL: 9.48E4
F: FTMS + p NSI d Full ms2 564.3258@hcd27.00
KIADYNYKL
Eluted peptide
147.1135 z=?
249.1591 z=?
335.1017 z=?
350.1338 z=?
464.1768 z=?
565.3136 z=?
723.3661 z=?
868.4236 z=?
886.4293 z=1
950.5532 z=?
999.5126 z=?
1064.3867 z=?
m/z

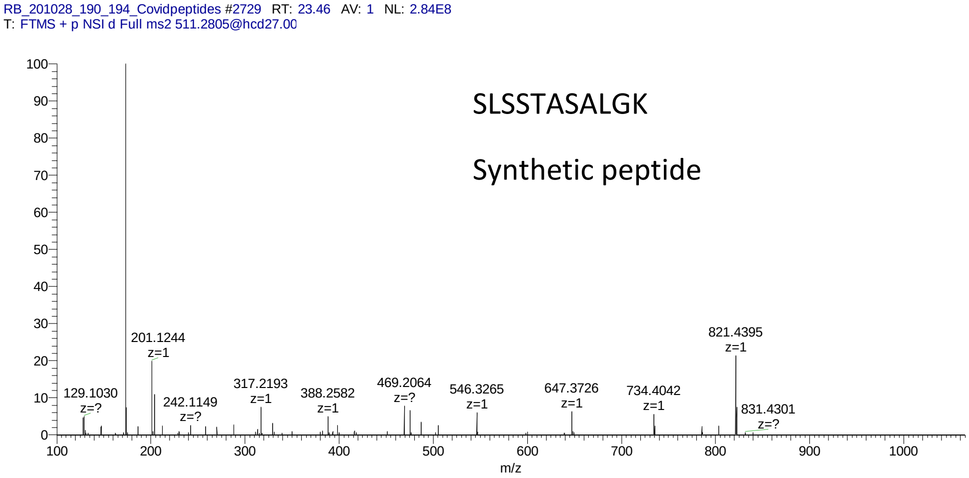
RB_201028_190_194_Covidpeptides #2729 RT: 23.46 AV: 1 NL: 2.84E8
T: FTMS + p NSI d Full ms2 511.2805@hcd27.00
SLSSTASALGK
Synthetic peptide
201.1244 z=1
129.1030 z=?
242.1149 z=?
317.2193 z=1
388.2582 z=1
469.2064 z=?
546.3265 z=1
647.3726 z=1
734.4042 z=1
821.4395 z=1
831.4301 z=?
m/z

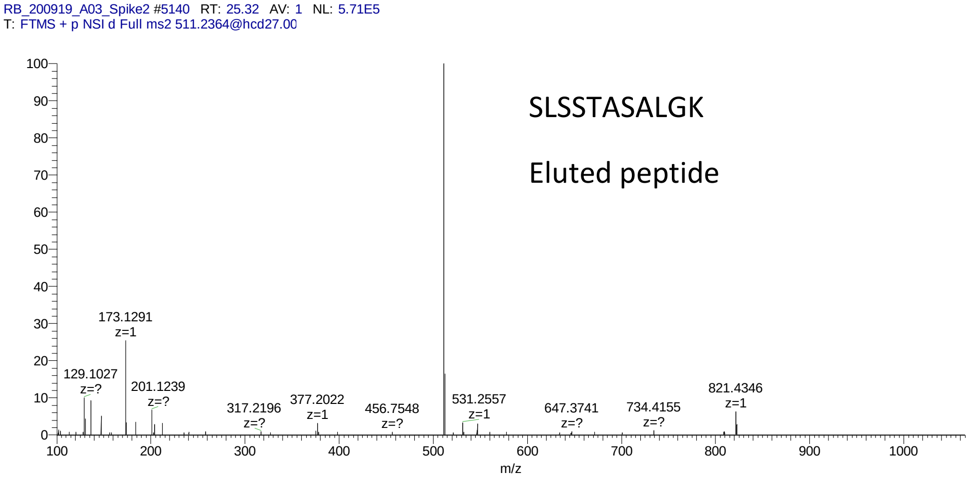
RB_200919_A03_Spike2 #5140 RT: 25.32 AV: 1 NL: 5.71E5
T: FTMS + p NSI d Full ms2 511.2364@hcd27.00
SLSSTASALGK
Eluted peptide
173.1291 z=1
129.1027 z=?
201.1239 z=?
317.2196 z=?
377.2022 z=1
456.7548 z=?
531.2557 z=1
647.3741 z=?
734.4155 z=?
821.4346 z=1
m/z

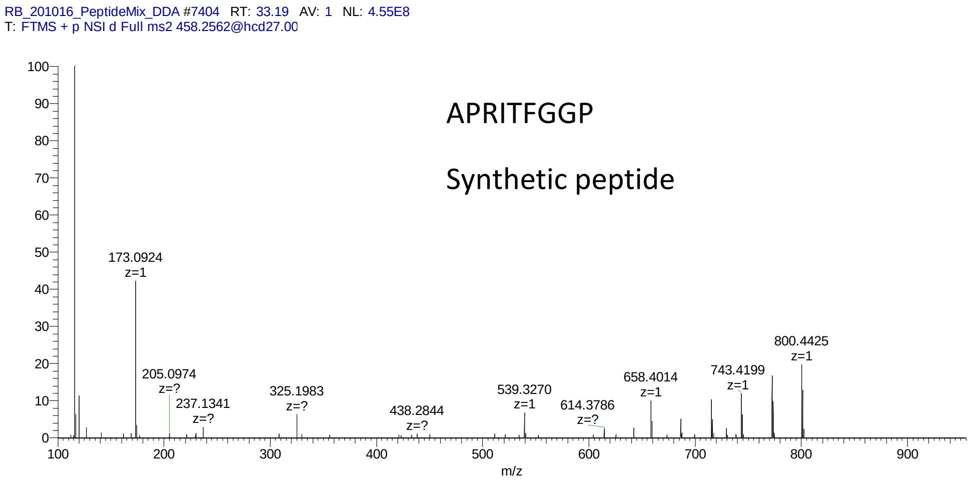
RB_201016_PeptideMix_DDA #7404 RT: 33.19 AV: 1 NL: 4.55E8
T: FTMS + p NSI d Full ms2 458.2562@hcd27.00
APRITFGGP
Synthetic peptide
173.0924 z=1
205.0974 z=?
237.1341 z=?
325.1983 z=?
438.2844 z=?
539.3270 z=1
614.3786 z=?
658.4014 z=1
743.4199 z=1
800.4425 z=1
m/z

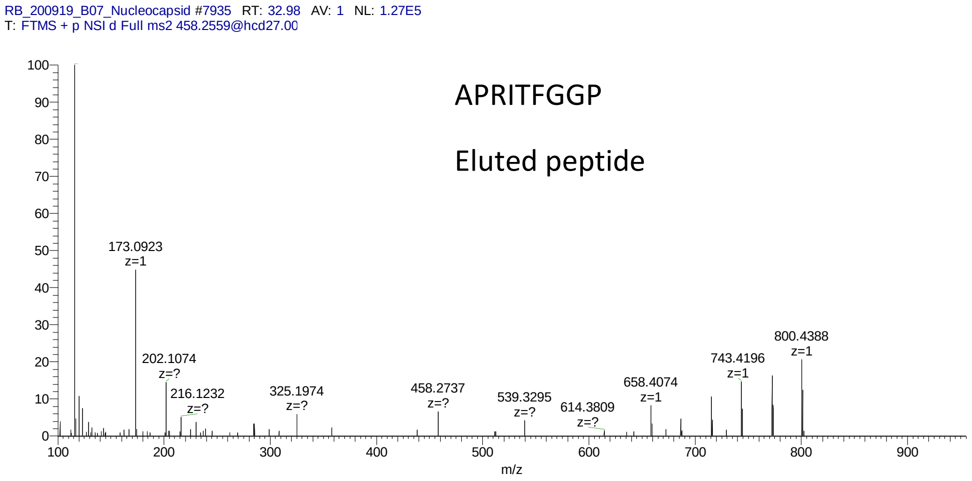
RB_200919_B07_Nucleocapsid #7935 RT: 32.98 AV: 1 NL: 1.27E5
T: FTMS + p NSI d Full ms2 458.2559@hcd27.00
APRITFGGP
Eluted peptide
173.0923 z=1
202.1074 z=?
216.1232 z=?
325.1974 z=?
458.2737 z=?
539.3295 z=?
614.3809 z=?
658.4074 z=1
743.4196 z=1
800.4388 z=1
m/z

RB_201016_PeptideMix_DDA #13936 RT: 50.27 AV: 1 NL: 4.30E5
T: FTMS + p NSI d Full ms2 575.8331@hcd27.00

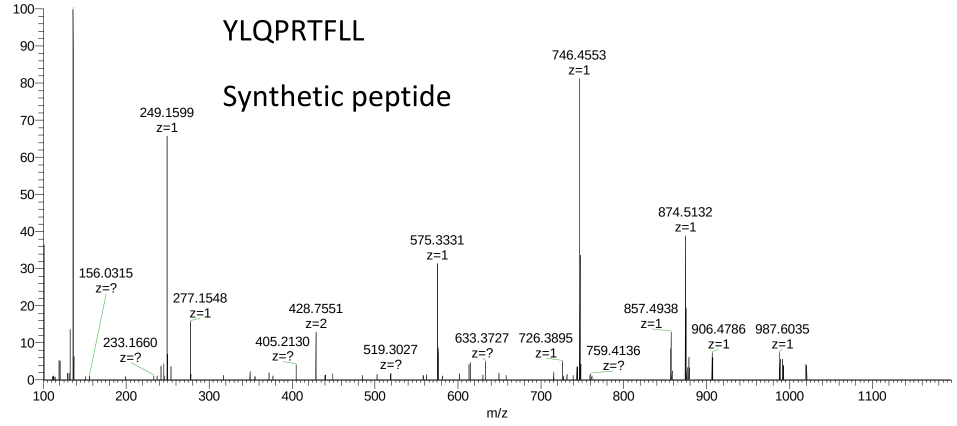


RB_200919_B08_Spike1 #13971 RT: 50.36 AV: 1 NL: 5.19E4
T: FTMS + p NSI d Full ms2 575.8345@hcd27.00

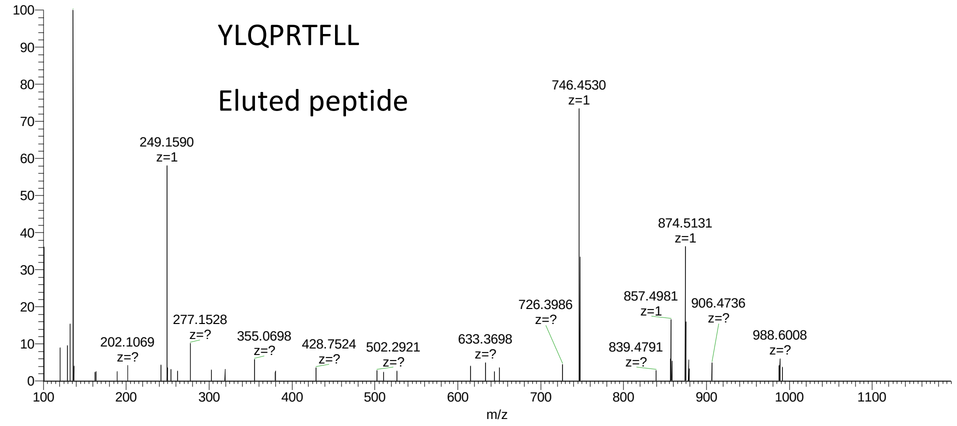

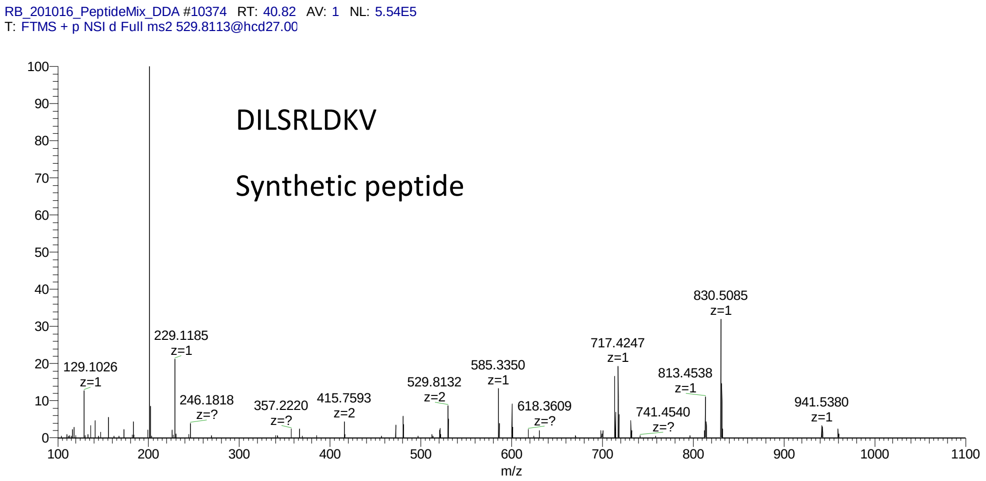
RB_201016_PeptideMix_DDA #10374 RT: 40.82 AV: 1 NL: 5.54E5
T: FTMS + p NSI d Full ms2 529.8113@hcd27.00
DILSRLDKV
Synthetic peptide
129.1026 z=1
229.1185 z=1
246.1818 z=?
357.2220 z=?
415.7593 z=2
529.8132 z=2
585.3350 z=1
618.3609 z=?
717.4247 z=1
741.4540 z=?
813.4538 z=1
830.5085 z=1
941.5380 z=1
m/z

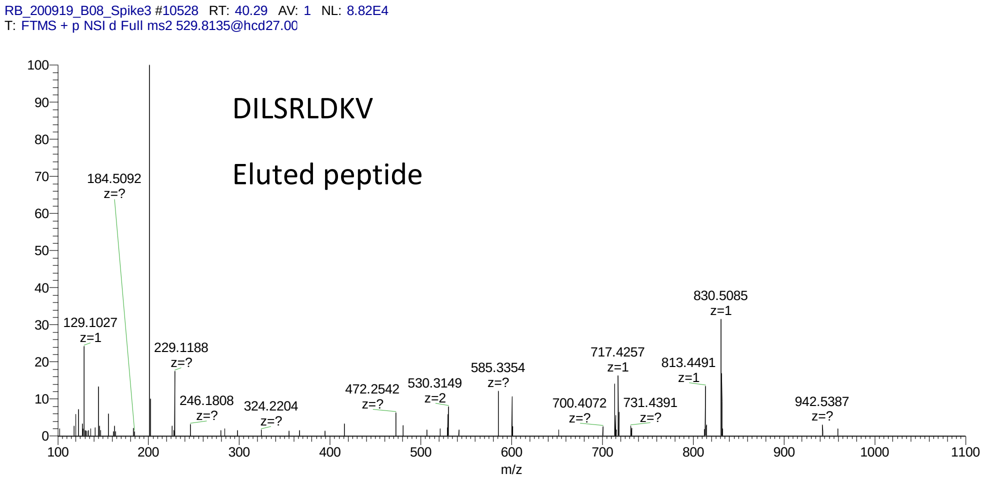
RB_200919_B08_Spike3 #10528 RT: 40.29 AV: 1 NL: 8.82E4
T: FTMS + p NSI d Full ms2 529.8135@hcd27.00
DILSRLDKV
Eluted peptide
184.5092 z=?
129.1027 z=1
229.1188 z=?
246.1808 z=?
324.2204 z=?
472.2542 z=?
530.3149 z=2
585.3354 z=?
700.4072 z=?
717.4257 z=1
731.4391 z=?
813.4491 z=1
830.5085 z=1
942.5387 z=?
m/z

RB_201016_PeptideMix_DDA #7919 RT: 34.51 AV: 1 NL: 1.07E6
T: FTMS + p NSI d Full ms2 521.3063@hcd27.00

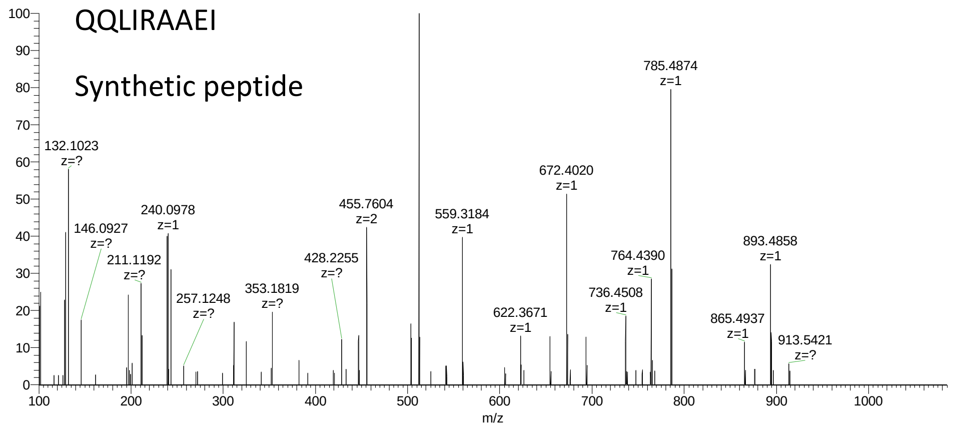


RB_200919_B08_Spike3 #8584 RT: 34.97 AV: 1 NL: 7.12E4
T: FTMS + p NSI d Full ms2 521.3058@hcd27.00

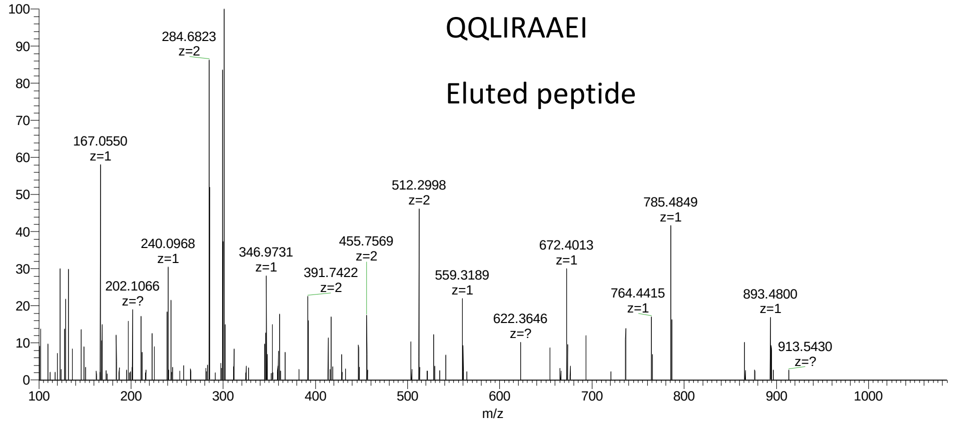

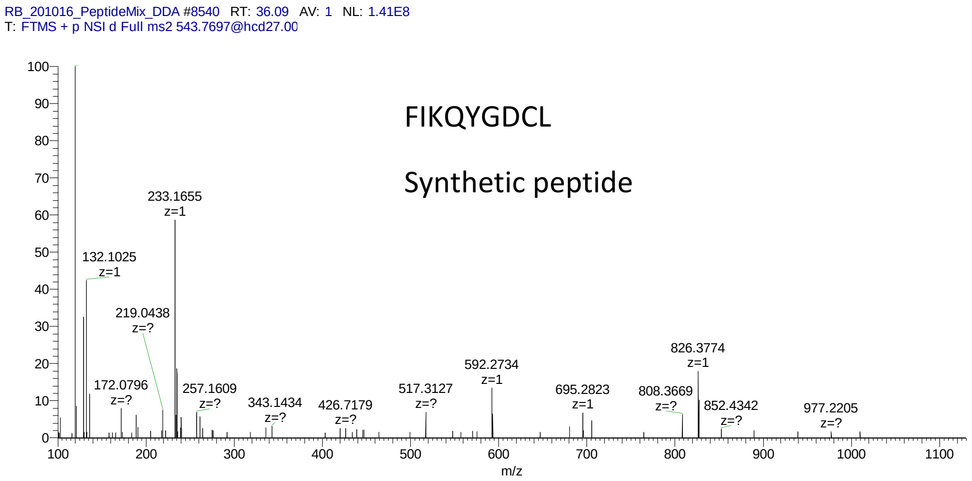
RB_201016_PeptideMix_DDA #8540 RT: 36.09 AV: 1 NL: 1.41E8
T: FTMS + p NSI d Full ms2 543.7697@hcd27.00
FIKQYGDCL
Synthetic peptide
233.1655 z=1
132.1025 z=1
219.0438 z=?
172.0796 z=?
257.1609 z=?
343.1434 z=?
426.7179 z=?
517.3127 z=?
592.2734 z=1
695.2823 z=1
808.3669 z=?
826.3774 z=1
852.4342 z=?
977.2205 z=?
m/z

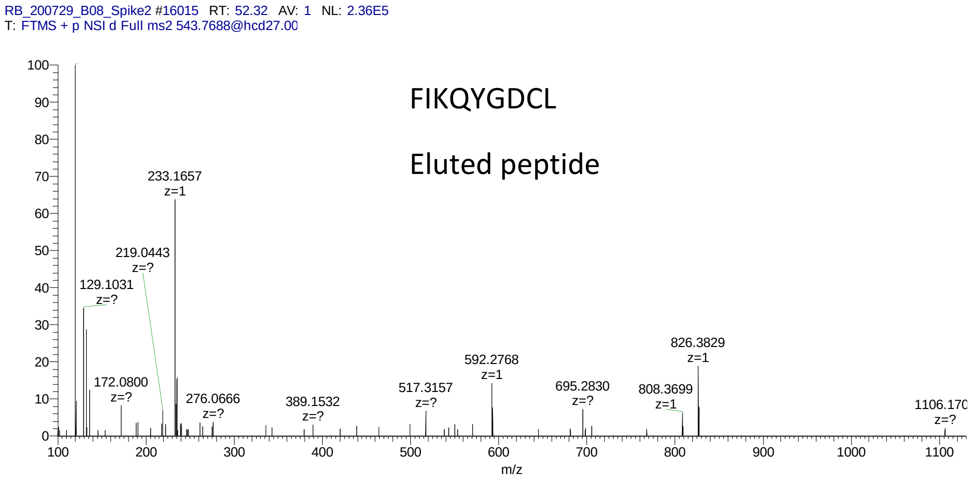
RB_200729_B08_Spike2 #16015 RT: 52.32 AV: 1 NL: 2.36E5
T: FTMS + p NSI d Full ms2 543.7688@hcd27.00
FIKQYGDCL
Eluted peptide
233.1657 z=1
219.0443 z=?
129.1031 z=?
172.0800 z=?
276.0666 z=?
389.1532 z=?
517.3157 z=?
592.2768 z=1
695.2830 z=?
808.3699 z=1
826.3829 z=1
1106.170 z=?
m/z
