## Supplementary material for "Beyond Spike: Identification of nine highly prevalent SARS-CoV-2-specific CD8 T-cell epitopes in a large Norwegian cohort": R code Population coverage

Adjusted population coverage


### Adjusted population coverage

###### EH Rustad & TS Madssen

### Introduction

Accompanying the manuscript: Public T cell epitopes shared among SARS-CoV-2 variants are presented on prevalent HLA class I alleles (Meyer, Blaas, Bollineni, et al.) As an essential tool in T cell vaccine design, several software tools have been developed to estimate the proportion of a population expected to mount an immune response to a set of epitopes, given the HLA restriction of the respective epitopes and frequencies of those HLA alleles in the population. Examples include the Immune Epitope Database (IEDB) population coverage tool (Bui et al, BMC Bioinformatics, 2005; http://tools.iedb.org/population/) and PopCover-2.0 (Nilsson JB et al, Front Immunol, 2021; https://services.healthtech.dtu.dk/service.php?PopCover-2.0).

Recent technological advances has made it possible to screen for immune responses to specific peptide/HLA combinations (pHLA) in multiple individuals at relatively high throughput. It has become clear that recognition of an epitope by CD8 T cells is not uniform even across individuals who express the relevant HLA allele(s). Indeed, epitopes vary extensively in the magnitude of immune responses they elicit in a given individual (i.e. immunodominance) as well as the number of individuals with a given HLA allele who will mount a response (i.e. immunoprevalence). Hence, immunoprevalence should be taken into account when designing vaccines for optimal population coverage; however, we could not find any such available software tool.

We set out to develop a simple algorithm which would that take into account immunoprevalence when estimating the population coverage of a set of epitopes. Our work started out by examining the existing IEDB population coverage software written in Python and implementing a similar algorithm in R. The IEDB population coverage tool calculates the proportion of a population with a given number of “hits” (i.e., number of epitopes recognized from a given set), and the population coverage is then the cumulative proportion with 1 or more hits. This concept relies on hits being binary, i.e. that a given epitope always (1) or never (0) elicits an immune response in individuals with a given HLA genotype. However, when immunoprevalence is introduced, immunogenicity is best understood as a probabiltity between 0 and 1.

Our algorithm is described in detail in the main text (Materials and Methods). In summary, for each HLA locus, we generate two sets of numbers to arrive at the population coverage for a set of epitopes. First, we list all possible combinations of HLA alleles in the population along with the proportion of individuals who have that combination. Next, for each of the HLA allele combinations, we take all the epitopes which may be presented on those alleles and calculate the probability that at least one of them elicits an immune response (i.e. recognition), treating the immunoprevalence of each epitope as the probability of recognition . We then multiplied the frequency of each HLA allele combination with the probability of having an immune response to at least one epitope presented on the respective alleles, taking the sum of all the products to achieve a population coverage adjusted for immunoprevalence. We repeated this procedure independently for each HLA locus and then combined in the end (i.e. the probability of having an immune response from at least one locus).

The true immunoprevalence of an epitope in the general population (with the relevant HLA allele) is often uncertain because a limited number of individuals have been evaluated for immunogenicity. We wanted to evaluate how this uncertainty affected the final estimates of population coverage through generating 95 % confidence intervals. We achieved this by a parameteric bootstrapping approach, where the immunoprevalence for each epitope-allele pair was treated as a random variable generated from a beta distribution, with shape parameters determined by the experimental immunoprevalence data. For each of these new bootstrapped immunoprevalence values, the procedure as described above was repeated, and the 2.5th and 97.5th percentiles formed the lower and upper bound for the 95 % confidence interval. Our bootstrapping approach treats immunoprevalence as the result of a binomial process and recapitulates how exact binomial 95 % confidence intervals are estimated in the binom.test R function.

### Functions for population coverage estimation

```
# load packages
packages <- c("tidyverse", "reshape2", "openxlsx", "RColorBrewer",  "stringi", "gdata")

invisible(suppressWarnings(suppressMessages(lapply(packages, library, character.only = TRUE))))

# custom functions
rotatedAxisElementText = function(angle,position='x'){
    angle     = angle[1]; 
    position  = position[1]
    positions = list(x=0,y=90,top=180,right=270)
    if(!position %in% names(positions))
        stop(sprintf("'position' must be one of [%s]",paste(names(positions),collapse=", ")),call.=FALSE)
    if(!is.numeric(angle))
        stop("'angle' must be numeric",call.=FALSE)
    rads  = (angle - positions[[ position ]])*pi/180
    hjust = 0.5*(1 - sin(rads))
    vjust = 0.5*(1 + cos(rads))
    element_text(angle=angle,vjust=vjust,hjust=hjust)
}

# calculate the probability of one or more responses to epitopes presented on a given HLA allele combination
# takes as input a vector of two HLA alleles (allele_set) and a data frame of immunoprevalence for the included set of epitopes

get_allele_comb_prob <- function(allele_set, immunoprev){
  e_set <- immunoprev[immunoprev$HLA %in% allele_set,]$prop
  
  # if only a single epitope from a given combination, then the probability of one or more responses is equal to the probability of one response
  if(length(e_set) == 1) { 
    return(e_set)
  } 
  
  no_response_prob <- 1-e_set
  
  product <- no_response_prob[1]
  for(i in 2:length(no_response_prob)){
    product <- product*no_response_prob[i]
  }
  
  out <- ifelse(length(1-product) == 0, 0, 1-product)
  return(out)
}

# calculate the population prevalence of each combination of HLA alleles, including "other" HLA-alleles not present in the input.
# takes as input a vector of HLA alleles to include (allele_vector) and a file with allele prevalence data (allele_freq_df)

get_allele_comb_prevalence <- function(allele_vector, allele_freq_df){
  hla_sub <- filter(allele_freq_df, Allele %in% allele_vector)
  freq_vector <- hla_sub$freq
  names(freq_vector) <- hla_sub$Allele
  freq_vector["UNKNOWN"] <- 1-sum(freq_vector)
  
  freq_mat <- freq_vector %*% t(freq_vector)
  rownames(freq_mat) <- colnames(freq_mat)
  
  freq_mat_vector <- unmatrix(freq_mat)
  
  return(freq_mat_vector)
}

# takes a vector of HLA alleles, returns a list of every possible combination of two HLA alleles from the input vector

make_allele_comb_list <- function(allele_vector){
  allele_comb_list <- list()
  for(i in allele_vector){
    for(j in allele_vector)
      allele_comb_list[[paste(i,j, sep = ":")]] <- c(i,j)
  }
  return(allele_comb_list)
} 

# LOCUS SPECIFIC FUNCTION

get_locus_coverage <- function(locus, locus_epitope_df, allele_freq_df) {
  
  locus_epitope_set <- locus_epitope_df %>% filter(grepl(locus, HLA))
  allele_vector <- unique(locus_epitope_set$HLA)

  # allele combination prevalence
  freq_mat_vector <- get_allele_comb_prevalence(allele_vector = allele_vector,
                                                allele_freq_df = allele_freq_df)
  
  # all allele combinations, including UNKNOWN
  allele_comb_list <- make_allele_comb_list(allele_vector = c(allele_vector, "UNKNOWN"))

  # probability of one or more epitope response for allele combinations
  allele_comb_probs <- unlist(lapply(allele_comb_list, function(x) get_allele_comb_prob(allele_set = x, immunoprev = locus_epitope_set)))

  # allele combination prevalence adjusted by probability of one or more epitope response
  freq_mat_vector <- freq_mat_vector[names(allele_comb_probs)]
  prev_adjusted_freq <- freq_mat_vector * allele_comb_probs

  # locus response population coverage
  locus_coverage <- sum(prev_adjusted_freq)
  
  return(locus_coverage)
}

# bootstrapping sets of immunoprevalence estimates from the beta distribution
# using the same approach as employed to calculate exact binomial confidence intervals (binom.test)

get_bootstrap_dataset <- function(epi_data, n_bootstrap){
  epi_boot <- list()
  
  for(i in 1:nrow(epi_data)){
    n_success <- epi_data$N_response[i]
    n_failure <- epi_data$N_tested[i] - n_success + 1
    boot_prop <- rbeta(n = n_bootstrap, shape1 = n_success, shape2 = n_failure)
    
    tmp_out <- data.frame(Sequence = rep(epi_data$Sequence[i], n_bootstrap),
                          HLA = rep(epi_data$HLA[i], n_bootstrap),
                          epitope_ID = rep(epi_data$epitope_ID[i], n_bootstrap),
                          iter = 1:n_bootstrap,
                          prop = boot_prop)
    epi_boot[[i]] <- tmp_out
  }
  
  epi_boot <- bind_rows(epi_boot)
  
  return(epi_boot)
}

# run all analysis and combine loci -- returning total population coverage

get_combined_coverage <- function(epitopes, allele_freq){
  # HLA A coverage
  coverage_A <- get_locus_coverage(locus = "HLA-A", locus_epitope_df = epitopes, allele_freq_df = allele_freq)
  
  # HLA B coverage
  coverage_B <- get_locus_coverage(locus = "HLA-B", locus_epitope_df = epitopes, allele_freq_df = allele_freq)

  # COMBINED

  # probability of no hits on both HLA A and B
  prob_no_hits <- (1-coverage_A) * (1-coverage_B)
  
  # probability of at least one hit
  coverage_combined <- 1-prob_no_hits
  
  return(coverage_combined)
  
}

get_population_coverage <- function(epitopes, allele_freq, bootstrap = TRUE, n_bootstrap = 100, conf.level = 0.95){
  out <- list()
  
  # point estimate
  out$est <- get_combined_coverage(epitopes = epitopes, allele_freq = allele_freq)

  # bootstrapping confidence interval
  if(bootstrap){
    
    epi_boot_dataset <- get_bootstrap_dataset(epitopes, n_bootstrap)
  
    pop_cov_boot <- list()
    for(i in 1:max(epi_boot_dataset$iter)){
      epi_boot_sub <- filter(epi_boot_dataset, iter == i)
      pop_cov_boot[[i]] <- get_combined_coverage(epitopes = epi_boot_sub, allele_freq = allele_freq)
    }
    pop_cov_boot <- unlist(pop_cov_boot)

    # confidence interval
    alpha <- (1 - conf.level) / 2
    out$conf.int <- quantile(probs = c(alpha, 1-alpha), pop_cov_boot)
      
    # add raw bootstrap values
    out$boot_data <- epi_boot_dataset
    out$boot <- unlist(pop_cov_boot)
  }
  out$data <- epitopes
  return(out)
}
```

### Data

Loading immunoprevalence data for the top 9 epitopes identified in our study.

```
epi_sub <- read.xlsx("./data/top_epitopes_this_study.xlsx")
epi_sub
```

```
##     Sequence         HLA prop N_tested N_response    epitope_ID
## 1  FTSDYYQLY HLA-A*01:01 1.00       20         20 A1 O3a207-215
## 2 LHSYFTSDYY HLA-A*01:01 0.86       14         12 A1 O3a203-212
## 3 YFTSDYYQLY HLA-A*01:01 1.00       20         20 A1 O3a206-215
## 4  LLYDANYFL HLA-A*02:01 0.91       53         48 A1 O3a139-147
## 5  YLQPRTFLL HLA-A*02:01 0.98       53         52   A1 S269-277
## 6  KTFPPTEPK HLA-A*03:01 0.90       29         26   A1 N366-374
## 7 KCYGVSPTKL HLA-A*03:01 0.83       29         24   A1 S378-387
## 8  KPRQKRTAT HLA-B*07:02 0.70       20         14   A1 N257-265
## 9  SPRWYFYYL HLA-B*07:02 1.00       32         32   A1 N105-113
```

Loading immunoprevalence data from IEDB, selecting only those epitopes studied in at least 2 studies or 20 individuals. Among those epitopes with sufficient supporting data, we included all immunogenic epitopes irrespective of immunoprevalence.

```
iedb <-  read.xlsx("./data/epitopes_iedb.xlsx")
head(iedb)
```

```
##      Sequence         HLA   prop N_tested N_response     epitope_ID N_studies
## 1  TTDPSFLGRY HLA-A*01:01 0.9375       32         30  A2 TTDPSFLGRY         5
## 2   SPRWYFYYL HLA-B*07:02 0.8571       42         36   A2 SPRWYFYYL         6
## 3   ILFTRFFYV HLA-A*02:01 0.8333       18         15   A2 ILFTRFFYV         3
## 4   YLQPRTFLL HLA-A*02:01 0.7871      263        207   A2 YLQPRTFLL         9
## 5 HTTDPSFLGRY HLA-A*01:01 0.7500       16         12 A2 HTTDPSFLGRY         3
## 6 TTDPSFLGRYM HLA-A*01:01 0.7143        7          5 A2 TTDPSFLGRYM         2
```

```
# number of epitopes per HLA allele in IEDB after filtering
table(iedb$HLA)
```

```
## 
## HLA-A*01:01 HLA-A*02:01 HLA-A*03:01 HLA-B*07:02 
##          19          41          18          15
```

To estimate the effect of immunoprevalence on population coverage, we generated 1000 random sets of epitopes by drawing from IEDB. Each set consisted of 9 unique epitopes with the same HLA distribution as our top 9 epitopes.

```
# randomly generate IEDB subsets

sub_hla_counts <- as.data.frame(table(epi_sub$HLA), stringsAsFactors = F)
names(sub_hla_counts) <- c("HLA", "count")

iedb_random <- list()
n_subsets <- 1000
for(i in 1:n_subsets){
  tmp_df <- data.frame(matrix(ncol = ncol(iedb), nrow = 0))
  names(tmp_df) <- names(iedb)
  
  for(j in 1:nrow(sub_hla_counts)){
    n_sub <- sub_hla_counts$count[j]
    hla_sub <- sub_hla_counts$HLA[j]
    
    random_subset <- iedb %>%
      filter(HLA == hla_sub) %>%
      sample_n(n_sub)
    
    tmp_df <- rbind(tmp_df,
                    random_subset)
    
  }
  iedb_random[[i]] <- tmp_df
}
```

Allele frequencies in europeans for relevant HLA alleles were obtained from http://www.allelefrequencies.net (Gonzalez-Galarza et al, Nucleic Acid Research 2020) and adjusted so that frequencies for each locus sum to 1.

```
allele_freq_df <- read.xlsx("./data/european_alleles_freq.xlsx")
allele_freq_df
```

```
##   Locus      Allele    freq
## 1     A HLA-A*01:01 0.13784
## 2     A HLA-A*02:01 0.27202
## 3     A HLA-A*03:01 0.13427
## 4     B HLA-B*07:02 0.11373
## 5     B HLA-B*08:01 0.09301
```

### Run analysis for our data

Population coverage of epitope set after adjusting for immunoprevalence.

```
set.seed(1)

pop_coverage <- get_population_coverage(epitopes = epi_sub, allele_freq = allele_freq_df, bootstrap = TRUE, n_bootstrap = 1000, conf.level = 0.95)


pop_coverage$est
```

```
## [1] 0.8344
```

```
pop_coverage$conf.int
```

```
##   2.5%  97.5% 
## 0.8279 0.8353
```

For comparison, we show the unadjusted HLA population coverage calculated using the IEDB population coverage software.

```
unadjusted_coverage <- 1-0.16323635

unadjusted_coverage
```

```
## [1] 0.8368
```

The loss of coverage by incorporating immunoprevalence was minimal for our epitopes.

```
unadjusted_coverage - pop_coverage$est
```

```
## [1] 0.002332
```

### bootstrapping confidence interval

Figure below identical to Supplementary Figure 7A. Uncertainty in immunoprevalence estimates for our top 9 immunoprevalent epitopes based on bootstrapping from the beta distribution. Each of 1000 iterations per epitope are represented as shaded gray dots. Boxplots show median and inter-quartile range (IQR); whiskers extend to +/- 1.5 times the IQR. The observed immunoprevalence in experimental data is drawn as a red dot.

```
ggplot()+
  geom_jitter(data = pop_coverage$boot_data, aes(epitope_ID, prop), col = "darkgray", height = 0, width = 0.4, alpha = 0.2)+
  geom_boxplot(data = pop_coverage$boot_data, aes(epitope_ID, prop), fill = NA, outlier.shape = NA)+
  geom_point(data = pop_coverage$data, aes(epitope_ID, prop), col = "firebrick2", size = 2)+
  theme_bw()+
  theme(axis.text.x = rotatedAxisElementText(90, "top"),
        text = element_text(color = "black"))+
  scale_y_continuous(limits = c(0,1))+
  labs(x = "Epitope",
       y = "Immunoprevalence\n(bootstrap estimates)")
```

Figure below identical to Supplementary Figure 7B. Coverage of European population with immunoprevalent peptides identified in this study. The dashed vertical blue line represents population coverage with peptides recognizing HLA-A*01:01, HLA-A*02:01, HLA-A*03:01 and HLA-B*07:02, based solely on HLA allele frequencies (not considering immunoprevalence). Estimated population coverage with our top 9 epitopes (prevalence 0.7 or higher) after adjusting for immunoprevalence is shown as a dashed vertical red line. Histogram in gray shows population coverage for the same 9 epitopes when considering uncertainty in immunoprevalence estimates, as represented by bootstrapped data shown in (7A).

```
hist(pop_coverage$boot, main = "", xlab = "Estimated population coverage (proportion)", ylab = "Simulations (n)", xlim = c(0.82, 0.84))
abline(v = pop_coverage$est, col = "firebrick2", lty = 2, lwd = 3)
abline(v = unadjusted_coverage, col = "darkblue", lty = 2, lwd = 3)
```

### random epitopes from IEDB

```
iedb_random_prev <- lapply(iedb_random, function(x) {
  get_population_coverage(epitopes = x, allele_freq = allele_freq_df, bootstrap = FALSE, n_bootstrap = 1000, conf.level = 0.95)
})

iedb_random_est <- sapply(iedb_random_prev, function(x) x$est)

summary(iedb_random_est)
```

```
##    Min. 1st Qu.  Median    Mean 3rd Qu.    Max. 
##   0.191   0.389   0.456   0.467   0.540   0.753
```

Figure below not included in the paper. Coverage of European population with 1000 sets of 9 random epitopes from IEDB, maintaining the same HLA allele distribution as in our epitopes. The dashed vertical blue line represents population coverage with peptides recognizing HLA-A*01:01, HLA-A*02:01, HLA-A*03:01 and HLA-B*07:02, based solely on HLA allele frequencies (not considering immunoprevalence). Mean population coverage from the 1000 random epitope sets after adjusting for immunoprevalence is shown as a dashed vertical red line.

```
hist(iedb_random_est, main = "", xlab = "Estimated population coverage (proportion)", ylab = "Random epitope sets from IEDB (n)", xlim = c(0.15, 0.84))
abline(v = mean(iedb_random_est), col = "firebrick2", lty = 2, lwd = 3)
abline(v = unadjusted_coverage, col = "darkblue", lty = 2, lwd = 3)
```

### Impact of immunoprevalence on population coverage

```
comb_data_toplot <- rbind(data.frame(coverage = iedb_random_est,
                                     source = "Random IEDB"),
                          data.frame(coverage = pop_coverage$boot,
                                     source = "This study")) %>%
  mutate(source = factor(source, levels = c("This study", "Random IEDB")))
```

Figure below identical to Figure 4E Coverage of European population with immunoprevalent peptides identified in this study as compared with 1000 random sets of immunogenic peptides from IEDB. The dashed black line represents population coverage with peptides recognizing HLA-A*01:01, HLA-A*02:01, HLA-A*A03:01 and HLA-B*07:02, based solely on HLA allele frequencies (not considering immunoprevalence). Red box plot represents estimated population coverage with our top 9 epitopes (immunoprevalence 70% or higher) after adjusting for immunoprevalence, bootstrapping from the beta distribution of each epitope prevalence to quantify uncertainty (1000 iterations). In blue are results from 1000 random draws of immunogenic epitopes from IEDB, while keeping the HLA allele distribution the same as in our top epitopes (3 HLA-A*01:01peptides, 2 HLA-A*02:01 peptides, 2 HLA-A*03:01 peptides and 2 HLA-B*07:02 peptides). Boxplots show median and inter-quartile range (IQR); whiskers extend to +/- 1.5 times the IQR and more extreme values are drawn as dots.

```
ggplot(comb_data_toplot)+
  geom_hline(yintercept = unadjusted_coverage, col = "black", size = 1, linetype = 2)+
  geom_boxplot(aes(source, coverage, col = source), fill = NA)+
  scale_color_brewer(palette = "Set1")+
  scale_y_continuous(limits = c(0, 1))+
  labs(y = "Estimated population coverage")+
  theme_bw()+
  theme(text = element_text(colour = "black"),
        axis.title.x = element_blank(),
        legend.position = "none")
```

These data demonstrate that immunoprevalence has a striking impact on the population coverage of an epitope set even when HLA restriction is held constant.
